# Large-scale prediction of transcription factor binding across human cell types informs regulatory genomics and reveals promiscuous occupancy associated with chromatin contacts

**DOI:** 10.64898/2026.06.16.732527

**Authors:** Emanuel Sonder, İpek Güneş Aymergen, Jieran Sun, Audran Feuvrier, Gerhard Schratt, Katharina Gapp, Johannes Bohacek, Mark D. Robinson, Pierre-Luc Germain

## Abstract

Our understanding of the mechanisms regulating gene expression has been hampered by our limited knowledge of which transcription factors (TFs) bind where in the genome, which is highly cell type-specific. While genome-wide TF binding can be experimentally assayed for individual TFs in individual cell types, profiling the full combinations of over 1600 TFs in hundreds of cell types is beyond practical reach. In this work, we developed a streamlined platform, *TFBlearner*, to train TF-specific models and predict bindings based on ATAC-seq data. We focused on biologically-motivated feature engineering and harnessed TF cooperativity and binding similarity across cell types to achieve state of the art binding predictions in unseen cell types in a scalable fashion. This enabled us to generate a compendium of binding predictions for 1108 Chromatin-associated proteins, of which 960 TFs, across 43 human cell types including widely-used cell lines and 36 physiological cell types representing all major human cell lineages. We show how the models additionally provide biological insights on the TFs, and show how the binding predictions can be used in downstream tasks such as TF activity inference. Our study additionally led to the observation of high promiscuity in TF occupancy. To investigate aspecific occupancy, we characterized crowded or high-occupancy (HOT) regions across cell types, providing evidence of their functionality, and reporting important cell type-specificity. Finally, we show that, across cell types, crowded regions engage in more 3D contacts, and that most TF occupancy at crowded promoters can be explained as tethered bindings from distal regulatory elements.

## Introduction

### Reconstruction of gene regulatory mechanisms

Central to much of biology is the regulation of gene expression, especially achieved through combinatorial interactions of transcription factors (TFs) and cofactors with the genome. The elucidation of these interactions is critical to a mechanistic understanding of gene expression, and while this can be readily achieved for specific loci or regulatory systems, there has been a long-standing interest into unbiased, systems-level computational approaches to the discovery of such mechanisms.

On the one hand, there is a long tradition of reverse-engineering gene regulatory networks based on coexpression analysis [1–6]. Such coexpression networks are composed of so-called ‘influence interactions’, which are undirected associations between genes that are most often not directly causal. As such, the more recent foundation models based on very large single-cell RNAseq datasets represent a powerful successor to such general approaches [7–9]. While generally promising, such methods have had limited success at predicting the expression outcomes of perturbations, and are often outperformed by simpler methods [10– 12]. While this is likely to improve with more interventional data (e.g. [13]), serious concerns have been raised about the capacity of such models to distinguish causation from correlation in complex molecular networks, and a convincing case has been made for the importance of prior (ideally causal) knowledge also in such endeavors [14, 15]. Indeed, the archetypical task of such models – predicting gene expression in response to perturbation – does not per se require distinguishing causal interactions from associations. As such, while this family of approaches is powerful for prediction and to identify regulatory modules, their capacity to yield mechanistic understanding is questionable [14, 15].

Another family of approaches explicitly relies on putative TF targets to reconstruct transcriptional networks or estimate TF activity, either from DNA accessibility [16, 17], the transcriptome [18, 19], or their combination. In particular, recent methods have harnessed the wealth of multi-modal single-cell sequencing data to reconstruct transcription networks [20–24]. A critical component of such approaches is where on the genome TFs bind to, which is highly context-specific. While genome-wide TF binding can be experimentally assayed for individual TFs in individual cell types, the underlying techniques are time-consuming and not easily multiplexed, and thus experimentally profiling the full combinations of roughly 1600 human TFs [25] in hundreds of cell types (let alone in treatment or disease conditions) is beyond practical reach. Indeed, while genome-wide binding data is available for over a thousand human TFs, the vast majority have been profiled in only one or two cell types. For this reason, researchers typically rely on proxies for TF binding, the most common of which is the presence of binding motifs, i.e. DNA sequence patterns with some degree of specificity for a given TF. However, because a myriad of factors other than the TF’s affinity for a sequence determines its binding, such motifs have very limited power in predicting in vivo binding.

### TF binding specificity

TFs are traditionally characterized by their binding to specific DNA sequences, however the reality is more complex: while it is commonly acknowledged that the vast majority of motif matches in the genome are not bound in a given cell type, it is more seldom appreciated that motifs are present in only a fraction of the sites showing occupancy of the respective TF (see Results). While this may be partly due to incomplete motif lexicons, another, critical factor is the interplay with the local chromatin environment and in particular with other TFs and cofactors [26].

Most TFs show a certain degree of promiscuity, or relatively low specificity. A particularly extreme example are ‘crowded’ or ‘highly-occupied targets’ (HOT) regions, i.e. regions showing high occupancy (as measured by ChIP-seq) for very many TFs (the exact definition varying). Such regions were characterized early on across multiple species [27–29], although it has been debated whether they represent (at least in part) artifacts [30–34]. Across species, these regions are enriched for promoters, and have been associated with loosely-defined ‘housekeeping’ (or virtually ubiquitously-expressed) genes [28, 33, 35] and RNA processing genes [28, 29, 34]. Importantly, these regions have been described as ‘motif-less’, or rather harboring a clear motif for only a subset of the TFs described to bind them [27, 28, 32]. A plausible hypothesis, therefore, is that bindings at these regions are dependent on protein-protein interactions [26, 36].

### Predicting cell type-specific TF binding

The prediction of TF binding to the genome is an well-established area of research. In particular, predicting cell type-specific TF binding from accessibility data has been a recognized challenge in the field, with several methods proposed, initially using DNase-seq [37–40]. More recently, ATAC-seq has received widespread adoption, in particular due to the low input material required, making it applicable to small in vivo samples and even single cells. Setting aside approaches that require multimodal profiling [41, 42], to our knowledge the best method to date is maxATAC [43], whose authors also performed the most systematic prediction effort to date, having trained deep learning models for the binding of 127 TFs. While the effort is impressive, this represents a mere 8% of human TFs, and the approach does not scale effectively for making predictions across hundreds of cell types.

In this work, we developed a prediction strategy that scales to thousands of TFs in hundreds of cell types. To this end, we favored extensive, biologically-motivated feature engineering. In particular, we sought to 1) use the rich information provided by ATAC-seq fragment sizes and insertion patterns, as well as patterns of association across cell types, and 2) harness TF cooperativity and binding similarity across cell types by using ChIP-seq data from other cell types and TFs as part of the predictions. We implemented a prediction pipeline, termed TFBlearner, with a focus on improving generalizability across cell types, and applied it to the prediction of 1085 TFs in 43 human cell types, with a special emphasis on brain cell types. We characterize these models, and show how the predictions can be used in downstream tasks such as TF activity inference.

## 1 Results

### 1.1 A consensus set of Putative Regulatory Elements (PREs)

Handling genome-wide data for thousands of factors across cellular contexts is computationally highly expensive. To streamline the process, we sought to restrict the space of hypotheses to a pre-defined set of regions, or Putative Regulatory Elements (PREs). To this end, we merged sets of DNaseI hyper-sensitive sites (DHS) with cell type-specific nucleosome-free regions, and used a merging-and-splitting procedure (see Methods) to obtain a set of 3.8 million non-overlapping PREs with a mean width 205 bp and 99% of the regions between 90 and 574 bp (Figure 1A). We confirmed that using these windows (instead of the whole genome) incurred minimal loss by looking at the proportion of ENCODE ChIP-seq peaks overlapping PREs (Fig. 1B). 99.8% of all ENCODE peaks overlapped PREs, and the median proportion of a TF’s peaks overlapping PREs was 99%. Some ChIP experiments, however, showed a lower overlap with PREs (Fig. 1B), despite being in well-characterized cell lines. These chiefly came from a small set of TFs including YBX1 (65%), HMGA2 (66%), MAFF/MAFK (75%), and several KRAB-containing zinc fingers (e.g. ZNF354C, ZNF8, ZNF324 with 55-64%). As the latter are involved in the silencing of transposable elements, it is expected that a substantial fraction of their binding will not overlap regions defined by their accessibility pattern.

**Fig. 1:**
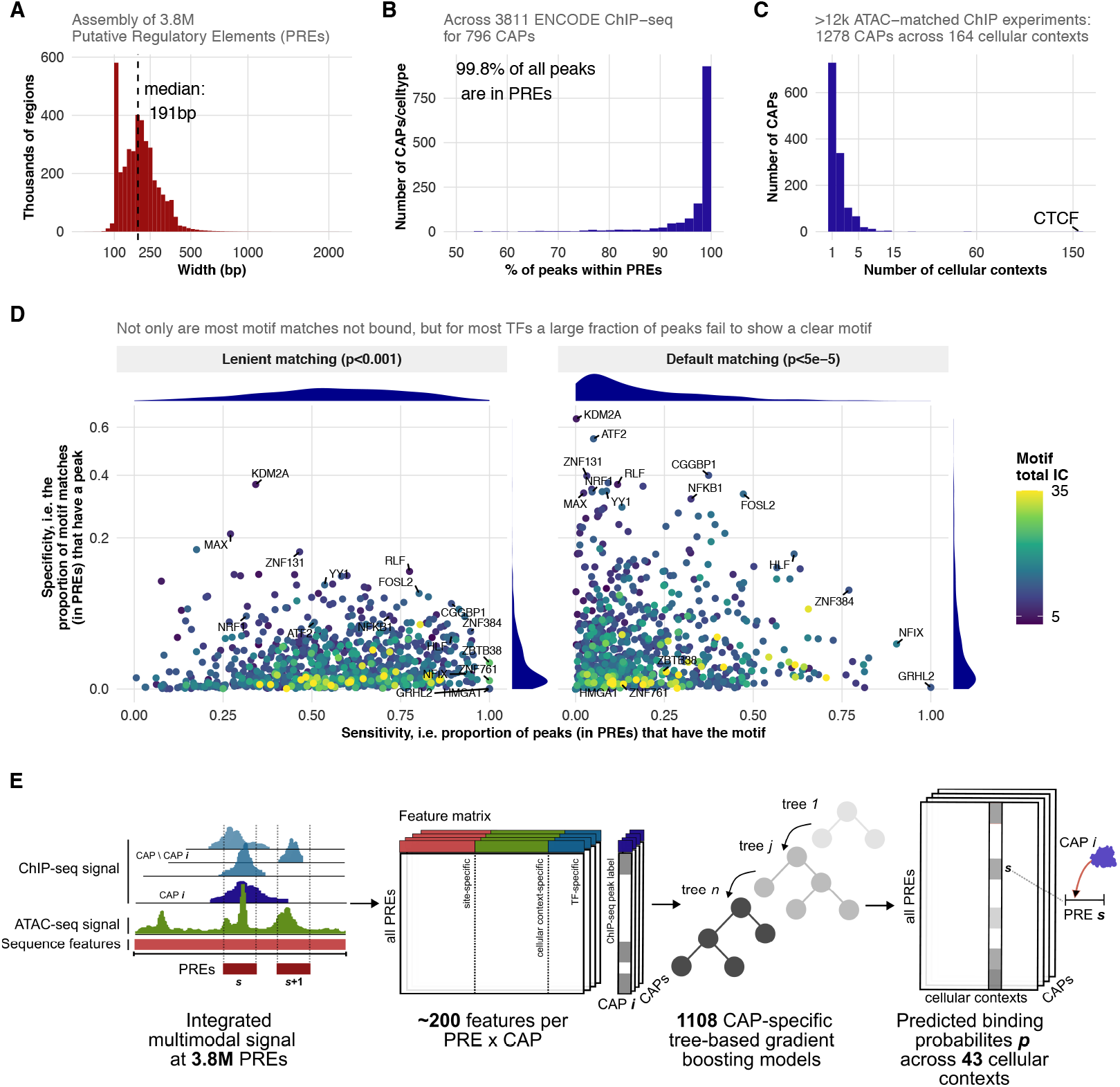
Overview data collection and prediction strategy: **A**: Width distribution of the Putative Regulatory Elements (PREs). Smaller regions were bidirectionally enlarged to a minimum of 100bp (unless limited by a neighboring region). **B**: Proportion, for each combination of ENCODE chromatin-associated protein (CAP) and cellular context, of the ChIP-seq peaks that overlap a PRE. **C**: Distribution, across CAPs, of the number of cellular contexts (i.e. combination of celltype and treatment/condition) for which (ATAC-matched) ChIP-seq data is available. **D**: Each dot is a Transcription Factor (TF) and represents the proportion of matches (in PREs) for the TF’s motif that overlap a ChIP-seq peak, against the proportion of said peaks (in PREs) that overlap a motif. The left panel represents a very loose motif matching cutoff (with the median motif matching half of all PREs), or a more commonly-used threshold (with the median motif matching 14% of DHS). Only CAPs annotated as TFs in [25] and used, however the set of motifs does include low-quality motifs when no high-quality one was available. **E:** Overview of the TFBlearner platform.

While this approach does not achieve base-pair resolution, it is well-adapted to the resolution of ChIPseq peaks, and the use of this pre-defined hypothesis space has the major advantage, for downstream applications, of being able to represent binding probabilities across TFs or cellular contexts in the form of sparse matrices.

### 1.2 Collection of matched ATAC-ChIP data

We next collected a large set of ChIP-seq experiments from multiple sources, including ENCODE, GTRD [44], Codebook [45], and BrainTF [46]. We semi-automatically matched them to ATAC profiles (see Methods A.1.5 and Supplementary Table 1), removed overlapping experiments and merged replicates, leading to 2537 combinations of 1278 chromatin-associated proteins (CAPs), of which 976 strict TFs (according to [25]), across 164 ‘cellular contexts’ (i.e. cell type × condition). These are represented in Figure 1C: as expected, most TFs were profiled in only one or two contexts.

**Table 1:** Overview of groups of constructed features: Feature groups usually result in multiple individual features in the feature matrix, depending on the TF *t* for which they are computed (see Supplementary Methods Table A4). For example, this is the case for motif matches of several distinct motifs that were taken into account or fragment overlap counts which were additional stratified by the number of nucleosomes they span. The NMF binding patterns and the C-score features are considered specific to both site and TF as for computation of these the ChIP-seq data of TF *t* was removed. Normalization across cellular contexts was performed for features based on ATAC-seq fragment or insertion counts.

| Feature Group | Feature Class | Specificity | Normalization |
| --- | --- | --- | --- |
| GC-content | Sequence | Site ● | - |
| CpG-density | Sequence | Site ● | - |
| Sequence embedding | Sequence | Site ● | - |
| Motif match scores | Sequence | Site × TF ● | - |
| Motif match counts | Sequence | Site × TF ● | - |
| Overlap counts | ATAC | Site × context ● | robust, min-$Q_{0.9}$, gbsq |
| Insertion counts | ATAC | Site × context ● | robust |
| Weighted insertion counts | ATAC | Site × context × TF ● | robust, min-$Q_{0.9}$ |
| Profile deviation | ATAC | Site × context × TF ● | - |
| Promoter association | Association | Site × TF ● | - |
| TF-activity association | Association | Site × TF ● | - |
| TF-activity | Context | context × TF ● | - |
| Context MDS | Context | context ● | - |
| NMF binding patterns | Binding patterns | site × TF ● | - |
| Cofactor binding signal | Binding patterns | site × TF ● | - |
| CTCF binding signal | Binding patterns | site × TF ● | - |
| C-score | Binding patterns | site × TF ● | - |
| Maximum overlap count | Region | site ● | min-$Q_{0.9}$ |
| Variance overlap count | Region | site ● | min-$Q_{0.9}$ |
| Conservation score | Region | site ● | - |
| Distance to TSS | Region | site ● | - |

● Site ● Context ● TF ● context × TF

The binary nature of peak calls hides differences in the occupancy and reproducibility of peaks. To address this, we used cases for which replicates were available to train a model that would predict replicability based on features of the peaks (see Methods A.1.6 and Methods A.1.6). For context-TF combinations where replicability cannot be directly assessed, this would serve as a proxy, and this replicability score as instance weights of the training of our model.

We took the opportunity of having this large collected dataset to estimate, for each TF, how informative of binding the corresponding motif is. As Fig. 1D shows, not only are most motif matches (in PREs) not bound (in any of the collected contexts), but for most TFs a large fraction of peaks fail to show a clear motif for the TF. This suggests that factors other than the TF’s sequence affinity exert a strong influence on its binding, and not simply by restricting which site is accessible.

### 1.3 TFBlearner model overview

We next trained classifiers to predict cell type-specific TF binding for a large proportion of all known human TFs (see Fig. 1C,E). Given the diversity in the binding behavior of different TFs, features were engineered to capture relevant aspects of TF binding (see Table 1 and Supplementary Fig. 3 for an overview of the features). Gradient-Boosted Decision Trees (GBDT) were selected to train TF-specific classification models as this modeling approach has been used successfully for similar tasks [38, 47] and offers several advantageous properties [48, 49]. Their tree-based structure enables the capture of non-linear relationships among features and intrinsically carries out selection of those features that are informative for a given TF. As a result, a high degree of explainability regarding feature importance is preserved, especially when combined with SHAP values [48] (see Fig. 4E,F and Methods 2.3.4). Further, efficient and well-established GBDT implementations are readily available [50].

For each TF, we trained four separate GBDT models on different data subsets, defined by lowering the threshold for inferred replicability (*r*-score) amongst the ChIP-seq peaks used as positive instances (see Methods 2.3.1 and Supplementary Methods A.1.6). Accounting for the low replicability of ChIP-seq peak calls by stacking models across levels of stringency consistently resulted in improved performance (see Supplementary Fig. 2 and Supplementary Fig. 1).

For each TF, we used model based optimization (MBO) [51] to identify hyperparameter configurations robust across celltypes (see Methods 2.3.2 and Supplementary Methods A.3.2), which tended to result in simpler models (see Supplementary Fig. 2C).

We released an R package, TFBlearner, that implements all feature construction steps, training and prediction (see Code availability and Supplementary Methods A.4).

### 1.4 TFBlearner compares favorably to alternative methods in a pre-registered benchmark

To compare TFBlearner to existing methods, a benchmark outline has been pre-registered on the Open Science Framework (OSF) [52]. Four previously developed methods that met the inclusion criteria (see Supplementary Methods A.5 and Methods 2.4.3) were identified: BMO [53], maxATAC [43], and TOP [54]. An additional method, Catchitt [37], was initially included but subsequently excluded as it repeatedly exceeded the pre-registered resource limits. Seven cellular contexts with matching ChIP-seq and ATAC-seq and having a high number of TFs from ENCODE were selected for the benchmark: two of them (H1 and Jurkat) were designated as held-out testing contexts (including a total of 32 TFs-context combinations), and the other five as training contexts (see Supplementary Table 3 for the specific datasets and Supplementary Methods A.5). Consequently, any data from the two training cell types (including other ESC or iPSC contexts) were excluded from the entire study except for the final benchmark. Supervised methods (maxATAC, TOP, and TFBlearner) were trained on the matched training ATAC- and ChIP-seq datasets, whereas BMO as an unsupervised method was directly fitted to the testing cellular contexts. Performance on the test set was evaluated on chromosomes 2, 4 and 9 (selected arbitrarily to reduce compute time), using AUPRC and precision at 5% and 10% recall, both on the set of PREs as well as on the complete chromosomes split into bins of 200 bp. The occurrence of ChIP-seq peaks (for further detail on the distinct sets used, see Methods 2.4.2 and Supplementary Methods A.5) at the PRE and, respectively, at the bins was used as ground truth.

Evaluated on the PREs, TFBlearner outperformed existing methods, closely followed by maxATAC when used with a higher-than-default number of epochs (see Fig. 2A-D, Supplementary Fig. 4, 5). The proportion of positives, i.e. of PREs overlapping a ChIP peak, varies by several orders of magnitude across TF x cellular context combinations in the test data, and is highly correlated with AUPRC (Supplementary Fig. 4E). We therefore additionally report the enrichment over random, i.e. the ratio between the AUPRC and the positive proportion. The enrichment over random shows similar trends, but with stronger distributional differences between TFBlearner and competitor methods (Fig. 2B, Supplementary Fig. 8), indicating its stable performance also on datasets with lower number of peaks.

**Fig. 2:**
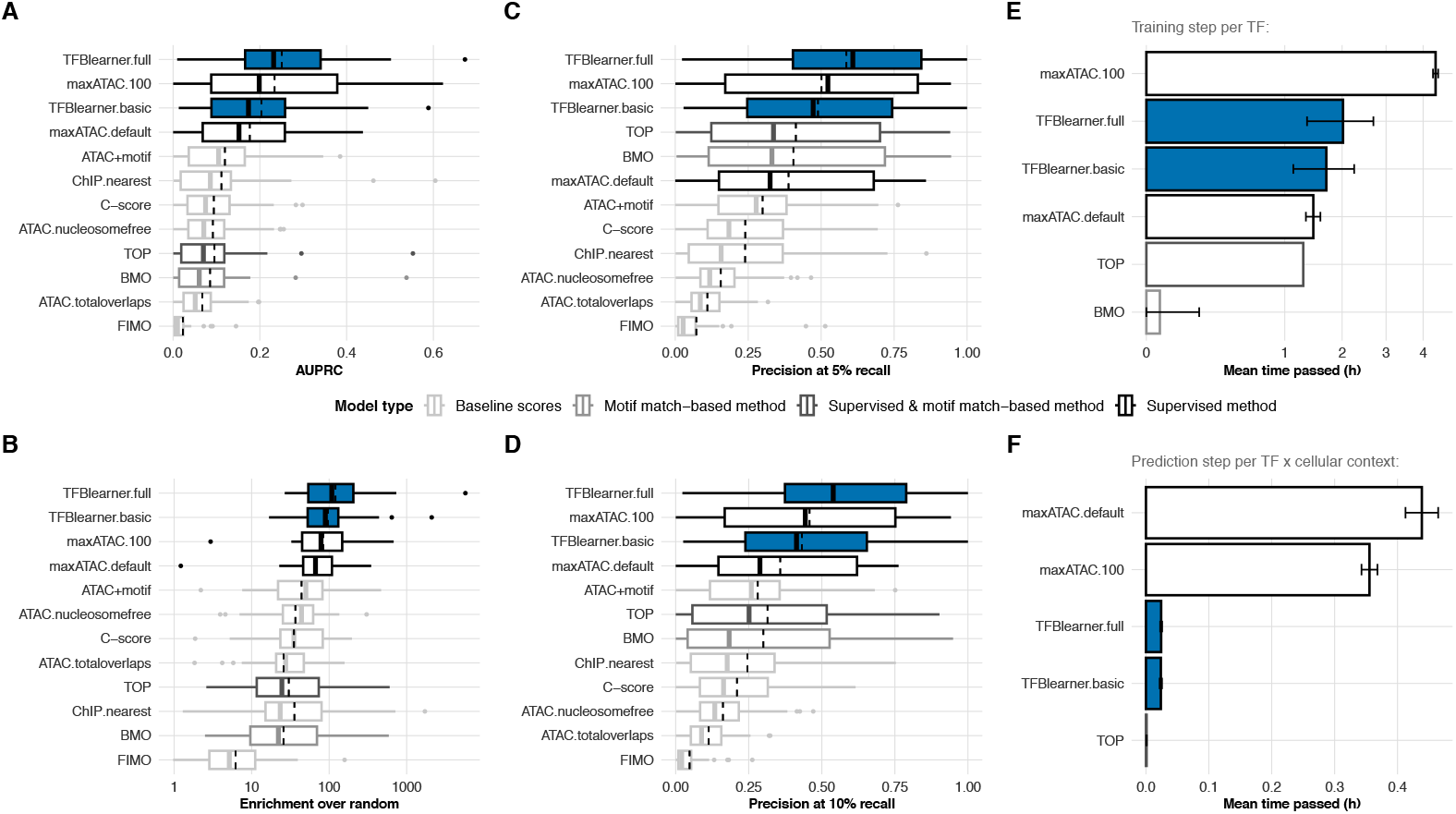
Performance of methods on pre-registered test data: **A**: AUPRC of predicted bindings assessed against ChIP-seq data of 32 TF–cellular context combinations held-out for testing. AUPRC was calculated separately for the merged peak set and the IDR-thresholded peak set and subsequently averaged across the two sets to obtain one value for each combination (Methods 2.4.3, Supplementary Fig. 5). Methods are ranked by the median (mean indicated by dashed line). Analogously for (**B**) the enrichment over random, i.e. the AUPRC divided by the proportion of positives, (**C**), precision at 5% recall and (**D**) at 10% recall. **E**: Mean time consumed by each method for training the model for one TF. For comparison, the training time of TOP given here is divided by the number of TFs as TOP is trained once for all TFs. Similarly, for BMO which is instead fitted per cellular context. **F**: Mean time consumed for predicting binding of one TF in one cellular context for all PREs on chromosomes 2, 4 and 9.

Generally, most methods achieved relatively high precision at both 5% and 10% recall, indicating their suitability for detecting top candidates (Fig. 2C,D and Supplementary Fig. 7, 8). Especially TOP and BMO, which both only predict binding at motif occurrences, showed competitive performance when restricted to the top predicted candidates.

Alongside the pre-registered methods, we also incorporated a group of easily computed baseline scores (see Supplementary Methods A.5). Particularly striking was the behavior of the context-agnostic crowdedness-score (C-score, analogous to [32]), which captures the number of distinct TFs detected as binding at a given site in any context (see also Supplementary Methods A.2.6). Also of interest is the slight performance gain when using the counts of nucleosome-free (ATAC.nucleosomefree) instead of all fragments (ATAC.totaloverlaps).

All metrics were further explored on different preregistered sets of peaks, showing mostly similar trends (see Supplementary Fig. 4A-D). Restricting the evaluation to non-crowded peaks, peaks without a motif occurrence, and excluding all peaks that appeared in any of the training contexts, proved to be harder tasks for all methods (see Supplementary Fig. 4A-D). The performance of all methods increased with higher proportions of peaks observed during training (Supplementary Fig. 9C). A more detailed exploration of this relationship showed that both top methods can distinguish whether a site with a peak observed during training will be bound or not in a new cellular context, but both have much greater difficulty predicting binding at sites where no peak was seen before (Supplementary Fig. 9A,B). In that scenario, maxATAC performed better, presumably because it relies primarily on the ATAC-seq signal and might leverage it more extensively. Consistent with this, simply using nucleosome-free read counts to identify peaks at sites not encountered during training produced in several settings better results than any supervised method (Supplementary Fig. 9A,B). Finally, when expanding the comparison to the full chromosomes (with TFBlearner predicting only in PREs), TFBlearner’s performance dropped more than that of maxATAC, however this was primarily due to a mismatch between the resolutions of PREs and bins, rather than missed positives (Supplementary Fig. 10, 11).

We also monitored computational resources used by the various steps of each method (see Fig. 2E,F and Supplementary Fig. 15, 16). Although having higher memory consumption in some steps, TFBlearner was substantially faster for steps that need to be executed for each TF cellular context combination (preprocessing and prediction), which rapidly add up in a large scale prediction effort.

### 1.5 Training and performance across 1108 CAPs

We next trained models for 1108 CAPs (Fig. 1E), of which 960 are *bona fide* transcription factors [25]. We first confirmed, across this large set of models, an absence of overfitting by comparing performance on training versus held-out chromosomes (Fig. 3A). For each CAP with more than one training cellular context (459 CAPs, and 1282 CAP-context combinations), we then ran leave-one-cellular-context-out (LOCO) rounds to evaluate performance, and determinants thereof, on unseen held-out cellular contexts (see Supplementary Methods A.3.3). As expected, AUPRC in held-out context increased with the number of available training contexts (Fig. 3B). However, an even stronger relationship was found between the AUPRC and the proportion of positives in the held-out contexts, as had been observed in the benchmark (Fig. 3C). The relationship was considerably weaker with the rate of positives in the training data (see Supplementary Fig. 17A,B), confirming that this is a mathematical feature of the AUPRC evaluation in the held-out data, rather than the availability of training data or features of the TFs. We therefore again resorted to compute the fold-increase over the base rate of the AUPRC, which showed a relatively stable distribution across the profiled TF-context combinations (Fig. 3D), with an average of a 95-fold enrichment (median 60-fold). Analogously to the benchmark, the main determinant of the performance in a held-out context is the proportion of peaks already observed in the training contexts (Fig. 3E, Supplementary Fig. 17A,B), which might explain the dependence on the number of training contexts. Nonetheless, also in this case the models could determine whether a site that showed a peak during training would be bound or not in a new context (see Supplementary Fig. 18, showing dependence of performance for different overlap proportions), ruling out a pure memorization effect. The proportion of peaks detected during training provides an estimate of how similar TF binding is across different contexts, and accordingly it also did exhibit a clear association with the performance difference between held-out and training data, quantified as the log2 ratio of their respective enrichments (Fig. 3G, Supplementary Fig. 17C,D). On this basis, we regressed out the effect of the overlap proportion and the base rates from the difference to estimate, for each TF, a generalizability of the model (see Supplementary Methods A.3.4). While both performance and generalizability showed large variability across TFs, some patterns emerged from the average of TF classes (Fig. 3G). In particular, forkhead family factors, which are relevant for several key cellular functions and include factors with strong pioneering activity [55], showed relatively high generalizability.

**Fig. 3:**
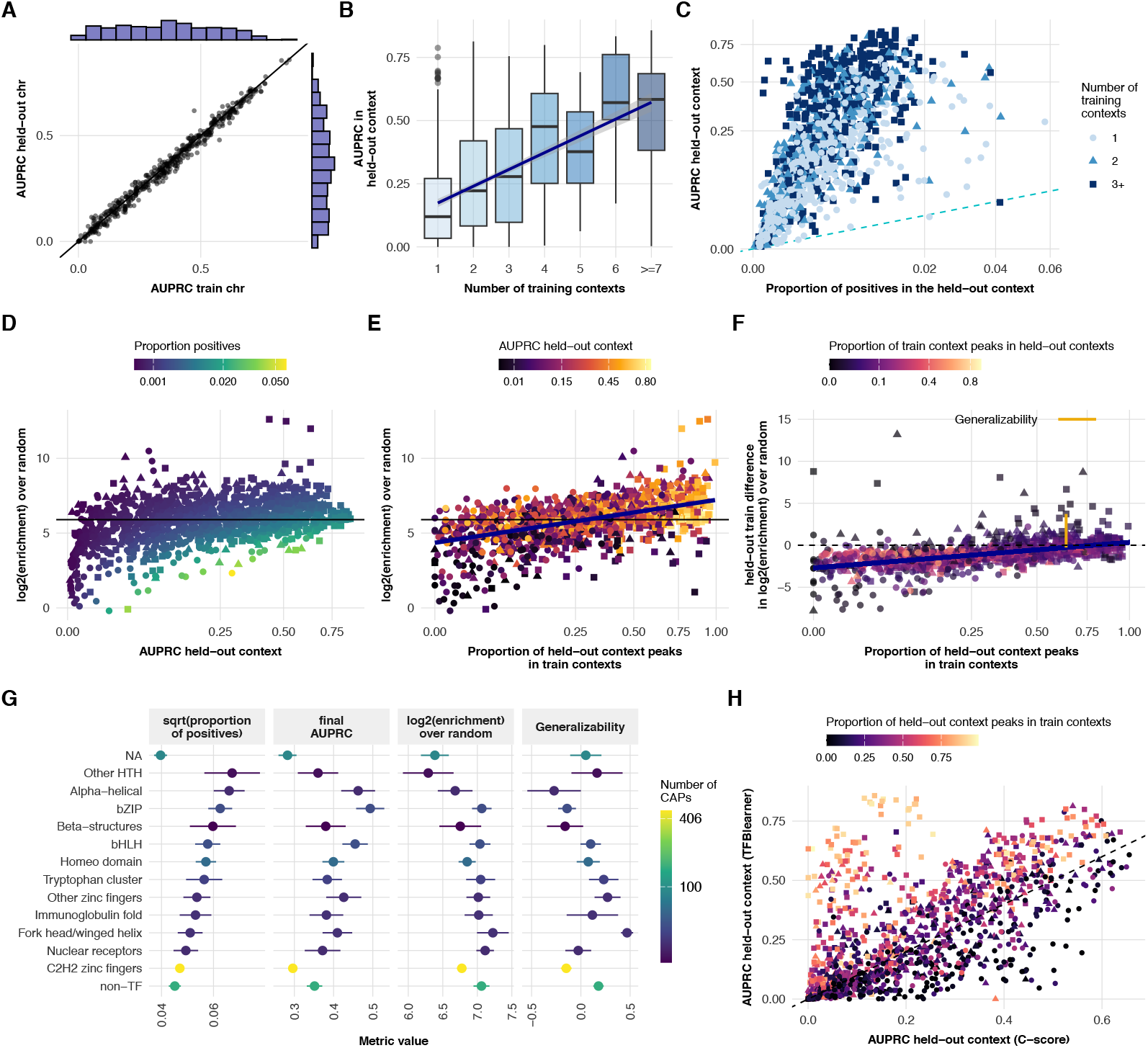
Performance on held-out contexts during model training for the 459 CAPs profiled in more than one context: **A**: AUPRC on chromosomes held-out from training contexts (2,4,9) versus chromosomes seen during training. **B**: AUPRC in held-out contexts during training (LOCO rounds, see Supplementary Methods Fig. 4) against the number of contexts used for training. **C**: Dependence of AUPRC in the held-out on its positive proportion. **D**: Log2-transformed enrichment over random, i.e. AUPRC in the held-out divided by the respective positive proportion, versus the AUPRC. **E**: Log2 of the enrichment over random in the held-out context in dependence of the proportion of peaks in the held-out already observed in at least one of the training contexts. **F**: Difference in performance between the held-out and training contexts, expressed as the log2 ratio of their respective enrichments relative to random shown in dependence of proportion of held-out peaks seen in training. Generalizability was estimated as the residuals when regressing out the effect of the proportion seen during training and other relevant determinants on the performance difference between held-out and train (Supplementary Methods A.3.4). **G**: Mean proportion of positives, model performance and generalizability across TF families. Shown are the mean and standard deviations across CAPs. The grouping is based on HOCOMOCO [56] annotations, themselves based on [57]; the hierarchical tree was cut at different heights to yield groups of TFs with more comparable sizes. ‘non-TF’ indicate CAPs not annotated as TF by [25], and ‘NA’ indicates Lambert TFs without HOCOMOCO family annotation. **H**: AUPRC on held-out contexts obtained by TFBlearner compared with AUPRC obtained by the C-score. Note, that the C-score shown here is not adjusted for the width of the site it is computed at.

Together, these results indicate that TFBlearner has a relatively stable performance across a large set of TFs, and that the generalizability of its predictions depend on biological factors.

### 1.6 Predictive features are informative about the TFs

We next investigated whether the importance assigned by the models to predictive features could yield insights into their respective TFs. In general, accessibility and cofactors were the most important groups of features, much more so than the motif(s) (Fig. 4). We therefore tested whether the importance assigned to cofactors differed across classes of TFs, first comparing monomeric (or homo-multimeric) binding TFs to obligate heteromers (Fig. 4A), finding as expected that the latter were more dependent on cofactors. In contrast, pioneers and TFs without intrinsically-disordered domains (important for interactions between TFs) were on average less dependent on cofactors (Fig. 4B-C). Surprisingly, we also found that TFs containing only repressive effector domains tended to be less dependent on cofactors (Fig. 4D). Feature importance also varied across TF classes (Fig. 4E-F), in ways that did not entirely recapitulate previously known characterizations of their binding. For example, the importance attributed to the TFs’ motif(s) was not associated to the motifs’ information content (IC), nor was the importance of ATAC features associated with the normalized chromatin information content (nVICE) as estimated by [53]. Since each model underwent its own hyper-parameter optimization, we reasoned that the models’ complexity could also yield insights onto the TFs. Generally, model complexity was strongly associated with the number of positives in training (Supplementary Fig. 17E), and more weakly associated to the strength of regularization. Adjusted for the amount of training data, model complexity also varied across TF classes, being for instance lower for pioneers (Fig. 4F). Finally, we confirmed that the models assigned the expected importance to major known cofactors (Fig. 4G). In summary, features of the models reflect features of their respective TF.

### 1.7 A compendium of binding predictions across major human cell types

A major roadblock to the adoption of binding prediction methods in broader biological research is the need for users to run predictions on their cell type(s) of interest. Therefore, to increase the accessibility and ease of use of our models and predictions, we predicted the binding of 1108 CAPs (of which 960 TFs) in 43 cell types (see Supplementary Table 4). This includes some of the cell lines most widely used in fundamental biology, as well as representatives of most major (nucleated) cellular components of the human body, with an emphasis toward brain cell types. All predictions are publicly available for download and browsable through our TFBPlatform, enabling researchers to visualize accessibility and TF binding predictions in regions of interest (selected from a genome browser). In addition, detailed information about the TF-specific models are also browsable, such as their performance and feature importance, providing insights into the TFs themselves.

### 1.8 Usage of the binding predictions for TF activity inference

We next investigated whether this compendium of binding predictions could be used to infer differential TF activity. We first evaluated relative TF activity inference from ATAC-seq data, using the human benchmark datasets from our previous study on the topic [17]. Specifically, we first implemented a faster version of chromVAR [58] that uses binding probability scores instead of binary motif absence/presence, and ran it using both predictions and motifs (see Supplementary Methods), followed by limma differential analysis. To simulate a realistic application setting, we did not use predictions based on condition-specific ATAC-seq profile, but generic ones from the same cell type, and ran the analysis using only ATAC peaks (as opposed to all PREs) as is usually done in practice.

Surprisingly, given their considerably higher predictivity for binding, the raw predictions were substantially worse than motifs for activity inference from ATAC-seq (Fig. 5A-C). We hypothesized that this could be due to the high correlation between the prediction probabilities of different TFs (chiefly due to the underlying accessibility pattern), which lead to large sets of TFs showing similar differential activity patterns between the conditions. We therefore re-weighted the prediction probabilities by regressing out the median probability across TFs. This lead to a substantial improvement in the rank and *t*-value of the true TF (Fig. 5A-B). Nevertheless, although the predictions enabled an accurate identification of Myc that was missed by the motifs, across datasets they did not bring a substantial general improvement. It could however help distinguish between TFs of the same family, which share highly-similar motifs, such as steroid hormone receptors in the example of NR3C1.

**Fig. 4:**
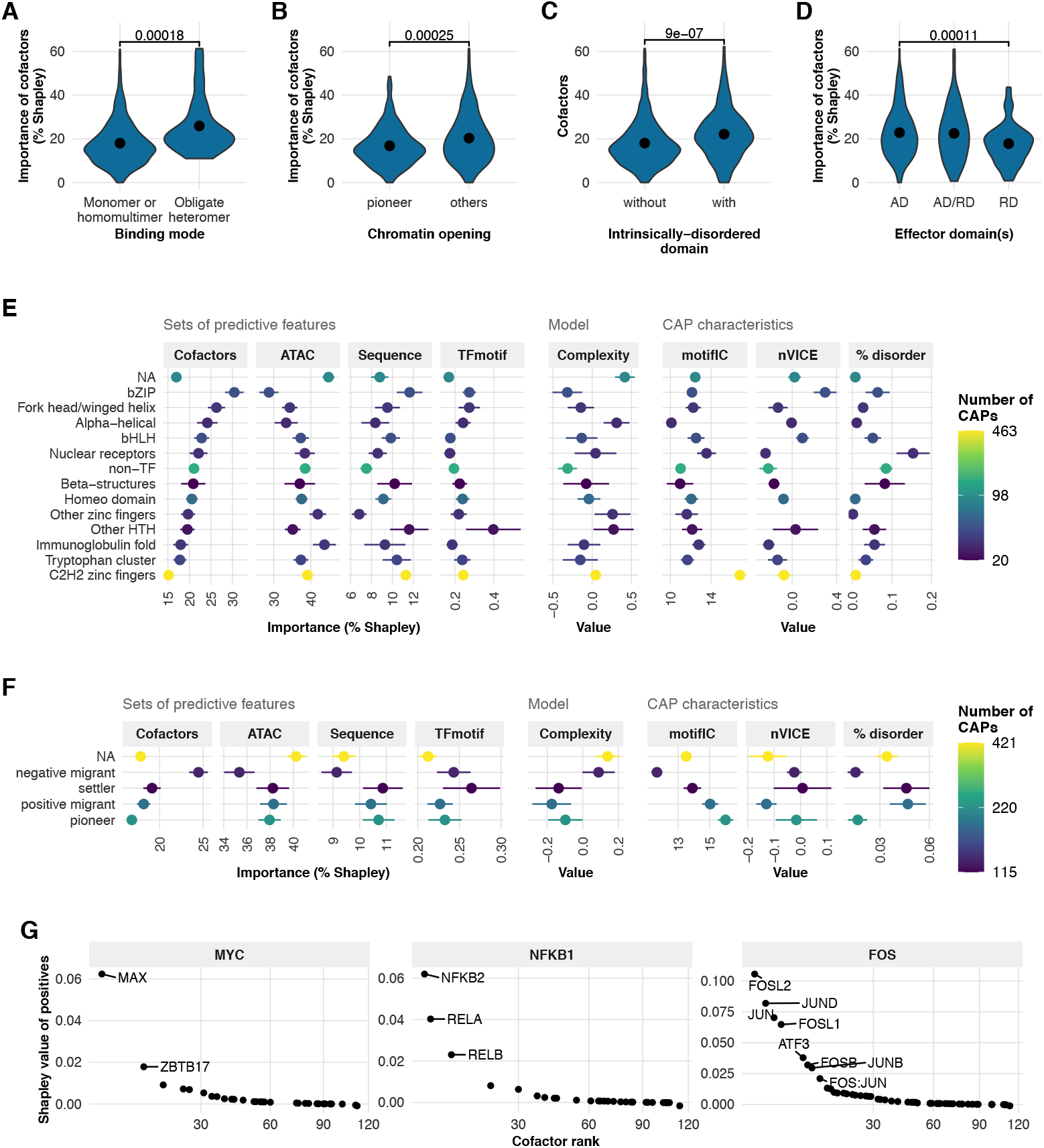
Model characteristics are informative of the TF: **A-D**: The predictive importance of cofactor information is higher for heteromers (**A**), lower for pioneers (**B**), higher for TFs containing intrinsically-disordered regions (**C**), and lower for repressors (**D**). **E-F**: Predictive importance of sets of features (first 4 columns), model complexity (middle), and other characteristics of the TFs (right) across TF families (**E**) and TF chromatin opening types (**F**). **G**: Examples of importance assigned to key cofactors of well-known TFs. Feature importance are available, along with predictions, on http://www.ethz-ins.org/TFBPlatform/.

**Fig. 5:**
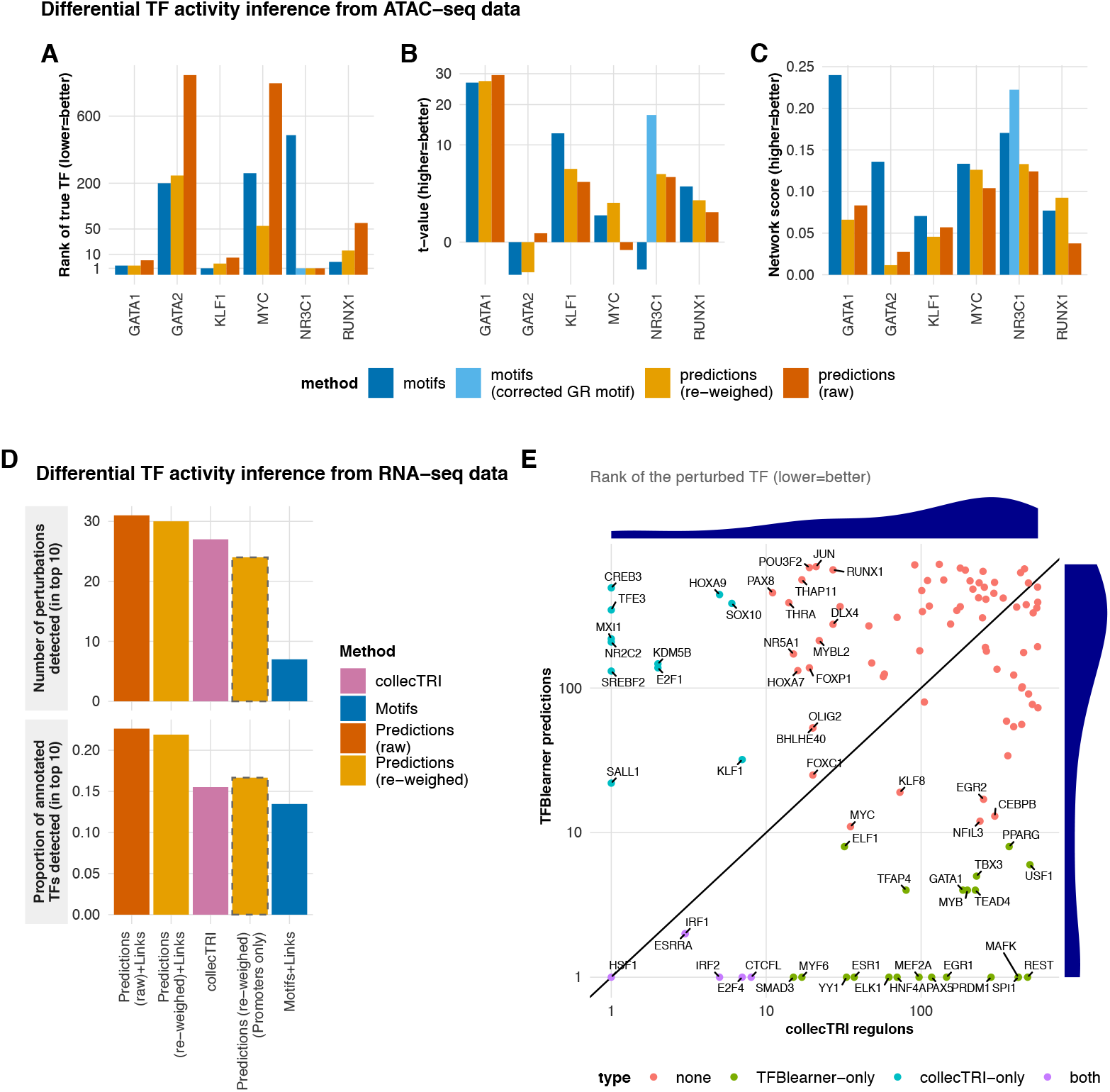
Usage of the binding predictions for TF activity inference. **A-C**. Differential TF activity inference from ATAC-seq data, using the human benchmark datasets from [17], showing the rank of the perturbed TF in the differential analysis (**A**), its *t*-value (**B**), and the network score (**C**). The t-values were inverted for TFs that were knocked-down so that the expected value is positive. **D-E**. TF activity inference from the transcriptome, using the dataset from [59]. **D**. Inference based on the predictions retrieves more of the perturbed TFs among the top 10 differentially-active TFs in each condition than alternatives, both in absolute terms (top) and relative to the number of perturbed TFs in the respective annotation (bottom). **E**. Comparison of the rank of each perturbed TF in inferences based on either our predictions or collecTRI (showing only TFs annotated in both), highlighting the complementarity of the predictions.

We next hypothesized that the binding predictions could instead improve activity inference from the transcriptome, which (contrary to ATAC-seq) does not provide information as to whether a putative binding site is actually occupied or not. To investigate this in a context completely held-out from our study, we used a dataset individually over-expressing a large number of TFs in pluripotent stem cells [59]. To link distal PREs to target genes, we used ENCODE rE2G predictions from the same cell type (see Methods 2.5.2), and used the cross-product of the binding prediction matrix with that of the region-to-gene matrix, yielding a TF-to-gene weight matrix. As baseline, we compared this with that using motifs instead of predictions, as well as the collecTRI regulons [19], which represent the state-of-the art resource for the inference of TF activity from the transcriptome. In all cases, we used the same unsupervised method to assign inferred TF activity scores based on the (weighted) regulons and a given transcriptome: multivariate-regression (see methods), which showed good performance in previous studies [17, 18]. In contrast to inferences from ATAC-seq, the predictions yielded a substantial improvement over motifs, and also outperformed the collecTRI regulons in terms of the number of perturbed TFs identified in the top 10 predictions (Fig. 5D, top). Importantly, the collecTRI regulons include a larger number of TFs among the perturbed set (Supplementary Fig. 19A), giving it an unfair advantage. As a proportion of annotated TFs, the relative performance of the predictions was even stronger (Fig. 5D, bottom). Re-weighing the predictions, as done for their use with ATAC-seq data, did not improve inference based on the transcriptome (Fig. 5D). These results were entirely reproduced when using logFCs rather than logCPMs as input (Supplementary Fig. 19B). Importantly, the TFs correctly identified using collecTRI and the predictions only partially overlapped (Fig. 5E and Supplementary Fig. 19C-D), indicating that, rather than one annotation superseding the other, the two approaches are complementary.

In summary, these results show that, at least with current methods (developed for motifs), the compendium of binding predictions offers limited advantage over motifs for TF activity inference from accessibility data, chiefly enabling to distinguish TFs with similar motifs. In the case of inference from the transcriptome, however, the binding predictions offer substantial improvement, showing improved recovery rates over state-of-the-art resources.

### 1.9 Crowded bindings show cell type specificity and are potentially functional

The surprisingly high performance of context-agnostic crowdedness in predicting binding, as well as the relatively low performance of (raw) predictions for TF activity inference from ATAC-seq, suggest a low specificity of TF occupancy as measured by ChIP. A particularly extreme example of low-specifictiy binding are so-called ‘crowded’ of HOT regions [27]. Taking advantage of the large dataset we assembled, we sought to investigate this phenomenon using the five cell types for which we have ChIP data for more than a hundred CAPs (Fig. 6).

**Fig. 6:**
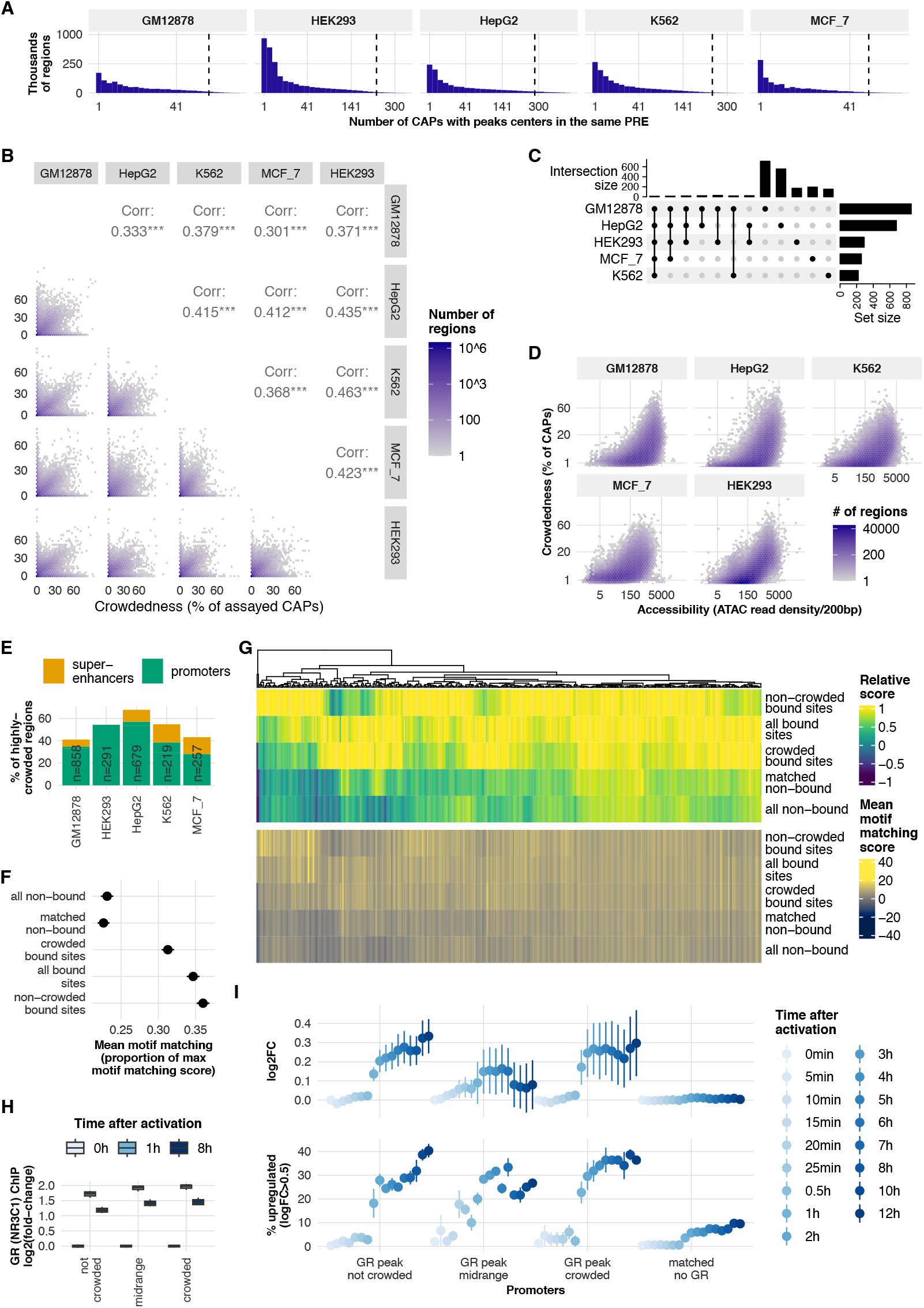
Cell type specificity and response of crowded regions: **A**: Distribution of the number of CAPs for which a peak center overlaps each PRE. The dashed line indicates half of the assayed CAPs. **B**: The crowdedness score is poorly correlated across cell types. **C**: Highly-crowded regions (*>* 40% crowdedness) are highly cell type-specific, suggesting that they are determined by biological rather than technical processes. **D**: All crowded regions are hyperaccessible, but not all hyper-accessible regions are crowded. **E**: Proportion of highly-crowded regions overlapping promoters (−2kb/+200bp) or cell type-specific super-enhancers (from [60]). **F**: Mean motif matching scores of peaks in crowded vs non-crowded (*<* 20% CAPs bound) regions (matching scores are relative to the maximum motif matching score obtained for each CAP, averaged across regions, and plotted are SEM across CAPs). While peaks in crowded regions tend to have lower matching scores as those in non-crowded regions, both are still much higher than in unbound regions. **G**: Comparison of motif matching in crowded, non-crowded and non-bound regions across CAPs. In general the vast majority of CAPs show stronger mean motif matching in crowded peaks than in unbound regions. While a subset of CAPs show stronger motif matching in non-crowded peaks than in crowded peaks, a similarly-sized subset shows the opposite pattern. **H**: Impact of GR activation by dexamethasone in different subsets of GR peaks (non-crowded regions sampled to have a baseline accessibility distribution comparable to crowded ones). Individual data points are the average across regions for each biological replicate. GR peaks in crowded regions show a similar GR binding dynamics (i.e. relative GR ChIP-seq signal) to those in non-crowded regions. **I**: Crowded GR-bound promoters respond to dexamethasone similarly to (accessibility-matched) non-crowded ones. The top panel shows the mean and standard error of the mean logFC of promoters in each set, while the lower panel shows the mean and standard error (across samples) of the proportion of promoters in each set that is upregulated.

While the vast majority of PREs showed binding by a limited number of CAPs, the distribution had a surprisingly long tail [30], with some regions being bound by over half of the assayed CAPs (Fig. 6A). We defined a crowdedness score as the proportion of the assayed CAPs in that cell type having a peak center in the region, normalized to a 200bp width. Contrarily to previous reports of HOT regions being ubiquitously expressed [28], crowdedness was only partially correlated across cell types (Fig. 6B), and the most highly-crowded regions (*>*=40% of CAPs) showed extensive cell type specificity (Fig. 6C), especially showing an on-off pattern (i.e. either crowded or hardly bound at all). In each cell type, crowded regions were hyperaccessible, but not all hyperaccessible regions were crowded (Fig. 6D), suggesting that accessibility might not be the only driver of crowdedness.

Across cell types, most highly-crowded regions were in promoters and super-enhancers (Fig. 6E), in line with previous reports [27, 33, 35]. Despite their limited overlap across cell types, highly-crowded promoters showed strikingly similar Gene Ontology enrichments for terms related to RNA-processing (Supplementary Fig. 20A). Across cell types, ChIP targets most associated with crowdedness were generally associated with activity (e.g. EP300) and in particular included early response TFs (e.g. CREM/CREB1, JUNB, etc.) as well as YY1 (Supplementary Fig. 20B). In contrast, targets least associated with crowdedness were often repressive (e.g. members of the polycomb repressive complexes).

It had previously been reported that crowded regions, although generally enriched in motifs, tended to show weaker motifs than peaks of the same TFs in non-crowded regions [27, 32]. We therefore evaluated the average motif matching score of peaks in crowded and non-crowded regions (Fig. 6F-G). While peaks in crowded regions showed significantly weaker motif matching than in non-crowded ones (Fig. 6G), the difference was mild, and both were much higher than in accessibility-matched non-bound regions. Indeed, while some CAPs showed much weaker motifs in crowded peaks, a large fraction showed little difference, and some even showed higher matching (Fig. 6G). As there have been debate in the literature on whether such high occupancy regions could be chiefly artifacts [30–34], we next sought to evaluate whether they displayed response to stimulation. To this end, we used ChIP and high-resolution transcriptomic data following ligand-based activation of the glucocorticoid receptor (GR) in A549 cells [61]. GR peaks showed the same dynamic pattern of strong relative gain followed by decrease of ChIP-seq signal in crowded and accessibility-matched non-crowded region (Fig. 6H). Since crowded regions tend to be promoters, we further assessed whether the corresponding genes changed in expression upon activation (Fig. 6I). On average, crowded GR-bound promoters increased expression upon activation similarly to non-crowded GR-bound promoters.

Together, these results support a model in which a large number of CAPs bind to the same hyper-accessible regions, in a cell type-specific fashion, and that these ChIP-seq peaks are not artifacts but show the expected dynamics upon stimulation and lead to functional impact on expression.

### 1.10 Peaks in crowded regions can be explained by distal elements

The crowded regions raise the question of how so many proteins can occupy relatively small stretches of DNA. We formulated three, non-mutually exclusive hypotheses (Fig. 7A). The first is that the different CAPs do not simultaneously occupy the region in a given cell, but dynamically compete for binding, so that while only a subset is bound at a given time in a given cell, averaged over a population they all appear to occupy the region. An implication of this ‘dynamic exchange’ hypothesis is that the CAPs would have a lower occupancy in crowded versus non-crowded regions. Using our ChIP signal-based replicability score as a proxy for occupancy, we tested this hypothesis, and found that, if anything, peaks in crowded regions were more reproducible (and hence had a stronger ChIP signal) than those in accessibility-matched non-crowded regions (Supplementary Fig. 20C). A limitation of this data is that, since crowded regions are hyper-accessible, it is plausible that at that high range the ChIP signal loses its quantitative relationship to occupancy.

**Fig. 7:**
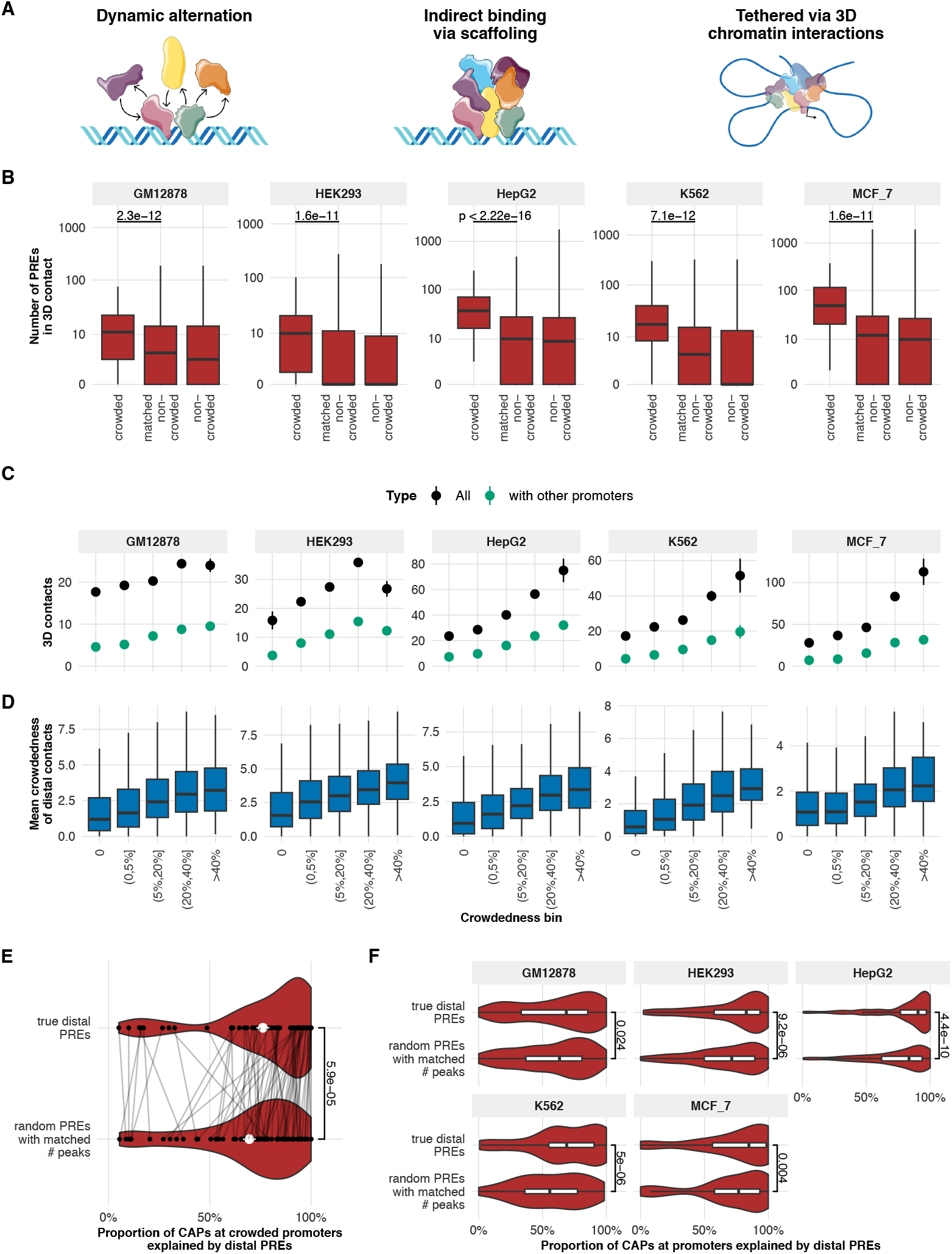
Peaks in crowded regions could be explained by distal elements: **A**: Schematic representation of the three hypotheses. **B**: Number of PREs in 3D contact with highly-crowded (*>*= 40%) and (accessibility-matched) non-crowded (*<* 20%) promoters, based on ChIA-PET data from the respective cell type. **C**: Mean and standard error (across promoters of each set) of the number of 3D contacts (and specifically those with promoters) of promoters in the different crowdedness bins. **C**: Average crowdedness of regions in 3D contacts with promoters of different crowdedness bins. **E**: For most highly-crowded promoters, the majority of the peaks found in the promoter are also found at associated distal PREs, and the proportion is significantly higher than in matched random PREs. (Crowded promoters from all cell types merged together.) **F**: When extending to more promoters (crowdedness score *>*= 20%), the effect is significantly reproduced across all cell types.

A second hypothesis is that only a subset of the TFs directly bind to DNA, creating a scaffold on which others can bind. Under this model, only a subset of the CAPs at crowded sites would show a strong motif, consistent with our observations (Fig. 6F-G). We hypothesized that such scaffolding might render the peaks dependent on cross-linking commonly used in ChIP. We therefore turned to data from Cut&Run, which avoids cross-linking, to test whether the crowded peaks tended to be less reproduced than non-crowded ones. Although we could find, among our cell lines, Cut&Run data only for Myc and Max, these did not show any trend towards a dependence of crowded peaks on cross-linking (Supplementary Fig. 20D).

Finally, a third, related hypothesis is that crowded promoters are in 3D contact with many distal enhancers, and that many of the CAPs whose peaks are found at crowded promoters are primarily directly bound to the enhancers, but due to 3D contact get immuno-precipitated with the promoters. An implication of this would be that crowded promoters have 3D contact with more regions that non-crowded ones. To test this hypothesis, we used ChIA-PET data collected in the ENCODE Screen [62] in the respective cell lines. Across all cell lines, highly-crowded promoters had on average significantly more contact with PREs than accessibility-matched non-crowded ones (Fig. 7B). There was positive relationship between the crowdedness of a promoter and the extent to which it formed 3D contacts, with an increasing proportion of contacts with other promoters (Fig. 7C).

Under this model, when a given region is immunoprecipitated, its contact regions would also tend to be. Indeed, we observed that the distal contacts of more crowded regions tend to also be more (Fig. 7D). A further implication of this hypothesis is that a large fraction of the CAPs found to bind at the promoter would also be found at some of their enhancers, which turned out to be the case in the vast majority of promoters (Fig. 7E). Importantly, the bindings at promoters were significantly better explained by its associated distal PREs than by randomly-selected PREs that have similar numbers of peaks (Fig. 7E). We further extended this pattern beyond highly-crowded promoters, and found that it holds across all cell lines individually (Fig. 7F). Under this model, some CAPs would bind directly at the promoters, and should thus have a strong motif match, whereas others would be tethered and have weaker motif matches at promoters (and presumably strong ones at some distal enhancers). To test this, we looked for an association between whether CAPs tend to bind more distally or closer to TSS, and whether the exhibited strong reduction in motif matching score in bound crowded sites compared to control bound sites. As predicted, CAPs that tend to bind closer to TSS show as strong a motif match in crowded versus non-crowded bound sites (and tend to be better predicted by crowdedness), whereas those binding more distally show weaker motifs in crowded bound sites (Supplementary Fig. 20E).

A limitation of these findings is that even though we compare crowded regions to control regions with similar ATAC-seq signal, it is possible that this signal is saturated, and that a technology with a higher dynamic range would show crowded regions to have even higher accessibility. Therefore, while convergent evidence suggests that crowdedness may be driven by many CAPs being brought in contact to the crowded regions through bindings at a large number of distal regulatory elements, experimental evidence will be necessary to confirm this hypothesis.

## 2 Methods

### 2.1 Definition of the Putative Regulatory Elements (PREs)

As a starting point for PREs, we first used consensus DNAseI hyper-sensitive sites (DHS) from [63] (ENCODE file ENCFF503GCK), to which we added per-cell type nucleosome-free peaks from the single-cell ATAC-seq atlas by [64] (GSE184462). For the latter, we split the authors’ aligned fragments by their cell type annotation, extracted fragments shorter than 125bp, and called peaks on those using macs2 [65] with `-q 0.01 -f BEDPE --keep-dup`. Those were joined to the DHS (see Supplementary Methods A.1.4). This yielded 2,868,116 non-overlapping regions. Because mouse and human datasets tend to overlap only partially in terms of experimental system, we employed the same procedure on mouse DHS (ENCODE file ENCFF910SRW) and a mouse brain scATAC dataset [66]. We lifted the mouse regions over to human coordinates, remove regions i) without synthetic human region, ii) with gaps larger than 50bp, or iii) of a width larger than 2000bp, and added the remaining that did not already overlap previous human regions to the PREs. Due to its high variability across ATAC protocols, we removed regions from the mitochondrial chromosome. The resulting 3,216,635 regions had a fairly wide width distribution, ranging from 20 to 8560bp. To get a narrower distribution, we first extended the regions from their center to a minimum of 100bp (or until encountering the next region). We next split the regions in places that had low overlap for features (nucleosome-free peaks and matches for motif archetypes from [67]), i.e. troughs, using the `resplitRegions` function (see Supplementary Methods A.1.4). The resulting final set of PREs E contains 3,799,654 elements.

### 2.2 Feature construction

For each PRE site *e* = {1, .., *l*} in a cellular context *c*, a set of features of different classes was obtained for a TF *t* (or more generally, a CAP but for simplicity, referred to as TF in this section). Features can be grouped according to two schemata, based on the data they utilize and the biological characteristics they aim to capture or on their level of specificity (see Table 1).

#### 2.2.1 Sequence features

Different sequence characteristics were collected for each PRE *e*.

##### GC-content

Proportion of G or C nucleotides in the sequence of *e*.

##### CpG-density

Proportion of CG or GC dinucleotides in the sequence of *e*.

##### Motif match scores

TF motif matching scores were collected for the canonical motif of TF *t* and motifs of known cofactors. Motif matching scores were also retained for motifs whose occurrences were identified as either co-occurring with or mutually exclusive to the ChIP-seq peaks of *t* in the training data (Supplementary Methods A.2.1). In total, five co-occurring motifs and five mutually exclusive motifs were selected. In addition, motif matching scores of CTCF, dimer and dinucleotide motif matching scores of *t* were obtained if available.

##### Motif match counts

For the motifs of *t* and the known cofactors also the number of matches within *e* were computed.

##### Sequence embeddings

An extra set of sequence features was obtained via StarSpace [68] sequence embeddings to capture potential grammar of the broader sequence context (Supplementary Methods A.2.2).

#### 2.2.2 ATAC features

For each PRE *e* in a context *c*, a set of features was computed directly based on the chromatin accessibility profile of the respective cellular context.

##### Overlap counts

Total counts of ATAC-seq fragments overlapping a site (by ≥ 1bp) were obtained. The counts were additionally stratified for nucleosome-free, mono-, di- and multi-nucleosome containing fragments.

##### Insertion counts

Insertion counts were obtained by counting specifically the fragments for which either end falls within the site. The same stratification of fragment types as for the overlap counts was applied for the insertion counts.

##### Weighted insertion counts

Weighted insertion counts were computed using an insertion weight profile (see Supplementary Methods A.2.3) to weight each insertion depending on its position relative to a motif match of TF *t*. Weighted insertion counts were calculated separately for insertions within the motif match and those occurring in the flanks of it (±200bp around the motif center). Therefore, weighted insertion counts were obtained only for sites *e* containing a motif match of *t*. Analogously, insertion counts with uniform weights in the vicinity of the motif match were also computed.

##### Profile deviation

At sites containing a motif match, Chi-squared (*χ*^2^)-deviations of the observed insertion counts from the insertion weight profile were used as an additional feature (see Supplementary Methods A.2.3).

#### 2.2.3 Association features

A class of features computed for each site *e* that quantify how chromatin accessibility at that site relates to the signal at other sites across cellular contexts.

##### Promoter association

Association between the GC smooth-quantile normalized (gbsq-normalized) total overlap counts [69] at site *e* and at the promoters of the TF *t* computed across all contexts with ATAC-seq data available. Pearson correlation and Cohen’s kappa were computed as measures of association.

##### TF activity association

Pearson correlation between the gbsq-normalized total overlap counts at site *e* and the chromVAR [58] TF activity estimates (see Supplementary Methods A.2.4).

#### 2.2.4 Cellular-context features

Meta-features which were derived from chromatin accessibility and that are specific to the corresponding cellular context.

**TF activity estimates:** chromVAR TF activity estimates for the motif of TF *t*, cofactor motifs, CTCF motif and the co-occurring or mutually exclusive motifs (Supplementary Methods A.2.1) were obtained in the respective context.

**Cellular context MDS:** Multidimensional scaling (MDS) coordinates of training cellular contexts. MDS was computed based on the double centered euclidean distance matrix of all contexts. Distances were determined on the total overlap counts at the the top 10^5^ sites with highest variance.

#### 2.2.5 Binding patterns

Features which were computed for each site *e* based on ChIP-seq data of other TFs than TF *t*.

##### NMF binding patterns

Non-negative matrix factorization (NMF) was applied to the ChIP-seq matrix *B*, containing the *r*-scores for a given TF–cellular context pair (column) across all sites *E* (rows). Prior to NMF all columns with specifications corresponding to TF *t* or testing cellular contexts were removed and *B* was aggregated in two distinct ways: (1) across TFs, by taking, for each TF at a site *e*, the maximum *r*-score observed across all its covered cellular contexts, and (2) across TFs within each context, taking, for a given site *e* in a context, the maximum *r*-score across all covered TFs. A L1-regularized version of NMF by alternating least squares was used (RcppML R package [70]) to decompose each of the two aggregated ChIP-seq matrices into a lower-rank (*k* = 40) feature matrix *W* and coefficient matrix *H* (see Supplementary Methods A.2.5). The feature matrices *W* of dimension |*E*| ×40 of both aggregations were subsequently used as binding pattern features.

##### Cofactor binding signal

Mean *r*-sores of cofactors of *t* at site *e* computed across the respective covered contexts in *B*. Scores were computed separately for each cofactor of *t* provided.

##### CTCF binding signal

The maximum CTCF *r*-score at site *e* obtained across all contexts in *B* with CTCF covered.

##### C-score

Sum of per-TF mean *r*-scores at a site *e*, mean scores were computed for each TF (except *t*) across the respective contexts in *B* (see Supplementary Methods A.2.6).

#### 2.2.6 Region Features

Features that were computed for each individual PRE site *e*, but that do not depend only on the sequence or signal at that site itself.

##### Maximum chromatin accessibility

The maximum total overlap count observed at site *e* across all contexts with available ATAC-seq data was obtained. Prior to identifying the maximum, total overlap counts in each context were normalized by the minimum-*Q*_0.9_ range (min-*Q*_0.9_).

##### Variance of chromatin accessibility

The same procedure was applied to compute variance of the total overlap counts at site *e*.

##### Conservation scores

UCSC hg38 sequence conservation scores were obtained based on multiple alignments of 100 vertebrate species by PhastCons, using the phastCons100way.UCSC.hg38 R package [71].

##### Distance to TSS

Distance from site *e* to the TSS of the closest TF gene was calculated with the distanceToNearest function from the GenomicRanges R package [72].

#### 2.2.7 Normalization

Feature matrices for a TF *t* were obtained separately for all its training cellular contexts, i.e. cellular contexts with matched ATAC-seq and ChIP-seq data for the respective TF. Feature matrices of different cellular contexts were normalized and row-bound into a final per TF training matrix (see Supplementary Methods A.3.3). Each ATAC-seq based cellular-context specific feature *m* (see Table 1) was normalized for each context *c* by robust normalization:

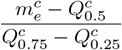

Where *Q*_0.5_ is the median of the feature in context *c* and the denominator the corresponding interquartile range. Additionally, total fragment overlap counts were separately normalized adjusting for GC-content effects by gbsq-normalization [69] and by dividing raw counts by the min-*Q*_0.9_ range.

Cellular context labels were kept for downstream tasks such as leave-one-cellular-context-out (LOCO) rounds for hyperparamter tuning or evaluating performance separately for each training context (Supplementary Methods Fig. 4).

#### 2.2.8 Labeling

The merged ChIP-seq matrix *B* was used to define the training labels 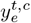 for each PRE site *e* for TF *t* in a training context *c*. A site *e* was considered positive if the corresponding entry of *B* contained an *r*-score greater than 0 in the respective column. PREs that were found to overlap the peak flanks were excluded from the training set. All remaining non positive PREs, i.e. these entries of *B* being 0 in the respective column, were considered as negative instances for the training process of the respective TF.

### 2.3 TFBlearner algorithm

#### 2.3.1 Classification Models

TFBlearner uses Gradient Boosting Decision Trees (GBDT) as implemented by the Light Gradient Boosting Machine (LightGBM) library [50, 73] to train TF-specific classification models. Models are optimized using binary cross-entropy loss, and trees are grown leaf-wise using information gain as the split criterion.

For each TF, four different GBDT models are trained on different subsets of the data. Positive instances in the set of all positives *P*, with |*P* | = *n*_+_, are ranked by their *r*-score in descending order. The first three single classification models *K*_*i*_ in the ensemble 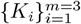, are trained on different overlapping subsets of positives *S*_*i*_ by including positives up to increasing ranks *k*_*i*_ in their respective training sets (see Supplementary Fig. 1). The cutoff rank *k*_*i*_ for model *K*_*i*_ is obtained as follows:

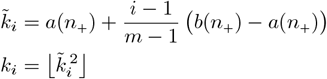

with:

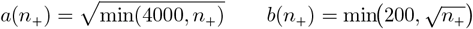

Thus, model *K*_*i*_ is trained on the subset *S*_*i*_ of the top-*k*_*i*_ positives:

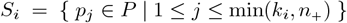

Since *k*_1_ ≤ *k*_2_ ≤ *k*_3_, the sets are nested: *S*_1_ ⊆ *S*_2_ ⊆ *S*_3_ ⊆ *P*. Hence models are trained on sets of positives with different lower-bounds of the *r*-score. Models *K*_1*−*3_ are trained using instance weights based on the *r*-score, whereas a fourth model *K*_4_ is trained on set of the same size as *S*_3_ but with randomly sampled positives and uniform instance weights (see Supplementary Fig. 2).

For all models *K*_1*−*4_, negatives are subsampled in the respective training data such that the positive fraction equals 0.25. Negative instances are partitioned into *clear* and *borderline* negatives: *clear* if the motif matching score is *< Q*_0.4_ or *<* 1, or if the total fragment overlap count is ≤ *Q*_0.3_; *borderline* otherwise. Negatives are then sampled uniformly at random without replacement, drawing approximately 30% from *clear* and 70% from *borderline* instances, with deficits reallocated between categories as available. The selected negatives are combined with the subsets of positives *S*_*i*_ to form the final training sets.

Predictions of all four single models *K*_*i*_ in the ensemble are stacked as weighted average to obtain the final binding prediction for each PRE *e* for a TF *t* in cellular context *c* (see Supplementary Fig. 1 and Supplementary Fig. 2B for a performance comparison using stacking).

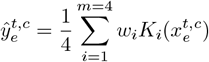

With 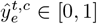 corresponding to an averaged predicted binding score. Per model weights *w* are obtained based on the relative performance of each model on two chromosomes (10,11) that are removed from model fitting (Supplementary Methods A.3.1).

#### 2.3.2 Hyperparameter tuning

Hyperparameters for each model *K*_*i*_ in the TF-specific stacked ensemble are tuned independently using model-based optimization (MBO). The mlr3mbo package from the mlr3-ecosystem is used with Kriging Regression as surrogate model and Augmented Expected Improvement (AEI) as acquisiton function (for further details see Supplementary Methods A.3.2). The hyperparameters - learning rate *η*, number of leaves, feature fraction, bagging frequency, L1/L2 regularization (*λ*_1_, *λ*_2_), and minimum gain to split - are tuned for up to 100 iterations with early stopping (see Supplementary Table A2 and Supplementary Fig. 2). In each iteration, the base learner (GBDT) is trained and evaluated with a given hyperparameter configuration in either leave-one-cellular-context-out (LOCO) or 5-fold cross-validation rounds (if only one cellular context in the training data).

#### 2.3.3 Training

Each model *K*_*i*_ of the TF-specific ensemble is trained separately up to a maximum of 2500 boosting rounds. Early stopping is applied if performance stagnates for 10 boosting rounds. Early stopping and the choice of the best boosting iteration are performed using a fraction of 0.15 of the original training data that is held out from training and reserved as a validation set. Datapoints are sampled at equal proportions from weight bins such that a similar distribution of instance weights is preserved in the validation set and a positive fraction of 0.01 (which is close to the typically observed proportion of sites with TF bindings) is obtained. Performance on the validation data is evaluated by computing AUPRC separately for each cellular context and taking the average across as measure. AUPRC is calculated using the PRROC R-package [74]. Further training was restricted to the autosomes.

Instance weights are used during training: each positive instance receives its *r*-score as initial weight. When training on multiple cellular contexts instance weights are rank-normalized to 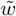 in a two-stage procedure by sorting positive instances by their *r*-score in each context. At first, the top-*k* positive instances of each context are rank-normalized across contexts, where *k* is the smallest absolute number of positives observed in any of the training contexts. Weights across contexts are normalized to the (respective within context) rank by attributing the top-*k* weights observed across all contexts to the top-*k* positive instances of each context. In second step, remaining low rank positives in contexts with surplus positives are separately rank-normalized in reverse order. The minimal-*k* weights observed across contexts are attributed to the minimal-*k* remaining positives of each context with surplus positives. In both stages, ties are broken at random. Negative instances all receive the same instance weight 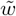 of 1. However, weights are additionally adjusted for class imbalance internally by the lightGBM implementation using `is_unbalance=TRUE`.

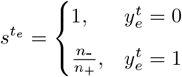

Thus, the final instance weight of each PRE *e* used for training is:

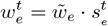

Models are trained within an outer LOCO loop if more than two training cellular contexts are available to estimate performances on unseen cellular contexts (see Supplementary Methods A.3.3). Additionally, chromosomes 2, 4, and 9 are held out from training to estimate performance on these.

#### 2.3.4 Feature Importances

Features importances were computed using LightGBM’s R functions, and averaged across the bagged models. The standard tree-based Frequency (how often the feature is used), Gain (improvement brought by splitting on the feature), and Coverage (number of items affected by such a split), were computed for each TF (made available, see Data availability). However, we chiefly concentrated on the SHAP values: by virtue of being additive, they allow aggregation of features into more interpretable feature groups.

LightGBM uses the TreeSHAP algorithm [75], which uses dynamic programming to efficiently traverse the trees, updating the probability of each feature’s inclusion and weighing its marginal contribution. Specifically, following [76], the weight for a feature *i* is computed as:

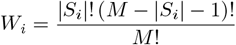

Where *M* is the total number of features, *S*_*i*_ is the set representing the subset features that precede *i*, and |*S*_*i*_| its cardinality (the number of features preceding *i*). The weights represent the fraction of possible feature permutations in which feature *i* appears after *S* in the tree. When leaves are reached, the algorithm distributes their values back to the features according to their weight, and the resulting Shapley values are scaled by the learning rate and summed across trees.

Shapley values are computed individually for each feature, each cellular context used in training, and each region of the chromosomes held-out from training (see Supplementary Methods A.3.3), subsampling negatives (i.e. not bound according to the ground truth) to a maximum of 100’000. To obtain global importance of the features, we sum the absolute Shapley values across these instances, and then average across the submodels, i.e. 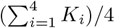. In addition, since the Shapley values represent the feature’s contribution *towards a positive prediction*, we computed the sum of the (signed) Shapley values across ground truth positives to distinguish positive and negative contributions.

To provide more interpretable information from the feature importances, we summed Shapley values by group of features, as indicated in Table 1. We additionally performed other aggregations to address specific questions (in Figure 4). To represent features stemming from ATAC-seq of the target cellular context (as opposed to e.g. associations across contexts), we used features counting overlaps or inserts, as well as based on the insertion weight profile. To represent sequence-based features, we aggregated the importance of the sequence embeddings, the motif counts and matching scores. For the importance of cofactors, we used features from the groups Motif matches, NMF binding patterns, and Cofactor binding signal (single feature selected by `tfCofactor*, selectedMotif*, patternTf*, ctcfBind*` and `coBind*`, see Supplementary Methods Table A4).

Since the Shapley values can be interpreted as log-odds (relative to the base rate), based on the feature, of the item being a positive, it can distinguish positive and negative contributions (to the probability of being a positive). To use this feature in estimating the importance of specific cofactors, we additionally summed the (signed) Shapley values of (ground-truth) positive items across the features relative to each given cofactor.

Finally, since the sums (across predictions) of Shapley values are dependent on the proportion of positives, they are not directly comparable across TFs. We therefore normalized them by expressing them as a percentage of the total Shapley values for the given TF.

### 2.4 Binding site prediction benchmark

#### 2.4.1 Datasets

Matched ATAC- and ChIP-seq datasets of five training cellular contexts (K562, GM12878, HepG2, HeLa-S3, MCF-7) and two testing cellular contexts (H1, Jurkat) for 32 TFs in total were downloaded from ENCODE and GEO (Supplementary Table 3).

Bam-files of ATAC-seq datasets from ENCODE were downloaded merged and for each cellular context (samtools merge v.1.18 [77]). Datasets from GEO were downloaded as fastq files and processed as described in A.1.3. ATAC-seq peaks were called using macs2 [65] (v2.2.9.1) with `callpeak --nomodel` and blacklisted regions were removed.

All ChIP-seq datasets were downloaded from ENCODE, including conservative IDR-thresholded and IDR-thresholded peaks, alongside with the coverage bam-files. ChIP-seq peaks from the Jurkat cell line were lifted over using the `liftOver` function from the rtracklayer [78] R package (v.1.68.0) using the UCSC hg19ToHg38.over.chain file. All input data was restricted to chromosomes 1-22.

These preprocessed ATAC-seq and ChIP-seq files were used as inputs for all methods. Whenever additional preprocessing was required by a specific method, it was carried out starting from these commonly preprocessed files.

#### 2.4.2 Peak Sets

For training, all supervised methods required ChIP-seq peaks as input to define positive instances. A common set of peaks was therefore derived by first merging conservative IDR-thresholded peak files, concatenating overlapping peaks using the `reduce` function from the GenomicRanges [72] R package (v1.60.0). Non-conservative IDR-thresholded peaks were then incorporated into this set through two additional recursive merging steps (see Supplementary Methods A.5.2), in which peak boundaries were defined by the median start and end coordinates of overlapping peaks. IDR-thresholded peaks included in that set were retained only if they overlapped at least one peak from a different ENCODE dataset with the same specifications. These merged peaks were used as positive instances for training of all methods, if a method did allow for defining ambigious negatives (Catchitt, TFBlearner), these were defined by using IDR thresholded peaks not overlapping any of the merged peaks.

For evaluation purposes, several peak sets were defined. Apart from the merged set of ChIP-seq peaks (‘Merged’), all IDR-thresholded peaks were used as positives (‘IDR-thresh.’) and a set of unambigious peaks was defined which included all the merged ChIP-seq peaks as positives and IDR-thresholded peaks not overlapping any of the merged ChIP-seq peaks were marked as ambigious negatives and subsequently removed from the evaluation. Additionally, we defined a set of non-crowded peaks by excluding those peaks falling within the top 1% of C-scores (see Supplementary Methods A.2.6). Similarly, we defined two exploratory peak sets: one consisting of peaks that did not contain a motif match for the corresponding TF (‘No motif’), and another comprising peaks located at sites where no peak was observed in any of the training contexts (‘Not in train’).

#### 2.4.3 Methods

BMO [53] fits negative binomial models of ATAC-seq fragments and the number of co-occurring motifs not using any ChIP-seq data.

The BMO Snakemake workflow (v1.0) was used on the test ATAC-seq data with motif matches obtained by FIMO [79] (detailed below).

FIMO motif matches were obtained using the `runFimo` wrapper-function of the memes package (v.1.16.0) based on meme (v.5.5.4) [80]. Motif matching scores (log likelihoods) were used as a baseline score. On the set of PREs a lenient threshold of 0.01 was used, whereas on the full genome the default of 10^*−*4^ was used. FIMO motif matches were used as an input for other methods (BMO, Catchitt, TFBlearner.basic, TOP).

Catchitt [37] was one of the methods that performed best in the ENCODE-DREAM Challenge 2017. It uses a weighted variant of the discriminative maximum conditional likelihood principle for training TF-specific models based on a diverse set of sequence and chromatin accessibility related features. Here, we used the Catchitt Java implementation (v.0.1.4) as documented on https://jstacs.de/index.php/Catchitt, using motif matching scores from FIMO and the processed ATAC-seq bam files. As the documentation does not provide clear instructions on how to train Catchitt based on ATAC-seq data from multiple cellular contexts, we used Catchitts preprocessing scripts (`access, motif, labels`) on the input data and subsequently concatenated data of the training cellular contexts into one file. Models were trained with default settings.

The maxATAC [43] algorithm is based on a deep convolutional neural network architecture. It takes as input the one-hot encoded DNA-sequence together with smoothed and read-depth normalized Tn5 cut sites obtained from an ATAC-seq fragment file. TF-specific models are trained using both ChIP-seq read files bigwig (bw) before peak calling and peak coordinates. TF bindings are predicted in a 32bp resolution. We used the maxATAC python implementation (v1.0.6) as recommended in the documentation. Input datasets were preprocessed using maxATAC functionality (`prepare, average, normalize`). The model was once trained with default settings of 20 epochs (maxATAC.default) and with 100 epochs (maxATAC.100) as in the original publication. MaxATAC.default was trained using CPUs for direct comparison with other methods, while maxATAC.100 was trained on a GPU (NVIDIA L4, 24 GB).

TOP [54] is a Bayesian hierarchical model that predicts bindings based on Tn5 insertions quantified in the vicinity of motif matches. The hierarchical approach models TF binding in three levels (TF-generic, TF-specific cellular context-generic, TF-specific and cellular context-specific). Accordingly, the TOP model was trained on all benchmark TFs together using the TOP R-package (v.1.0.1). Data was preprocessed using TOP functions for counting and normalizing Tn5 insertions at motif matches (`process_candidate_sites, count_genome_cuts, get_sites_counts, normalize_bin_transform_counts`). TOP’s logistic mode was used to obtain binary predictions of TF binding.

The TFBlearner features were constructed as previously specified based on two different levels of data support. Features were once constructed solely on the benchmark data sets (TFBlearner.basic). However, as many features (e.g., Association, Binding patterns) depend on sufficiently diverse data, we additionally derived features from the pre-compiled motif match, association, and ChIP-seq matrix with all columns corresponding to TFs in the benchmark datasets removed (TFBlearner.full). Both models were otherwise trained in the same way as previously described on the benchmark datasets.

### 2.5 Differential TF activity inference

#### 2.5.1 Relative TF activity inference from ATAC-seq

For TF activity inference from ATAC-seq data, we used the count matrices from [17]. The peaks were overlapped with PREs, and the maximum prediction score per peak and TF was used. The motifs (from the motif aggregation in section A.1.8) of the corresponding TFs were used for the motif-based equivalent. We noticed that the motif-based approach failed to identify the NR3C1 intervention, which was instead accurately identified in [17], and upon closer inspection noticed that the MOBI-reported motif for NR3C1 did not match the classical glucocorticoid response element. We therefore included the correct motif from HOCOMOCO, which enabled the correct identification of the perturbation. The metrics used for evaluation are described in [17].

#### 2.5.2 TF activity inference from RNA-seq

To link regulatory elements to putative target genes, we took ENCODE’s rE2g predictions [81], specifically the files ENCFF040PZI, ENCFF095CQE, ENCFF328COH, ENCFF639BAS, and ENCFF680XTP. Scores for each enhancer-gene were averaged across samples, before mapping to PREs and taking the maximum score per PRE. Proximal links were added to these region-gene links by assigning a score of 1 to the region 2.5 kb upstream the gene’s TSS, and assigning a score of 0.75 to regions between 2.5 and 5 kb upstream or between 500 bp and 1500 bp downstream. The region immediately downstream the TSS was ignored, as it is generally associated with transcriptional repression [82]. Motif matches were obtained for the same set of PREs.

The perturbation transcriptomic data from [59] was downloaded from https://doi.org/10.5281/zenodo.8071808 (version 9), published with [83]. Genes with a mean logCPM of 1 or above were retained. Only perturbations with replicates, with a Pearson correlation of logFCs between replicates of 0.3 or more and with at least 20 differentially-expressed genes (DEGs) were retained. The collecTRI regulons were fetched on 2025.12.15 using the *get_collectri* function of the decoupleR package, splitting complexes. For the collecTRI regulons, it is necessary to remove identical regulons to avoid colinearity errors. We did so by preferentially retaining perturbed TFs and thus avoiding the loss of any relevant TF. Only the regulons of TFs among the perturbation or among expressed genes were retained. TF activity inference was done using the fastMLM function from [17], assigning the *t*-scores of a multivariate regression of the expression (logCPM or log2FC) on the regulons. This is simply a faster equivalent of the decoupleR `mlm` function.

### 2.6 Crowdedness

For our characterization of crowdedness across cell types, we first computed a crowdedness score in a cell type-specific fashion as the proportion of profiled TFs (i.e. TFs for which ChIP data in that cell type is available) which have a peak with a *r*-score above 0.1 whose center overlaps the region. To avoid crowdedness being driven by the region’s width, this was then normalized to a 200bp width. Specifically, crowdedness 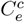 for a region *e* in cell type *c* is defined as:

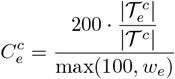

where 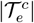 is the number of TFs bound to the region *e* in cell type *c*, |*T*^*c*^| is the total number of TFs profiled in cell type *c*, and *w*_*e*_ is the width of region *e*.

A region was classified as highly-crowded if 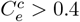, i.e. if at least 40% of all unique TFs with available ChIP-seq data in that specific context showed a peak within that region (adjusting for width).

## 3 Discussion

We developed a method and pipeline to train TF-specific models predicting binding to DNA in a cell type-specific manner, requiring only ATAC-seq data of the target cell type. A pre-registered benchmark on held-out cell types shows that our method, TFBlearner, is either superior to or on par with the best alternative method across a wide variety of evaluation metrics, while being less computationally-intensive than deep learning methods. This was achieved through a number of innovations, including pre-defined genomic windows capturing *>*99.8% of peaks and enabling matrix representation of the data, stacked learners, observational weights based on peak replicability, and model-based hyperparameter optimization enabling appropriate regularization and hence improved generalizability across cellular contexts. In addition, rather than employing a brute force approach based on sequence and ATAC profile alone, we relied on information sharing across cellular contexts and cofactors, as well as extensive, biologically-inspired feature engineering. This being said, it has to be pointed out that maxATAC also achieved a very good performance on a large fraction of TFs while using less information and background knowledge, and more computational power, and that its predictions have a higher genomic resolution. Importantly however, our modeling approach and the importance the models assigned to these features could recover key biological features of the TFs, and we describe how model complexity and feature importance varies across TF families.

Our streamlined platform has enabled us to train and predict across an unprecedented scale. We trained models for 1108 CAPs, representing the majority of all human TFs, and an order of magnitude more than previous efforts [43]. We then systematically predicted their binding across 43 human cell types, covering all important human lineages and most widely-used cell lines, with a special focus on brain cell types, which although biomedically-relevant are as yet poorly characterized in terms of TF binding. All models and predictions (totally 200 billion single predictions) are openly accessible, and browsable on the TFBPlatform.

The scale of our study enabled a number of observations relevant to the field. First, as the main determinant of the area under the precision-recall curve is, not unexpectedly, the fraction of positives in the testing data, we recommend using the enrichment over expectation as a more robust performance metric. Once this is accounted for, the most important determinant of performance is the extent to which peaks in held-out contexts were seen in at least some training context. In other words, while prediction methods (not only TFBlearner, but all supervised methods benchmarked) are excellent at predicting whether a peak seen in training will be bound or not in the testing context, they all perform very poorly at predicting entirely unseen peaks (Supplementary Fig. 9). While maxATAC was slightly better than TFBlearner in these specific cases, both were inferior to simply using nucleosome-free ATAC signal alone. While some TFs (e.g. forkhead/winged helix TFs) were nonetheless highly generalizable, this indicates that all models fail at finding, for most TFs, predictive patterns that are truly generalizable across diverse contexts. This being said, the proportion of all peaks observed increases with the number of training contexts, such that predictions can nevertheless become quite accurate.

Having predictions for the majority of TFs enables downstream tasks, such as the inference of differentially-active TFs based on ATAC-seq or, notoriously more difficult, on the transcriptome. We demonstrate how the binding predictions can be used to recover, in a fully unsupervised fashion, differentially-active TFs from transcriptomic profiles, comparing favorably with the curated collecTRI regulons. Surprisingly, however, the binding predictions did not provide significant gains on activity inference based on DNA accessibility when compared to motifs, despite the latter being over an order of magnitude less predictive of binding. A plausible explanation for this is that much of TF binding – and in particular binding that is easy to predict – is not specific to the TF, i.e. that various TFs tend to bind to the same places. For the purpose of reverse-engineering perturbed TFs, such unspecific binding is not very informative, while motifs are in contrast more specific to the TF (or at least to the TF family). Instead, the predictions did improve inference from the transcriptome. We speculate that this is because, contrarily to ATAC-seq, the transcriptome does not provide information as to the relative activity of different regulatory elements, which might be embedded in the binding predictions.

The importance of TF-unspecific binding is also underscored by the surprisingly high performance of the TF- and context-agnostic C-score in predicting TF binding (Fig. 2 and Fig. 3H). Most TFs, it would seem, tend to bind to regions where other TFs bind. To further understand this phenomenon, we studied crowdedness (the density of CAPs binding a given region) across five cell types for which a large amount of ChIP-seq data was available. We found that crowdedness was associated with accessibility, although not all hyper-accessible regions were crowded. Focusing especially on ‘highly-crowded’ regions, we confirmed, across cell types, previous reports that highly-crowded or HOT regions were enriched for promoters and super-enhancers, and their enrichment for RNA processing genes. Despite the high reproducibility of these enrichments across cell types, the highly-crowded regions themselves were not for the most part shared across cell types. We additionally provided evidence towards the functionality of crowded bindings, both in terms of their dynamic regulation and association with transcription in response to stimuli, consistent with earlier reports suggesting functionality [84]. Finally, we showed that, across all cell types, crowded promoters engage in 3D contacts with more, and on average more crowded distal regulatory elements, and in particular with other promoters. This would be consistent with their common recruitment in transcription factories or condensates [85]. Finally, most bindings at crowded promoters are also found in some of their distal contacts, suggesting that crowdedness could be explained by tethered (or diffusing) bindings enabled through 3D interactions.

Together, these findings support a wolf-pack or two-step model of TF binding [26], where complex protein-protein interactions, especially in high-accessibility regions, determine a large fraction of binding events for many, if not most TFs. Whether the cells function despite this promiscuity, or rather contributes to the cells’ function, for instance by ensuring robust expression of necessary genes across a range of variations in TF activities, remains to be determined.

## Data availability

All data (processed matched ATAC- and ChIP-seq data and motif matching scores), insertion weight models, predictions and per-TF feature importance metrics are deposited on Zenodo: https://doi.org/10.5281/zenodo.18198234. Predictions and feature importance metrics can also be accessed and visualized on a dedicated webpage (TFBPlatform): http://www.ethz-ins.org/TFBPlatform/

## Code availability

The TFBlearner R package is available from: https://github.com/ETHZ-INS/TFBlearner. All code used for training, predictions and analyses presented in this manuscript is accessible via https://github.com/ETHZ-INS/TFB-analysis, and code used for the benchmark at https://github.com/ETHZ-INS/TFB_Prediction_Benchmark. The pre-registered protocol is available at https://osf.io/kv2tw/overview.

## Authors’ contributions

*Emanuel Sonder* Conceptualization, Data curation, Formal analysis, Investigation, Methodology, Writing – original draft

*İpek Güneş Aymergen* Methodology

*Jieran Sun* Investigation, Methodology

*Audran Feuvrier* Investigation

*Johannes Bohacek* Funding acquisition

*Katharina Gapp* Funding acquisition

*Gerhard Schratt* Supervision

*Mark D. Robinson* Supervision, Funding acquisition

*Pierre-Luc Germain* Conceptualization, Data curation, Formal analysis, Investigation, Methodology, Project administration, Supervision, Writing – original draft, Funding acquisition

## Appendix A Supplementary Methods

### A.1 Data acquisition and processing

#### A.1.1 Download and matching of ENCODE data

To download data from the ENCODE project, we first searched its database (on 2025.04.17) for GRCh38 ChIP and ATAC data (see repository for the exact queries). We attempted to match ‘cellular contexts’, i.e. combinations of cell type and condition/treatment, across modalities. When the ‘biosample summary’ text matched between ATAC and ChIP data, we considered it a context match. If not, we first removed age and sex of the donor to gain additional matches. If that still did not give a match and we did not have (from any of our sources) other ChIP data for the TF in question, we tried finding a match omitting genetic modifications. We additionally downloaded and processed ATAC-seq dataset for H1 (GEO IDs GSM8260976 and GSM8260977) and HEK293 (GSM3905874, GSM3905875, GSM4133299, GSM4133300, GSM2902624, GSM2902625, GSM2902626, GSM2902627, GSM5743943, GSM5743942), and manually matched them to ENCODE ChIP-seq data. For ATAC data, we then downloaded the aligned reads. For ChIP data, we downloaded the (per-replicate) peaks, using the following filetype priority: IDR thresholded peaks, optimal IDS thresholded peaks, pseudoreplicated peaks, conservative IDR thresholded peaks, peaks.

#### A.1.2 Download of ATAC and ChIP data from other sources

ChIP-seq data were additionally downloaded from GTRD v20.06 on 2021.07.20. Matching ATAC-seq data from GTRD were reprocessed from raw reads downloaded from the SRA. Codebook [86] ChIP-seq peaks were downloaded (on 2024.11.19) from https://codebook.ccbr.utoronto.ca/index_v1.php, treating the different sets (i.e. GrecoBit, Toronto and McGill) as replicates. brainTF [46] peaks were downloaded from https://github.com/aanderson54/Loupe_BrainTF (commit 458d8a2). For brainTF, only the OLIG2+ and NeuN+ fractions from the DLPFC, FP and OL samples were used, and the IPs from the different regions were treated as replicates. Corresponding ATAC-seq data profiles were generated by aggregating fragments of the relevant cell types from the [87] single-cell ATAC-seq atlas. Specifically, Oligodendrocytes and their progenitors were used to match the OLIG2+ cells, while to approximate NeuN+ cells, neurons population excluding highly region-specific ones were aggregated (i.e. LAMP, MSN, PVALB, SNCG, SST, VIP, L6B, CHO, SUB and IT-neurons).

#### A.1.3 ATAC-seq processing

ATAC-seq data processed manually was first trimmed with trimmomatic v.0.39 [88] before being aligned with bowtie2 v.2.4.4 [89] with `--dovetail --no-mixed --no-discordant -I 15 -X 2000`. Duplicates were marked with Picard MarkDuplicates v.2.27.5.

#### A.1.4 Merging and splitting regions

To obtain a consensus set of non-overlapping regions from multiple sets of regions (e.g. from different cell types, or different motifs), we first used the `reduceWithResplit` function, available in our repository. The function merges the different sets of regions (similarly to `GenomicRanges::reduce`), but handles loci with overlapping regions above a certain width in a special fashion. Briefly, the pre-merging regions are used to compute a coverage across the merged region (i.e. number of pre-merging regions overlapping each position), and the algorithm attempts to identify ‘trough(s)’ within the region’s coverage profile. By default, such a trough needs to have a depth, relative to the maximum coverage on either side, of at least 2 and at least 1/4 of the maximum coverage, and should not lead to split regions of a width lower than 100. Finally, the `resizeToNonOverlapping` function was used to simultaneously extends smaller regions bidirectionally to a size of 100bp or until the next region is reached.

#### A.1.5 Matching of contexts and data between sources

As a first step, the annotations of the collected experiments were manually cleaned to remove inconsistencies. For ATAC-seq data, the level of specification considered for the following was the cellular context annotation. Analogously, for ChIP-seq data, the relevant specification level was defined as the combination of the cellular context and the target TF. For each individual DB source containing both ATAC-seq and ChIP-seq data, the two modalities were subsequently matched based on the cleaned cellular context specification.

Subsequently, experiments from all sources were pooled. Because the ENCODE and GTRD collections shared many datasets, as GTRD also incorporates ENCODE datasets, we prioritized ENCODE data. Consequently, GTRD ChIP-seq datasets that were also available in ENCODE were excluded from the pooled collection together with their matched ATAC-seq datasets. Correspondence between ChIP-seq datasets in the two databases was established using the ENCODE Dataset ID, which was accessible through the external metadata provided by GTRD. Annotations of the matched and pooled datasets were additionally once manually cleaned. A unique ID for each cellular context was given to each ATAC-seq and ChIP-seq dataset in the pooled collection. For ChIP-seq an additional combination ID composed of the gene symbol of the TF and the cellular context ID was attributed. All matched and pooled ATAC-seq and ChIP-seq datasets are listed in Supplementary Table 1.

#### A.1.6 ChIP-peaks merging and labeling

Whenever the same ChIP-seq experiments were available from ENCODE and GTRD, the ENCODE source was preferred and the GTRD ChIP-seq datasets with their matched ATAC-seq data were removed from the pooled collection of experiments.

A considerable fraction of ChIP-seq experiments retrieved from GTRD were available with peak calls of several peak callers (MACS2, SISSR, GEM, PICS). Whenever a specification was only available from the GTRD DB, only the maximum *q*-value from peaks called by MACS2, SISSR or GEM overlapping a PRE was retained. In case a peak was called only by PICS but no other peak caller applied, the PREs overlapping said peak were marked as ambiguous and subsequently removed from the training labels/data of the respective TF. The rational for removing these peaks was based on the low replicability observed for peaks only called by PICS (see Supplementary Methods Figure 2).

**Supplementary Methods Fig. 1.**
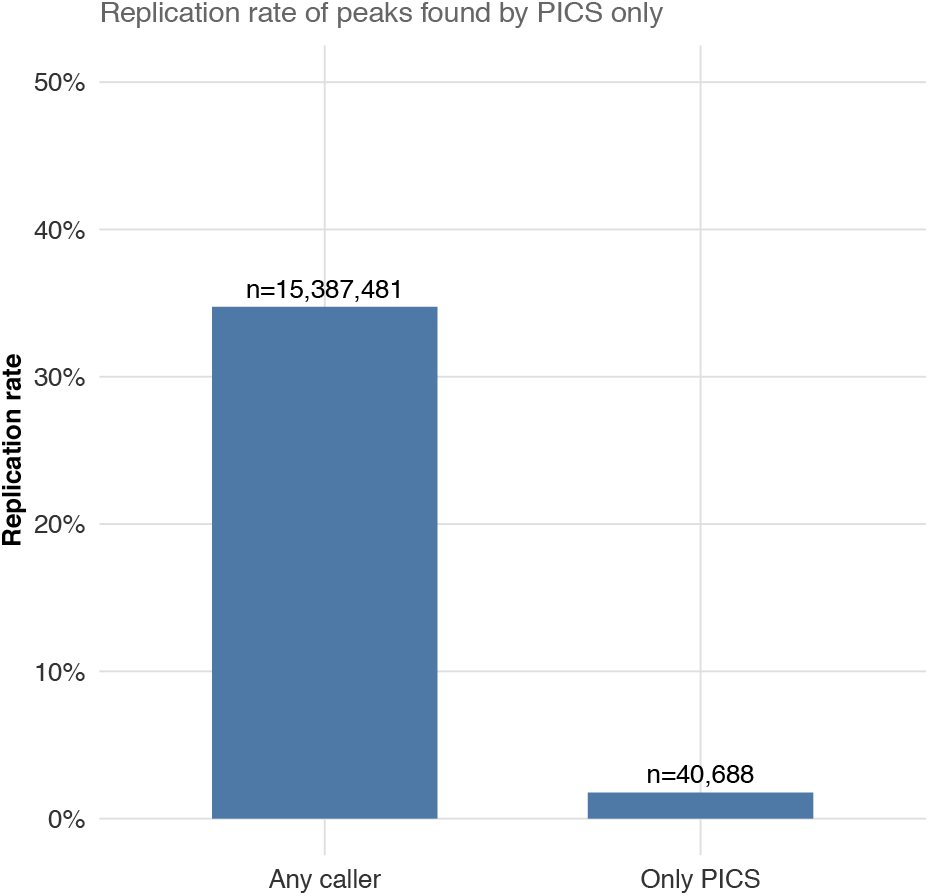
Replication rate of peaks: A peak was considered replicated if it was found to overlap the same PRE in at least two replicates from any ChIP-seq experiments with the same specifications. Peaks shown in the left bar were detected by at least one caller other than PICS, while those in the right bar were detected exclusively by PICS. Shown are peaks from all specifications with at least two replicates across experiments. Peaks were downsampled so that at most 10^4^ peaks were kept for each specification. This data was also used for fitting the GAM.

ChIP-seq experiments designated for training were grouped by their specification, i.e. the TF *t* and cellular context *c*. To merge ChIP-seq experiments with the same specification, peaks of each replicate in each experiment were overlapped separately with the set of PREs E. Peaks were regarded as overlapping if a ± 30bp window around their center — when available, otherwise around their midpoint — intersected with the PRE. For each PRE, we retained the highest peak-calling *q*-value observed among peaks that overlap it across all replicates of all experiments with the same specification. All the merged ChIP-seq specifications and their matched ATAC-seq data used subsequently for training are listed in Supplementary Table 2.

**Supplementary Methods Fig. 2.**
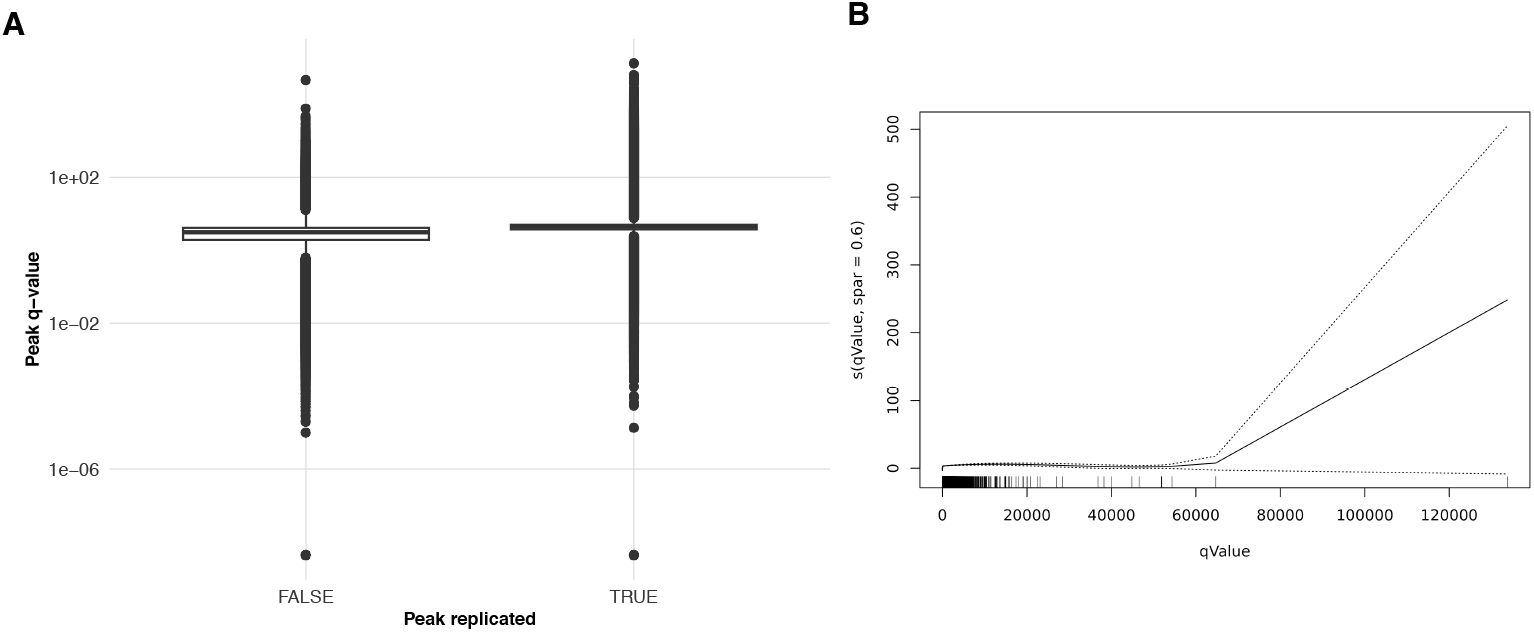
R-score. **A:** Boxplot of q-values of peaks depending on if they have found to be replicated. **B**: GAM smoothing spline used to transform the q-value into the *r*-score (see Methods A.1.6).

#### A.1.7 Inferred replicability score (r-score)

All specifications with at least two replicates across experiments in the training datasets were used to compute a replicability probability (*r*-score) for each PRE with an overlapping peak. In case a PRE was found to overlap peaks from at least two replicates from any experiment, the peak at this PRE was considered to be replicated. A logistic Generalized Additive Model (GAM) (binomial family with a logit link) was fitted using smoothing splines (`s(.,spar=0.6)`) to model whether a peak is replicated as a function of its merged *q*-value, employing the gam package [90] in R. The *r*-score was then computed for all specifications with the later GAM for each PRE based on the merged (maximum) *q*-value. In case of at least two different replicates with peaks overlapping a PRE, the *r*-score was set to 1. Furthermore, if none of the replicates showed a peak whose center window overlapped with the PRE, but at least one peak overlapped only with its flank (i.e., the portion of the peak outside the ±30bp margin around its center), the corresponding PRE was marked as a flanking for the respective TF.

The resulting ChIP-seq matrix has the form *B* ∈ [0, 1]^| *E*|*×*|*S*|^, where *S* = ∈ *T* × *C* denotes the unique specifications, i.e. TF-cellular context pairs, for which ChIP-seq data was collected. Each entry represents the maximum *r*-score of an overlapping peak at a PRE for a particular specification, or is assigned a value of 0 if no peak was observed.

#### A.1.8 Motif aggregation and scanning

To obtain one primary motif per TF gene (from [25]), we aggregated motifs from multiple sources, with the following decreasing order of priority: Codebook [45], MOBI primary motifs [32], the best HOCOMOCO v12 Core motif [56], MotifDb v. 1.50.0 [91], and then either (for ENCODE peaks) de-novo discovered motifs (see below) or, for others, cisTarget v10 clustered motifs [92]. For the latter, the best 5 motifs per gene were merged.

In addition to these core set of motifs, we included dimer motifs from [93] and [94], as well as core dinucleotide motifs from HOCOMOCO v11 [95]. Dinucleotide motif scans were done using sarus 2.1.0 (https://github.com/autosome-ru/sarus), keeping the best score per PRE. All other motif scans were done with motifmatchr v1.1.1 [96], considering the best score per PRE with p-value lower than 0.001. For motif counts, matches with the default cutoff (5e-05) were used.

#### A.1.9 ChIA-PET data

The distal regulatory elements and their links to genes based on ChIA-PET data were downloaded on Nov 13, 2025 from the ENCODE Screen (v4) website on https://downloads.wenglab.org/Human-Gene-Links.zip (file V4-hg38.Gene-Links.3D-Chromatin.txt). The ENCODE Screen regions were then mapped to our PREs.

#### A.1.10 GR data

For the full transcriptome, we downloaded the transcript quantifications from the ENCODE platform and aggregated the counts to the level of gene symbols. To obtain unprocessed RNA, we downloaded the filtered bam files from the ENCODE platform. We first used featureCounts v.2.0.3 [97] with `-t transcript --nonSplitOnly --largestOverlap --nonOverlap 3 --fracOverlap 0.9 --primary -s 0` to obtain all unspliced counts, and subtracted unspliced counts entirely contained within an exon (and hence ambiguous), obtained with `-t exon --nonSplitOnly --largestOverlap --nonOverlap 3 --fracOverlap 0.9 --primary -s 0`. For both types of quantifications, we filtered expressed genes using `edgeR::filterByExpr with min.count=20` [98]. Since there is a very strong technical difference between two batches of samples, we used the ComBat empirical Bayes method, as implemented in the sva R package [99], on the log(CPM) values to correct this, and computed log2(fold-changes) based on these corrected expression values. Finally, we considered a gene bound if it had a GR binding site within 5kb upstream of (any of) its TSS.

For the ChIP-seq and ATAC-seq data, we downloaded filtered bam and thresholded peak files from the ENCODE platform (for ATAC-seq, we used accessibility at baseline, i.e. time-point 0). We merged peaks (of the same modality) using `GenomicRanges::reduce`, keeping only regions found in more than one sample. We then counted fragments overlapping each set of peaks. For the ChIP-seq data, where standard normalization methods assuming an absence of global difference would be inappropriate, we used background normalization, as implemented in the `epiwraps` package v0.99.108 (https://github.com/ETHZ-INS/epiwraps).

The complete list of ENCODE file accession IDs used are included in Supplementary Table 1.

#### A.1.11 Cofactor data

To identify cofactors, we downloaded human interactions from the stringdb v12 [100], keeping only (loosely-defined) TFs and used only interactions with experimental support or with a combined score above 200. We then selected the top 10 interactions with experimental or database support, followed by the next top 10 interactions by combined score.

### A.2 Feature construction

#### A.2.1 Selection of Associated Motifs

Motifs that are observed to co-occur or be mutually exclusive with the ChIP-seq peaks of TF *t* in the training dataset are selected during feature construction. The ChIP-seq peak labels of *t* are aggregated into a binary vector by marking a site as positive if a peak (*r*-score*>*0) is found in any of the training contexts. The resulting label vector *ℓ* ∈ {0, 1}^|*E*|^ and the motif matching score matrix *M* ∈ ℝ^|*E*|*×m*^ are both subsampled by the same randomly drawn 2*e*5 sites *R*.

For each motif *j* {1, …, *m*}, a per-motif threshold is defined based on the maximum matching score observed.

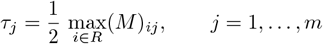

Based on *τ*_*j*_ a motif-occurrence indicator matrix *C* ∈ {0, 1}^|*R*|*×m*^ and an motif-absence indicator matrix *E* ∈ {0, 1}^|*R*|*×m*^ are obtained by:

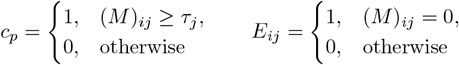

A Jaccard index is computed between *ℓ* and for each column of *C* and *E*.

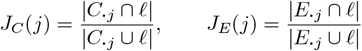

Co-occurring motifs are obtained by ranking *J*_*C*_(*j*) in descending order and selecting the top *k* = 5. Analogously, ranking *J*_*E*_(*j*) yields the top mutually exclusive motifs.

#### A.2.2 Sequence Embedding

To capture sequence regulatory grammar, we created a co-embedded of sequence and (cell type-agnostic) bindings using StarSpace [68], similarly to [101]. We created two embeddings of 10 dimensions each: a first focused on the local 300bp, and another focused on the broader region of 1024bp. In each case, we first identified all matches for motif archetypes [67] in each sequence, keeping only the strongest matches for motifs that overlapped by more than 4 bp. We then translated each sequence into an ordered set of 8-mers around the center of motif matches, representing the left-hand-side features. We then randomly selected a million regions among those that had, across our datasets, peaks for more than 2 TFs. We assigned these regions with right-hand-side labels of the TFs with peaks with a replicability score of at least 20%. Naturally, data from the benchmark’s testing cell types was entirely excluded. We then ran StarSpace, with `-trainMode 0 -minCount 15 -dim 10 -lr 0.1 -maxNegSamples 20 -epoch 100`. Based on the model, we generate embeddings for all PREs, and joined the local and broad embeddings.

#### A.2.3 Insertion Weight Profiles

TF footprints, represented as insertion weight profiles, are computed by counting Tn5 insertions at positions relative to the motif matches of a TF *t*. Insertion positions are obtained by shifting the ATAC-seq fragment end coordinates on both sides inwards by an offset of 4bp to account for the Tn5 insertion bias.

Insertion weight profiles were pre-computed using the matched ATAC-seq and ChIP-seq datasets from the training cellular contexts. We then selected occurrences of the canonical motif of *t* within PREs that overlap ChIP-seq peaks with an *r*-score *>* 0.5 in the training cellular contexts. Across these motif matches all insertion events within ±200bp margin of the motif match center were summed at positions *p* ∈ *P*, with *P* = {−200, …, 0, …, 200}, relative to the motif center (*p* = 0).

The resulting counts *c*_*p*_ were adjusted at positions lying within the motif match of length *l* by averaging the observed count with the median count across all positions in the match:

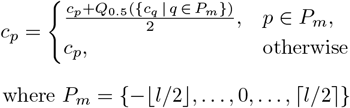

This procedure was used to reflect the greater uncertainty in the counts measured at positions that may be shielded from Tn5 insertions by the occupant TF.

The per TF insertion position weights were obtained by scaling the positional counts by the maximum count and normalizing weights to unit sum:

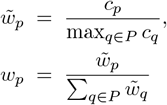

Insertion profile weights were calculated independently for each training cellular context. The median value of each positional weight across all training contexts was then taken to generate a final insertion weight profile used for feature construction (see Supplementary Methods Fig. 3). All pre-computed insertion weight profiles are deposited under: 10.5281/zenodo.18198234

**Supplementary Methods Fig. 3.**
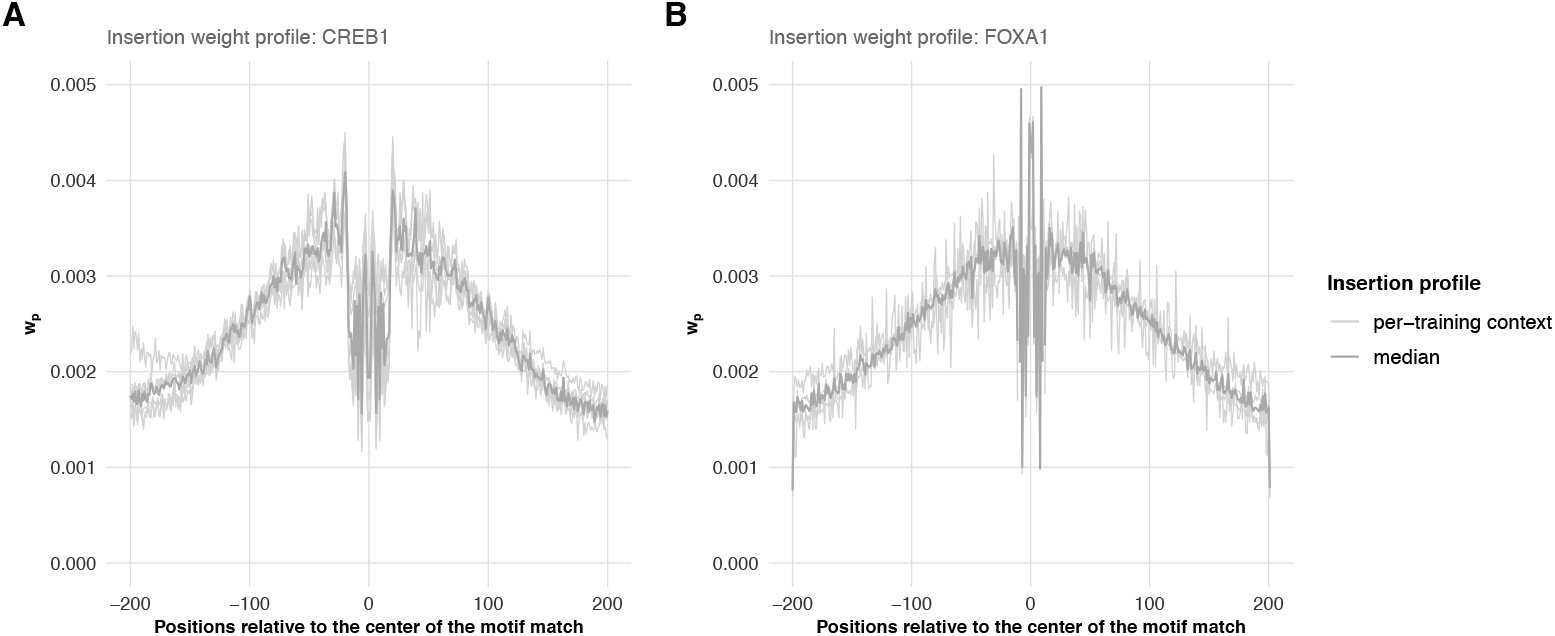
Insertion weight profiles: Per-training context computed profiles are indicated by lightgrey lines whereas the median profile computed from the per-context profiles is shown in darkgrey. Examplified for two TFs CREB1 (**A**) and FOXA1 (**B**).

For each PRE site *e* with a motif match, we also included as a feature the deviation of the observed insertion events at that site from the expected insertion weight profile.

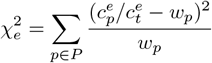

where 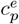 is the observed insertion counts at position *p* and 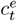 the total observed insertion counts in the ±200bp window around the motif match center overlapping site *e*.

#### A.2.4 chromVAR: Motif-activity estimation

The chromVAR algorithm [58] calculates how much the total fragment overlap counts in peaks that contain a given motif match differ from the expected count, where the expectation is derived from the average across all samples. It then evaluates these deviations by comparing them to those obtained from random background peak sets that share similar GC content and average accessibility with the motif-containing peaks. Here, we took the collected ATAC-seq matrix *A* ∈ ℝ^|*E*|*×*|*C*|^ and removed all sites whose mean accessibility was higher than the *Q*_0.9_ or lower than the *Q*_0.1_ of the mean accessibility distribution. Subsequently, the matrix was further downsampled to 5*e*10 rows. Deviations were computed by the chromVAR R package for each motif and each cellular context with ATAC-seq data. The resulting deviations were subsequently centered and scaled to unit variance as recommended in [17].

#### A.2.5 Binding Patterns

The ChIP-seq peak matrix has the form *B* ∈ [0, 1]^|*E*|*×*|*S*|^, where *S* := (*t, c*) ∈ *T* × *C* denotes the specifications, i.e. TF-cellular context pairs, for which ChIP-seq data was collected. Each entry represents the maximum *r*-score of an overlapping peak at a PRE for a particular specification, or is assigned a value of 0 if no peak is observed. For each TF, non-negative matrix factorization (NMF) is applied to the ChIP-seq matrix *B* after removing all columns corresponding to the TF of interest *t*_*i*_ or testing cellular contexts. Specifically, the matrix is aggregated in two distinct ways: (1) across TFs, by taking, for each TF at a site *e*, the maximum *r*-score observed across all its training cellular contexts, and (2) across TFs within each context, by taking, for a given site *e* in a context, the maximum *r*-score over all TFs. Both the per-TF aggregated matrix 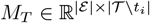 and the per-context aggregated matrix 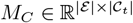 are factorized using an L1-regularized variant of NMF, based on alternating least squares (RcppML [70] R package). The factorization into lower-rank (*k*) matrices is of the form:

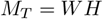

With a feature matrix 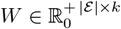 and coefficient matrix 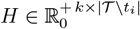. The columns of feature matrix *W* represent distinct binding patterns shared by (clusters of) TFs as indicated by *H*. Columns of *W* can also be understood as clustering centroids of TFs clustered by their observed bindings (presence of peaks at PREs), whereas *H* expresses the cluster membership of each TF. The original ChIP-seq matrix has been aggregated to a) lower the bias for TFs with more cellular contexts covered by ChIP-seq and b) to favor general binding patterns (of clusters) of TFs found to co-bind at the similar positions across contexts. A L1-regularization penalty of 0.5 has been used for both *W* and *H*, and a rank of *k* = 40 has been selected. Analogously, the per-context aggregated matrix *M*_*C*_ was decomposed. However, here with the aim to find patterns of shared occupancy by any TF across contexts.

The columns of both feature matrices *W* were subsequently used as binding pattern features for each site *e*.

#### A.2.6 C-score

The C-score, or crowdedness score, was first introduced by [32], and in model training and benchmark we use it in a cell type-agnostic fashion. Specifically, the C-score as a characteristic of a site *e* and used as a feature is computed as the sum of the per-TF mean *r*-scores at that site. Per-TF means at a site were computed across all the training cellular contexts with ChIP-seq data for that TF. In the benchmark, crowded regions were defined as regions in the top 1% percentile of the distribution of the number of unique TFs with a peak, analogous to previously-described hot regions [30].

### A.3 Modeling

#### A.3.1 GBDT Model

Single GBDT classification models *K*_*i*_ from the TF-specific ensemble 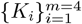 are trained using the Light-GBM R package [73]. Trees are grown leaf-wise and gradients are computed based on subsets of the data using Gradient-based One-Side Sampling as specific to and implemented by the LightGBM framework. Binding probability predictions of model *K*_*i*_ are obtained by including trees up to the best iteration, which is determined on an internal validation set (see Methods 2.3.3).

The final predicted binding probability is obtained by a weighted average of the predictions of models *K*_1*−*4_:

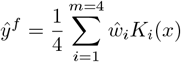

For each model, the weight 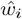 is determined by its relative performance on chromosomes 10 and 11, which are excluded from the model fitting. The AUPRC of model *K*_*i*_ is divided by the sum of the AUPRCs of all four models to obtain its weight.

#### A.3.2 Hyperparameter tuning

Hyperparameters for each model *K*_*i*_ in the TF-specific ensemble are tuned independently using model-based optimization (MBO). MBO, as described in great detail in [102], is based on a surrogate model *f*_*s*_ : *X* → ℝ that predicts the performance *y* of the base learner (here, a GBDT) as a function of the hyperparameter configuration *x* ∈ *X*, with *X* being the explored hyperparameter space (see Table A2). At each iteration, an acquisition function *f*_*a*_ : X → ℝ proposes candidate hyperparameter configurations to evaluate. The base learner is trained using the proposed hyperparameters *x*, evaluated to obtain *y* and the observation (*x, y*) is added to the data. Subsequently, the next iteration starts with *f*_*s*_ being refit on the updated data. The procedure starts with an initial design phase that evaluates a diverse set of configurations to fit the initial surrogate model. After the initial design phase, the loop function directs the iterative process until a termination criterion is met. In each iteration, the base learner is trained and evaluated with a given hyperparameter configuration *x* in either leave-one-cellular-context-out (LOCO) or 5-fold cross-validation rounds (for a single cellular context in the training data) to estimate *y*. Sensible defaults from the mlr3mbo package were used for the main MBO components, exact specifications are listed in Table A1.

**Table A1:** MBO building blocks used. An AUPRC implementation from the PRROC [74] R package has been used to compute performance.

| MBO Component | Method | mlr3-key |
| --- | --- | --- |
| Loop function (defines also the initial design) | Efficient Global Optimization | `bayesopt_ego` |
| Surrogate function | Kriging regression | `LearnerRegrKM` |
| Acquisition function | Augmented Expected Improvement | `aei` |
| Acquisition optimizer | Random Search | `random_search` |
| Performance measure | AUPRC | - |
| Base learner | GDBT (lightGBM) | `LearnerClassifLightGBM` |
| Termination criterion | max of 100 iterations or stagnation for 10 iterations (threshold=0.001) | `combo` |

**Table A2:**
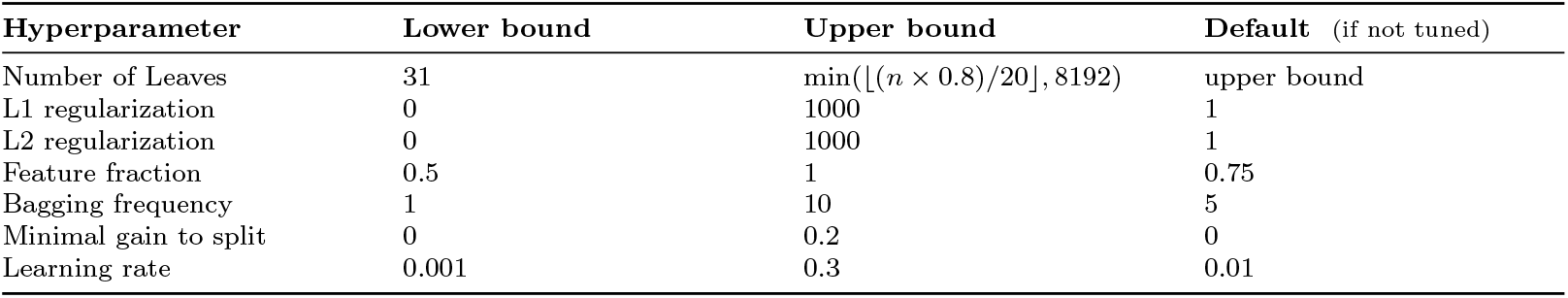
Hyperparameter space *X* explored for each model *K*_*i*_. The upper bound for the number of leaves is computed based on the number of training data points *n* used for the respective model.

**Table A3:** Packages of the mlr3-ecoystem used.

| Package | Version |
| --- | --- |
| `mlr3` [103] | 1.0.0 |
| `mlr3misc` [104] | 0.18.0 |
| `mlr3mbo` [51] | 0.3.0 |
| `mlr3tuning` [105] | 1.4.0 |
| `mlr3learners` [106] | 0.12.0 |
| `mlr3extralearners` [107] | 1.0.0 |
| `mlr3measures` [108] | 1.0.0 |
| `paradox` [109] | 1.0.1 |

#### A.3.3 Training and performance evaluation

Each TF-specific ensemble model was trained on the row-bound feature matrix *D* with three chromosomes (2,4,9) removed and held-out for performance evaluation using AUPRC. Additionally, performance of the final model on the previously seen training data was retained.

For TFs which had more than one training cellular context, LOCO-rounds were performed, evaluating performance on cellular contexts not seen during training (see Supplementary Methods Figure 4). If more than four training contexts were available a maximum of ⌈*n*_*train contexts*_*/*4⌉ training contexts was held out per round. During each round, hyperparameters were tuned separately and models trained from scratch on the respective training data. Performances on the training data, chromosomes held-out from training contexts and completely held-out context were tracked.

After LOCO-rounds, models were trained on the feature matrix including all training cellular contexts but still with held-out chromosomes removed.

#### A.3.4 Generalizability estimation

The generalizability of a TF-specific classification model is quantified by removing the effect of several determinants that are not specific to the model itself (see Fig. 3H) from the observed performance difference between the training and held-out contexts.

We define this performance difference as the log2 enrichment over random *e* in the respective sets:

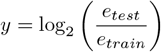

The contribution of the latter determinants is modeled using the lm function from the R package ‘stats’:

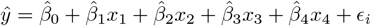

where *x*_1_ is the square root of the proportion of held-out peaks that are also present in the training data, *x*_2_ is the number of training contexts, *x*_3_ is the square root of the positive fraction in the held-out context, and *x*_4_ is the log2 of *e*_*train*_.

The generalizability of the classification model we then defined as,

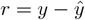

which is the residual performance difference between an unseen context and the training contexts not explained by these non-model-specific performance determinants.

#### A.4 Implementation

TFBlearner is available as an R package including all functionality for mapping data of the different assays to a common set of regions (here the PREs *E*), constructing features and training models.

The matched ATAC- and ChIP-seq data and motif matches are organized in a MultiAssayExperiment [110] object (see Supplementary Methods Fig. 5). Separate functions are available for computing features of different levels of specificity (see Table 1 and Supplementary Methods Table A4) which are then added as separate assays to the object using sparse matrix formats [111] if appropriate. Mapping between the assays of different levels of specification are maintained within the `sampleMap` of the MultiAssayExperiment object. Per-TF feature matrices are constructed from the MultiAssayExperiment object and are organized as SummarizedExperiment objects [112]. As both feature matrices in SummarizedExperiment containers and the original MultiAssayExperiment object tend to be large, we chose an on-disk representation using the HDF5 backend provided via the HDF5Array package [113].

The functions performing hyperparameter tuning and model training as previously specified make use of the lightGBM package [73] and packages of the mlr3-ecosystem [103].

The package can be installed from: https://github.com/ETHZ-INS/TFBlearner.

### A.5 Benchmark

The benchmark protocol has be pre-registered on the Open Science Framework (OSF) [52]. In the following parts of the pre-registered protocol repeated for convenience.

#### A.5.1 Inclusion criteria

We had defined two cellular contexts, embryonic stem cells and Jurkat cells, as testing contexts that were completely left out from our method development efforts. These cellular contexts constitute our global test/held-out dataset for the benchmark.

We then selected all the TFs that: 1) are in the list of bona fide TFs [25]; 2) have an associated motif; 3) have been profiled in either of these held-out cellular contexts; and, 4) that have (non-archived) data available on the ENCODE portal. We then identified the other cellular contexts on ENCODE where these TFs have also been profiled, and selected the 5 most common cell contexts (i.e. with the most TFs) from which ATAC data is also available. We then selected all TFs profiled in one of the held-out cell types, as well as in more than one of these 5 most common cellular contexts. We finally excluded CTCF, since it represents an unusually easy TF profiled in a large number of contexts. Wherever multiple ATAC-seq datasets are available from ENCODE for the same cellular context, their reads will be pooled. For cellular contexts from training and testing for which no ATAC data is available via the ENCODE platform, we retrieved matching datasets from Gene Expression Omnibus (GEO) (see Supplementary Table 3 for the final list of datasets).

**Supplementary Methods Fig. 4.**
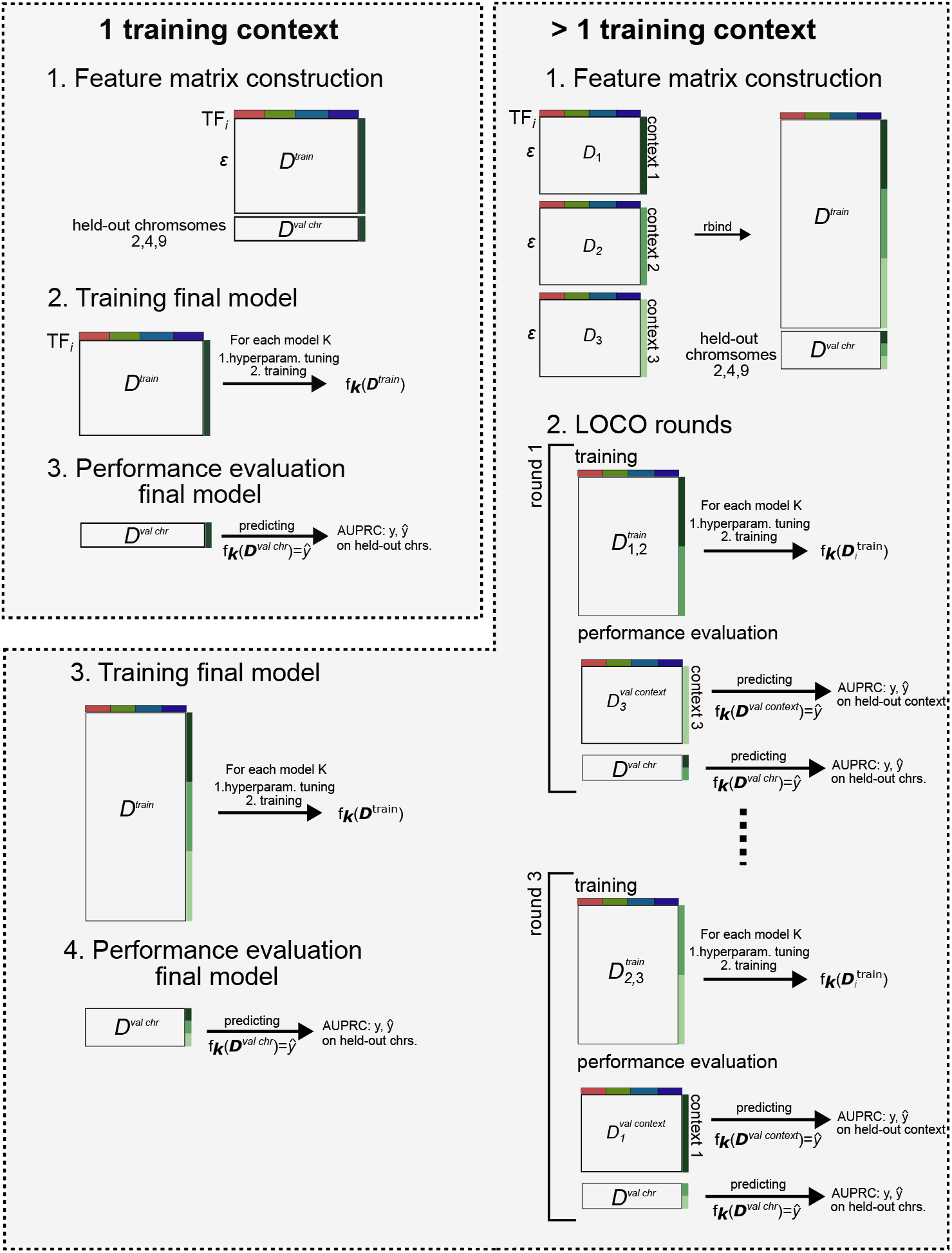
Performance evaluation model training: **Left upper**. training and model evaluation routine for TFs with one training cellular context. Hyperparameters for the final model are tuned with MBO assessing performance in 5-fold cross-validation rounds. **Right**. Similarly, for TFs with more than one training contexts for which LOCO rounds were carried out to evaluate performance on cellular contexts not seen during training. Hyperparameters were tuned by internal LOCO rounds, meaning that during each outer round they would be tuned separately on the contexts used for training. In the example here they would be tuned always on two training contexts (used for an inner LOCO) in each of the three outer LOCO rounds. For training of the final models hyperparameters would be tuned based on internal LOCO rounds on all three training contexts.

**Supplementary Methods Fig. 5.**
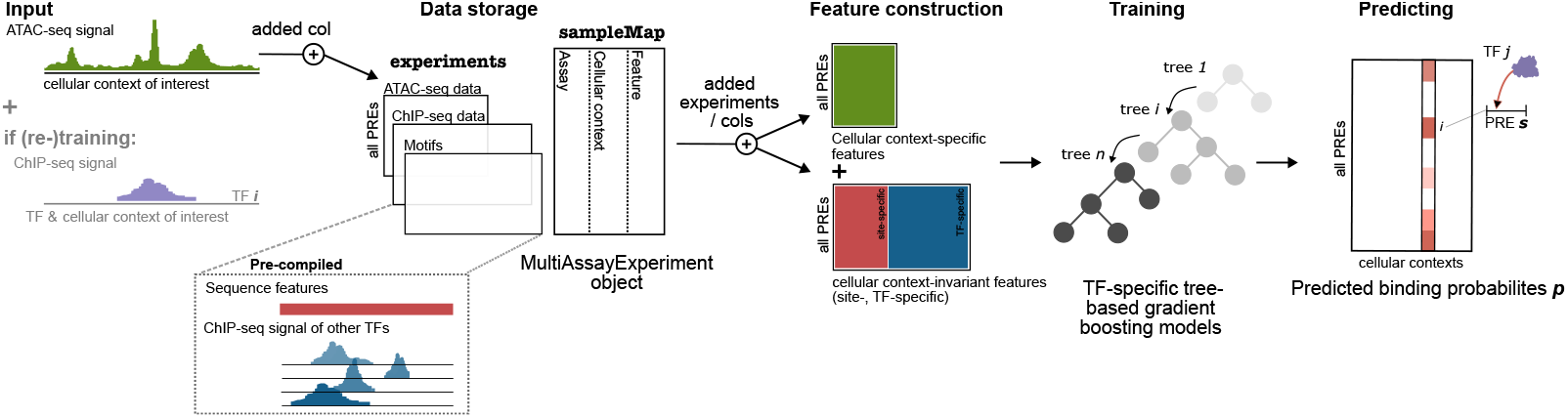
Overview TFBlearner package.

We first searched pubmed (on June 21st, 2024) for articles matching the query ‘predict “transcription factor” binding (“accessibility” OR “ATAC”)’, focusing on publications published after the ENCODE-DREAM challenge (i.e. 2018 onwards). We manually curated the resulting papers for prediction efforts of the kind investigated in our study, adding 3 prominent ones that had been missed by the search (FactorNet, TOP, and Leopard), resulting in 13 studies. We then went through each study, and noted the methods against which they compared and that seemed, according to that study, to be performing well. We excluded methods that require additional experimental data from the target cellular context (e.g. histone modifications). If a method was published before the ENCODE-DREAM challenge, it was excluded. If a method was shown inferior to another in a study, it was excluded. If a method was published by the same authors as a more recent method, it was excluded. Finally, we excluded methods that did not provide code or documentation. This left us with the following methods: maxATAC 1.0.6 [43] (version 1.0.6), Catchitt [37] (version 1.0.4) and TOP [54] (version from the publication), and our own method TFBlearner. In addition, we included another method, a faster reimplementation of BMO [53], as a baseline unsupervised motif-based method.

Subsequently, for training of the methods, they were excluded if they repeatedly used more than 120 GB of RAM or more than a day for a training step (using 20 threads if supported).

#### A.5.2 Peak merging

The merged peak set (‘Merged’) used for evaluation and for labeling positive instances during training was constructed based on conservative IDR-thresholded peaks and IDR-thresholded peak files from ENCODE. Conservative IDR-thresholded peak files across replicates and different ENCODE datasets with the same specification were stacked and combined by `reduce` function of the GenomicRanges [72] R-package (v1.60.0). Non-conservative IDR-thresholded peaks were then incorporated into this set through two additional recursive merging steps, in which peak boundaries were defined by the median start and end coordinates of overlapping peaks (see Supplementary Methods Fig. 6). IDR-thresholded peaks which were found with no overlapping peak in any other ENCODE dataset with the same specification were removed from the set.

#### A.5.3 Baseline scores

Apart from the pre-registered methods several baseline scores *s* were defined:

- ATAC+motif: Computed as the width-normalized nuclesome-free fragment overlap count (see below) multiplied by the square-root of the motif matching score *m* as follows:

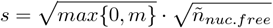
- ATAC.nucleosome.free: Nucleosome-free fragment overlap count normalized by the width *w* of the evaluation site *e*:

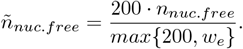
- ATAC.totaloverlaps: Total fragment overlap count normalized by the width as above.
- ChIP.any: If any ChIP-seq peak was observed at an evaluation site *e* in any of the training cellular contexts (binary)
- ChIP.max: Maximum ChIP-seq signal (*r*-score) observed at *e* in any of the training cellular contexts.
- ChIP.mean: Mean ChIP-seq signal (*r*-score) observed at *e* in any of the training cellular contexts.
- ChIP.nearest: ChIP-seq signal (*r*-score) of the same TF in the training cellular context closest to the respective testing context. Closest training context determined by cosine similarity on the ATAC-signal at the PREs.
- C-Score: The C-score was added as an context unspecific baseline score as detailed in (Supplementary Methods A.2.6). For the PREs the C-score was computed based on the full training ChIP-seq matrix *B* used in the large scale prediction effort with all benchmark TFs removed.
- FIMO: Motif matching scores (log likelihoods). For PREs *E* a lenient motif matching threshold of 0.01 was used, whereas for the 200 bp bins on the full chromosomes the default threshold of 1e-4 Also used as inputs for methods requiring motif matches.

**Table A4:**
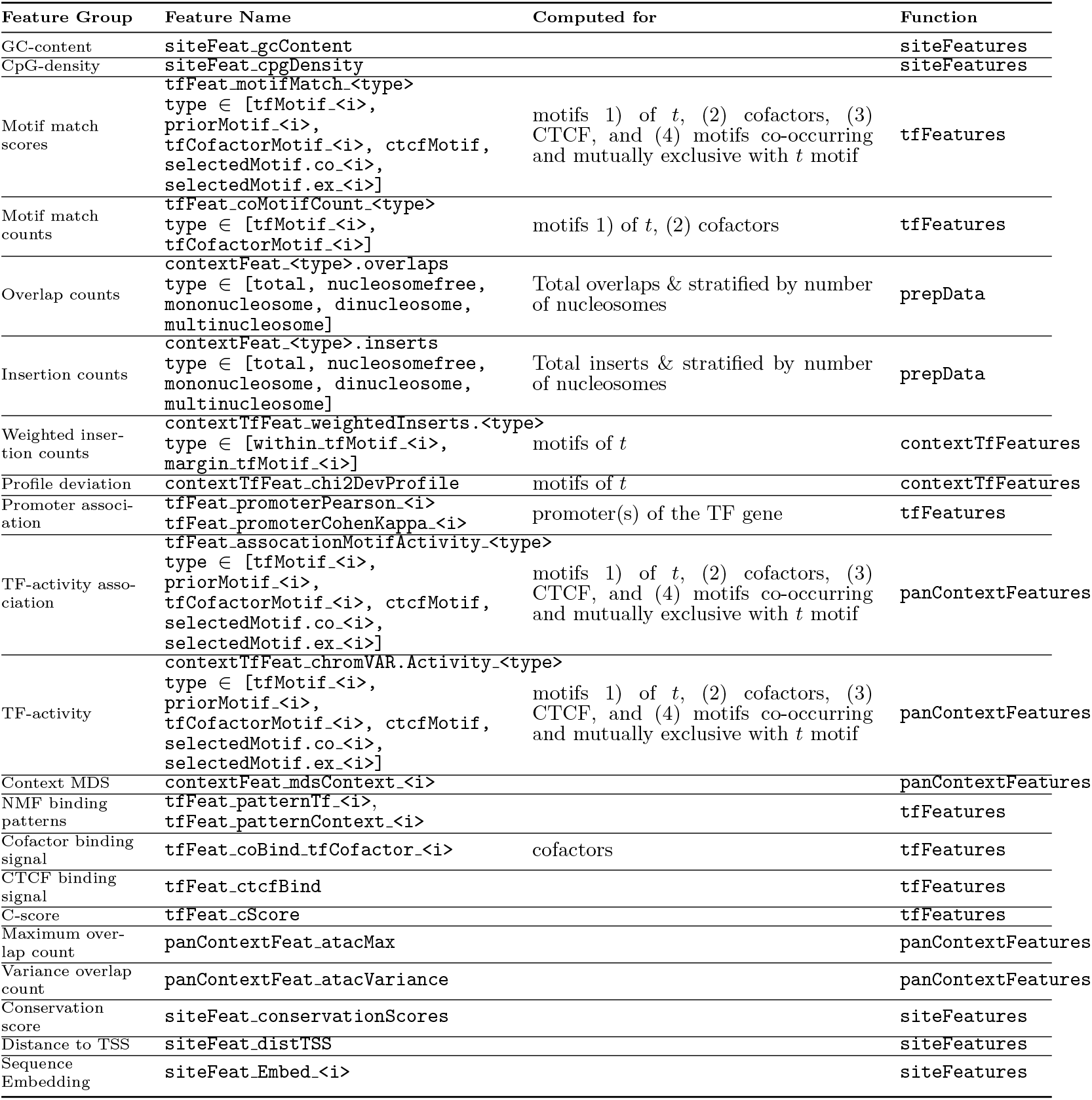
Overview constructed features: All feature groups, along with their respective construction functions and column names in the feature matrix, are shown.

**Supplementary Methods Fig. 6.**
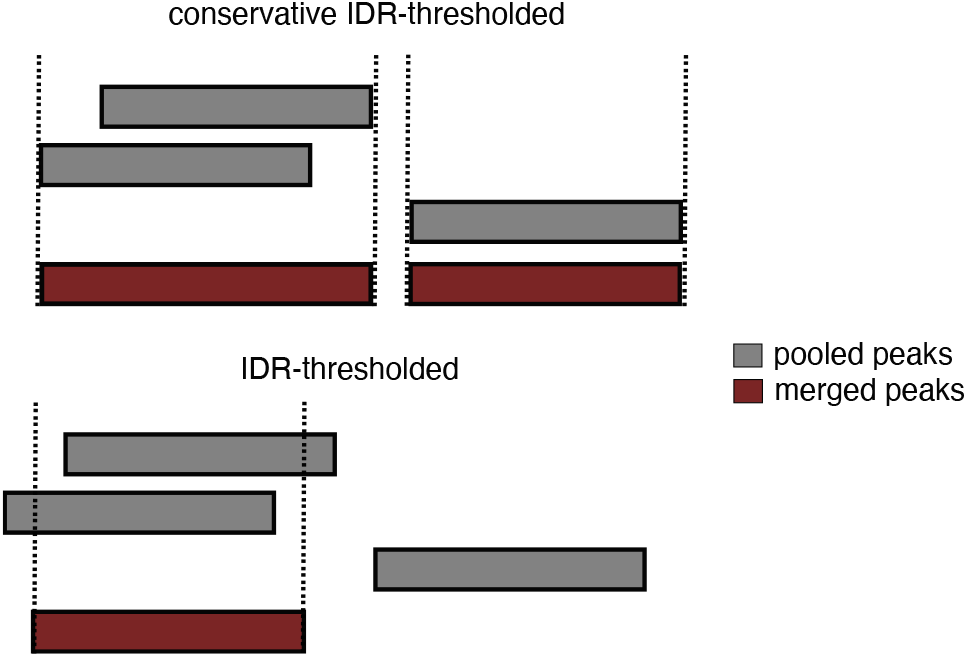
Peak merging. Conservative IDR thresholded peaks were merged if found overlapping by taking the full span of the overlapping peaks (`GenomicRanges::reduce`). If no overlapping peak was found the original peaks were kept as is. For IDR-tresholded peaks only these overlapping peaks form another ENCODE dataset with the same specifications were kept. Overlapping was performed in the recursive merging rounds during which each time the median of the overlapping peaks was taken as the merged boundaries.

#### A.5.4 Methods score evaluation

For both comparisons, a) on the set of PREs *E* and b) the full chromosomes in 200 bp bins, evaluation was restricted to chromosomes 2,4 and 9.

For each method, scores for an evaluation site (200 bp bin or PRE) were obtained by first checking if any of the predicted ranges was found within that site. If so the score of that prediction was taken for evaluation. If not we took the maximum score of any predicted range overlapping the evaluation bin as the method’s score.

Labels for each evaluation site were obtained by checking if any peak center margin (peak midpoint ±15 bp) was found overlapping. If so the evaluation site was labeled as positive for the respective TF-cellular context combination. If no peak center margin was found overlapping but any other part of the peak, the evaluation site was labeled as ambigous negative and removed from the evaluation. The remaining instances with no peak found to overlap were used as negative instances.

AUPRC was computed using the PRROC [74] R package (v.1.4), precision at 5% and 10% recall based on a custom implementation deposited on the repository with the benchmark’s code.

#### A.5.5 Deviations from preregistration

Non pre-registered (additional) exploratory analysis:

- The enrichment over random was not listed as a metric in the protocol. However, its computed directly based on the pre-registered AUPRC divided by the positive fraction.
- The detailed analysis of the performance on different definitions of positives and negatives based on if a peak was observed already during training as shown in Supplementary Fig. 9.
- Apart from ChIP.mean none of the baseline scores added for comparison were pre-registered. Pre-registered but not performed:
- Pre-trained models were not found to be available for any of the competitor models, hence a comparison of pre-trained models was dropped.

Deviations from the pre-registered protocol:

- For BMO we used the original method instead of the re-implementation.
- For TOP we used the version v.1.0.1 which might differ from the version in the publication. Further, we had not specified in the pre-registered protocol which model (quantitative occupancy or logistic) to use. For the benchmark, we employed the logistic model to enable direct comparison with the other classification models.
- The Catchitt version pre-registered was mistakenly given as v1.0.4, however the version which actually exists and we used is v0.1.4.
- Methods were trained using 4 Snakemake threads to allow training of more TF-specific models in parallel.
- maxATAC.100 was trained on a GPU instead of CPUs as the other methods.

#### A.5.6 Data and Code availability

Pre-registered protocol: https://osf.io/kv2tw/overview

The benchmark was implemented as a snakemake workflow (v.9.8.2), with a main workflow containing all common preprocessing steps and separate rule-files for all methods. Maximum RSS and computation time were tracked using the snakemake `benchmark` directive. The code used is available under: https://github.com/ETHZ-INS/TFB_Prediction_Benchmark

## Appendix B Supplementary Figures

**Supplementary Fig. 1.**
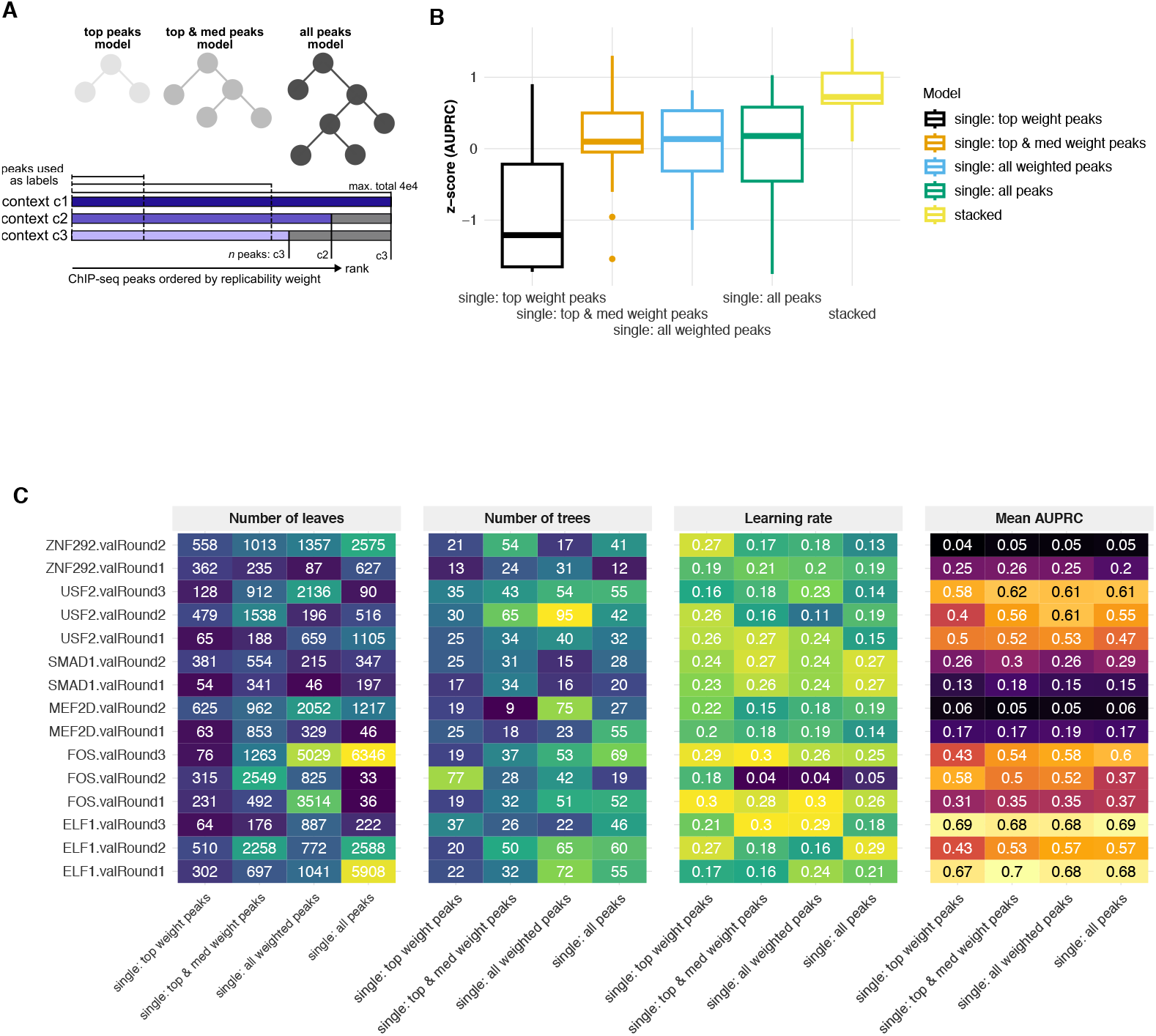
Overview model stacking. **A**: Single GBDT models are trained on different subsets of the data. Therefore, ChIP-seq peaks are ranked in descending order by their *r*-score and three different nested subsets of increasing size are obtained by including positives with successively higher ranks in the respective sets (see Methods). The maximum set size of positives considered is 4e4 for the third model (‘all peaks’). Three models are trained on these subsets using the *r*-score as instance weight, a fourth model is trained by randomly sampling positives with the same number of positives as the third model but without instance weights. For training, negative instances are subsampled such that a positive proportion of 0.25 is obtained. **B**: AUPRC z-score obtained for the individual models and the ensemble model in cross-validation rounds leaving out entire held-out contexts. **C**: MBO selected hyperparameters for the individual models and the respective AUPRC obtained on the held-out contexts.

**Supplementary Fig. 2.**
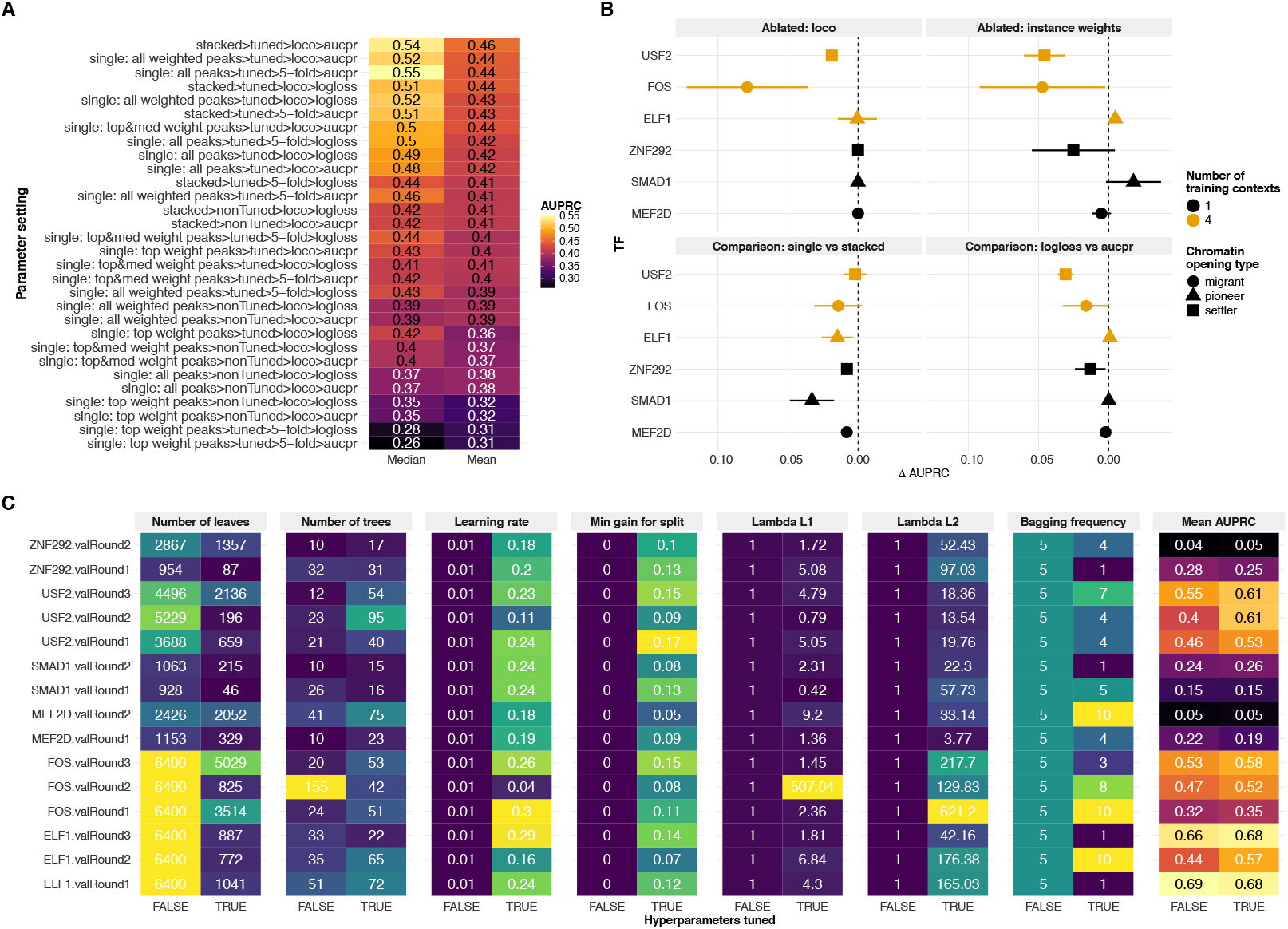
Model component comparisons and ablations. **A**: Comparison of different modeling choices from left to right (*>*); stacked model/individual model, MBO hyperparameter tuning applied, leave-out procedure for MBO, metric for MBO (see Supplementary Methods A.3.2). All performances are estimated in cross-validation rounds with entire cellular contexts held-out. Mean and median computed across all indicated TFs and their validation rounds. **B**: Ablation of model components or specific comparisons of modeling options. For each TF mean and standard errors of differences across validation rounds are shown. When ablated, LOCO rounds are compared to random split 5-fold cross-validation rounds. For the ablation of instance weights two of the individual GBDT with (‘all weighted peaks’ model) and without instance weights (‘all peaks’ model) are compared. For the single vs stacked model comparison, the model trained with instance weights on the largest training dataset (‘all weighted peaks’ model) is compared to the stacked model. **C**: Comparison of hyperparamters and performance of an individual model (‘all weighted peaks’ model) trained with MBO tuned hyperparameters and non-tuned hyperparameters chosed based on the training dataset size (see Supplementary Methods Table 3). Selected hyperparameters are indicated always in the left column and compared to their default counterpart (determined based on the training dataset size).

**Supplementary Fig. 3.**
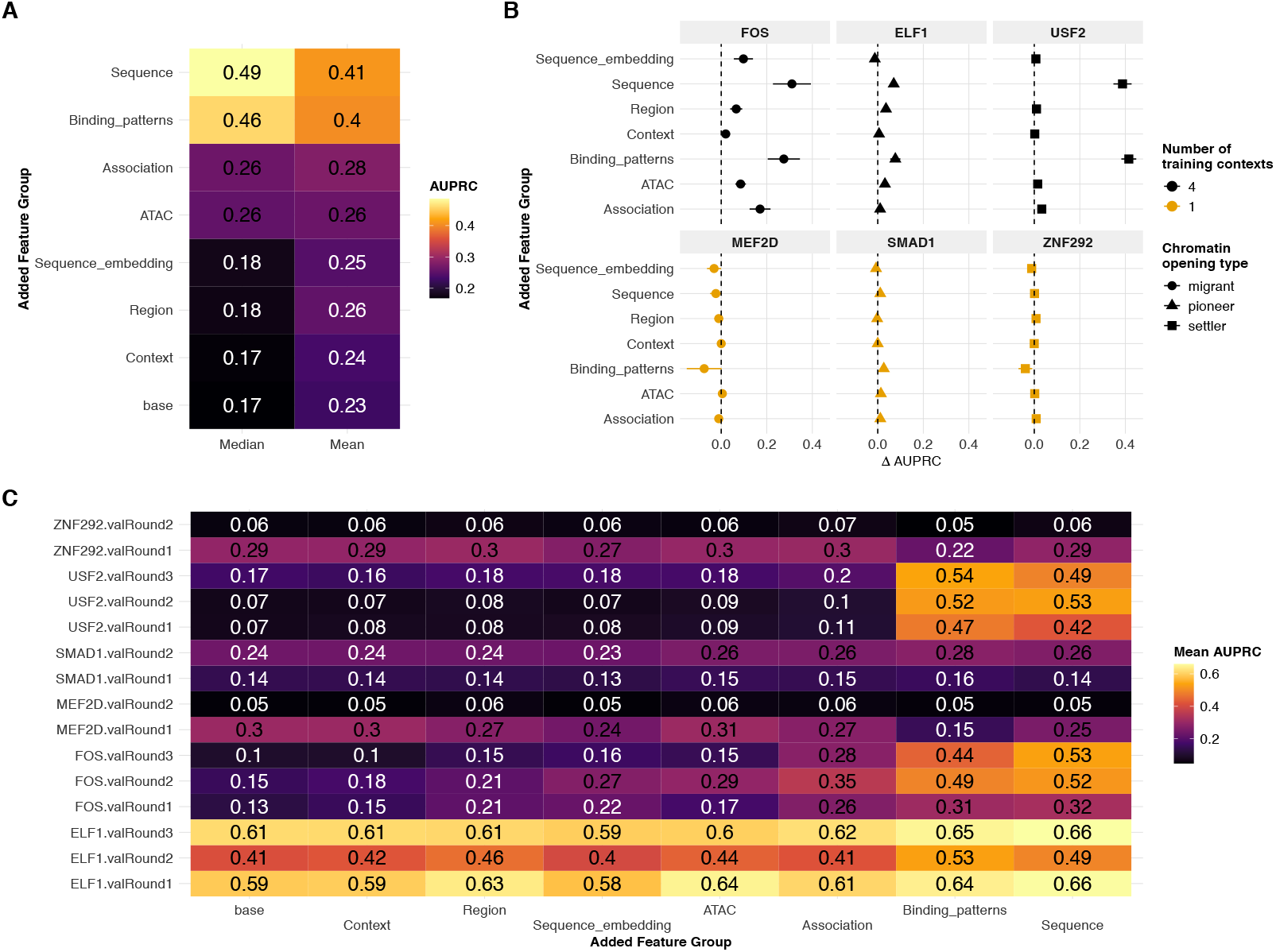
Feature group additions. **A**: Median and mean AUPRC obtained for models trained on base set of features (ATAC-seq fragment counts and motif matches of the given TF) compared with models trained with one additional feature group (see Table 1). Further detailed for the individual TFs (**B**) and each validation round of each TF (**C**).

**Supplementary Fig. 4.**
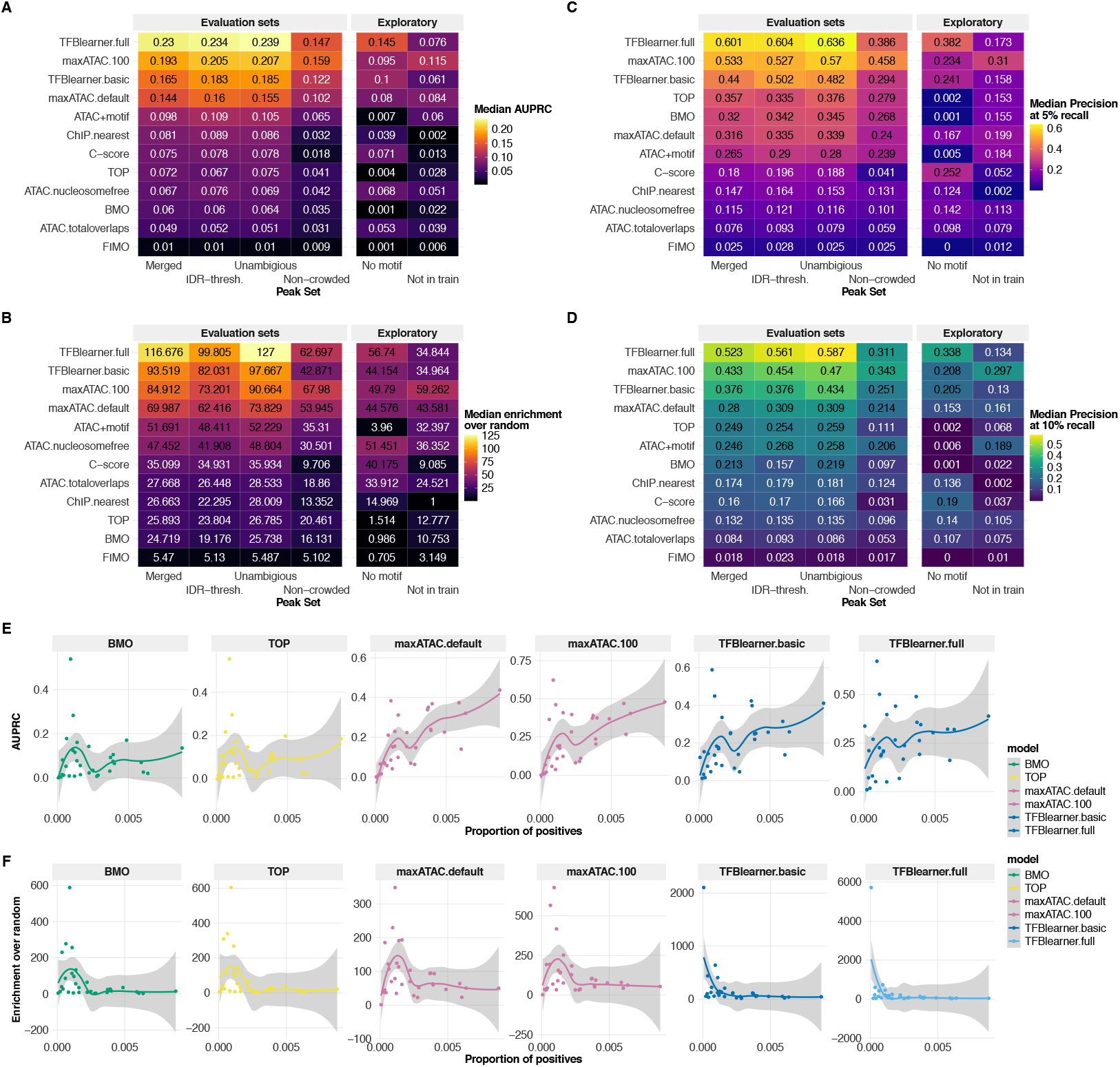
Benchmark performance on different peak sets. **A**: Median AUPRC obtained across 32 testing TF-cellular context combinations. For comparison between peak sets, positive proportions of all sets were matched to the merged set by subsampling. The equivalent for fold change over enrichment (**B**), precision at 5% recall (**C**) and precision at 10% recall (**D**). **E**: AUPRC in dependence of the positive proportion in the respective held-out TF-cellular context combinations. Trend estimated by Local Polynomial Regression Fitting (Loess) using `geom smooth` of `ggplot` R package. **F**: The same but for fold enrichment over random.

**Supplementary Fig. 5.**
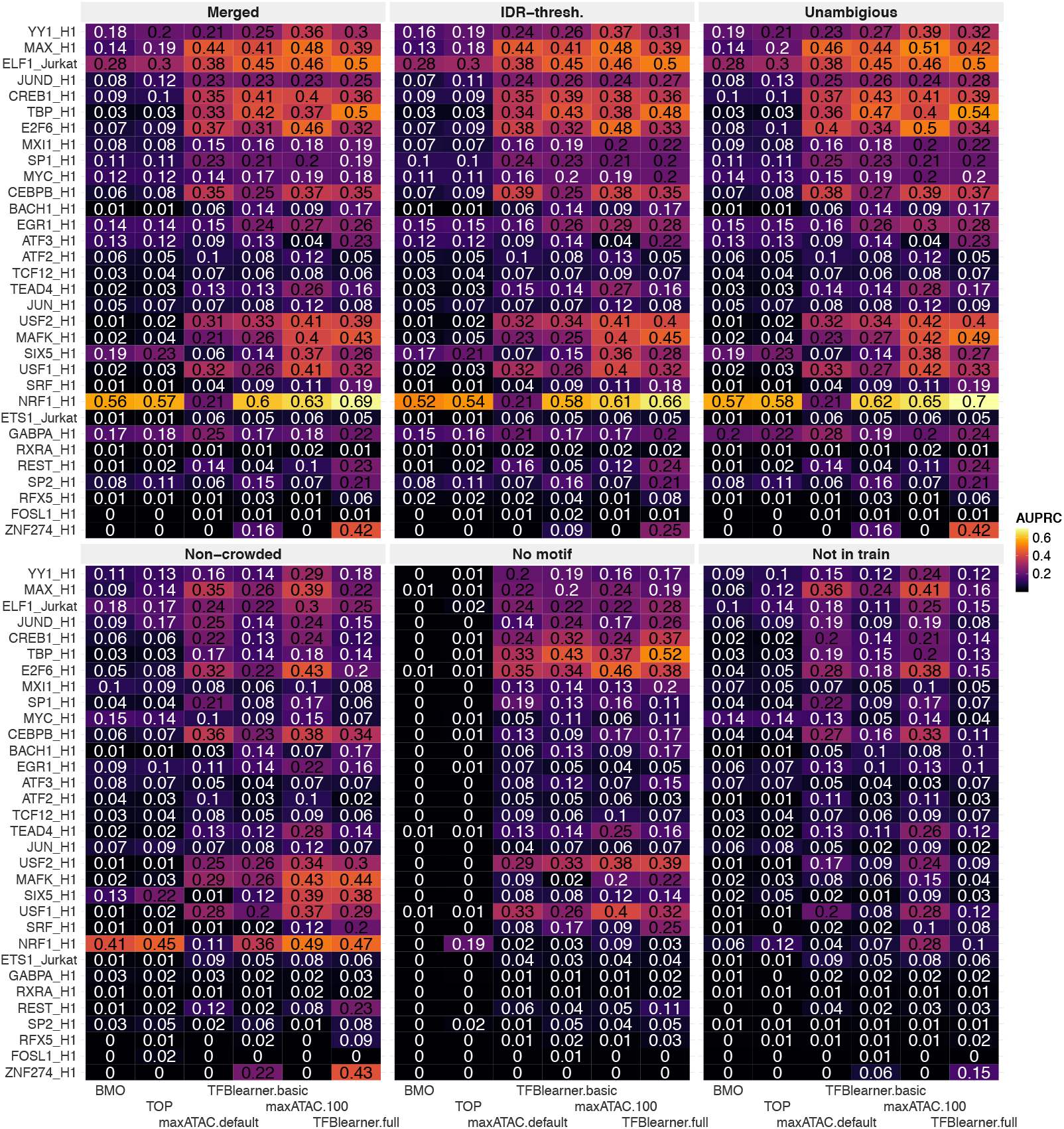
AUPRC for all testing TF-cellular context combinations. AUPRC shown for all methods and all TF-cellular context combinations evaluated on the set of PREs.

**Supplementary Fig. 6.**
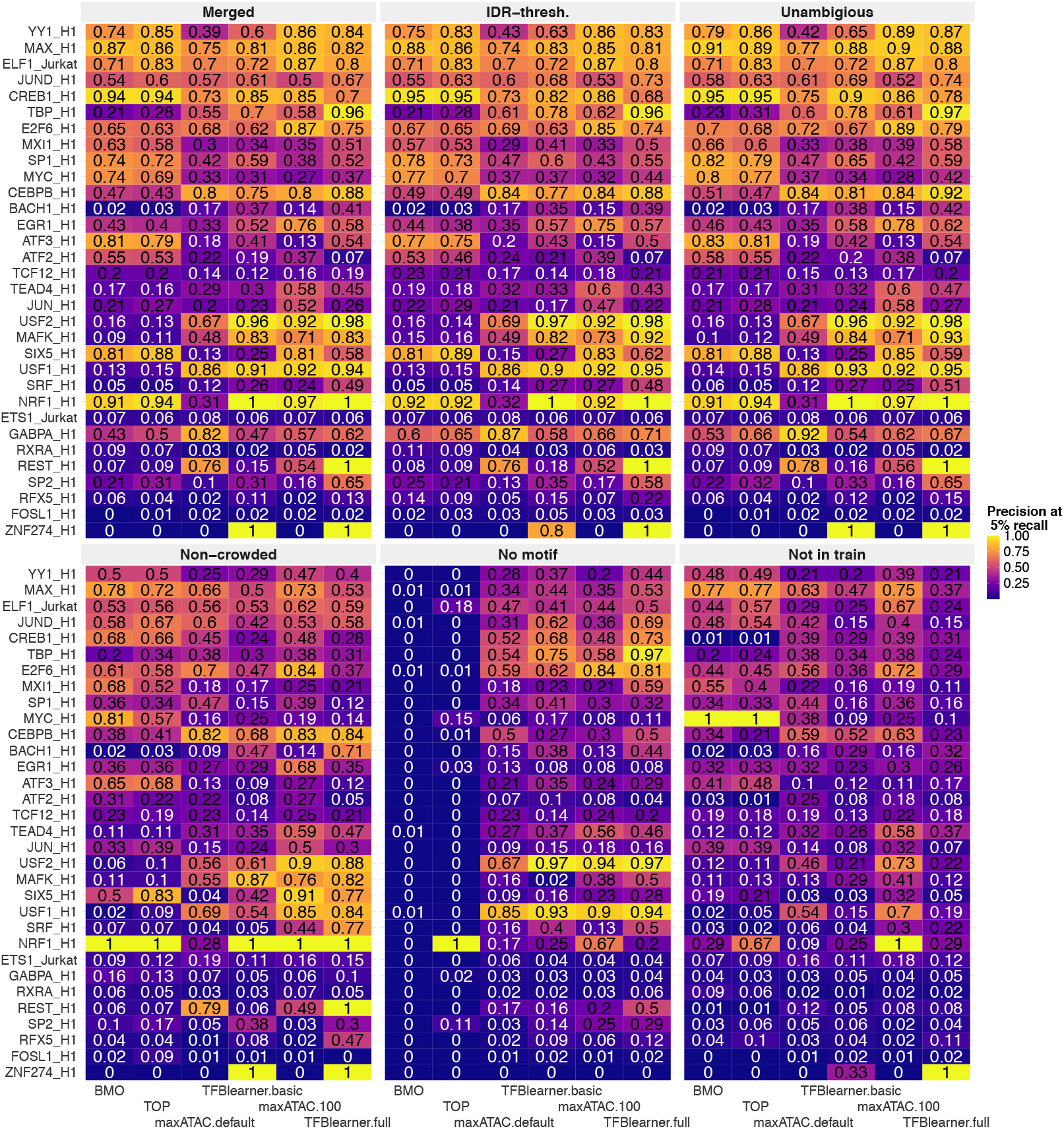
Precision at 5% recall on all testing TF-cellular context combinations. Precision at 5% recall shown for all methods and all TF-cellular context combinations evaluated on the set of PREs.

**Supplementary Fig. 7.**
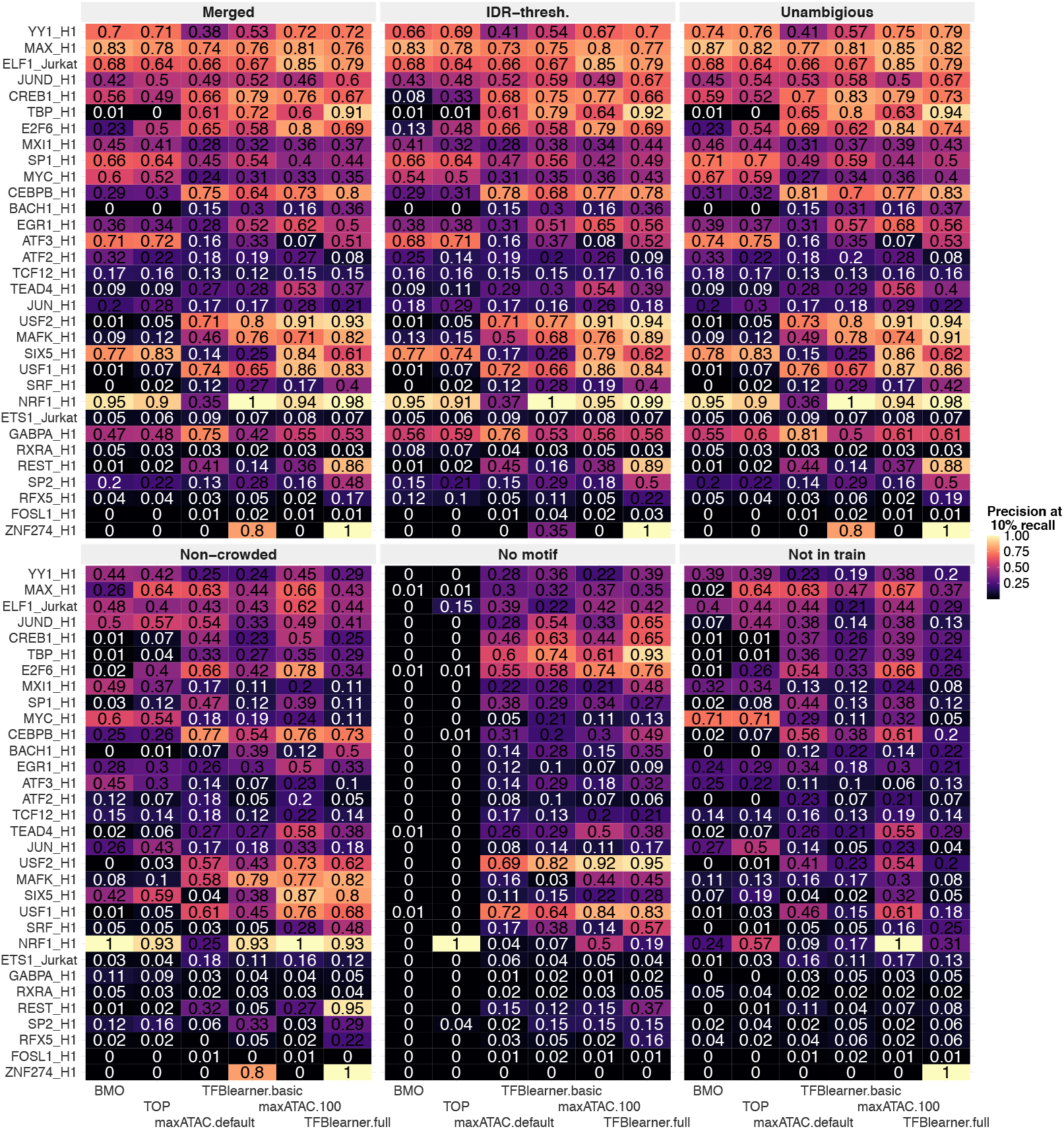
Precision at 10% recall on all testing TF-cellular context combinations. Precision at 10% recall shown for all methods and all TF-cellular context combinations evaluated on the set of PREs.

**Supplementary Fig. 8.**
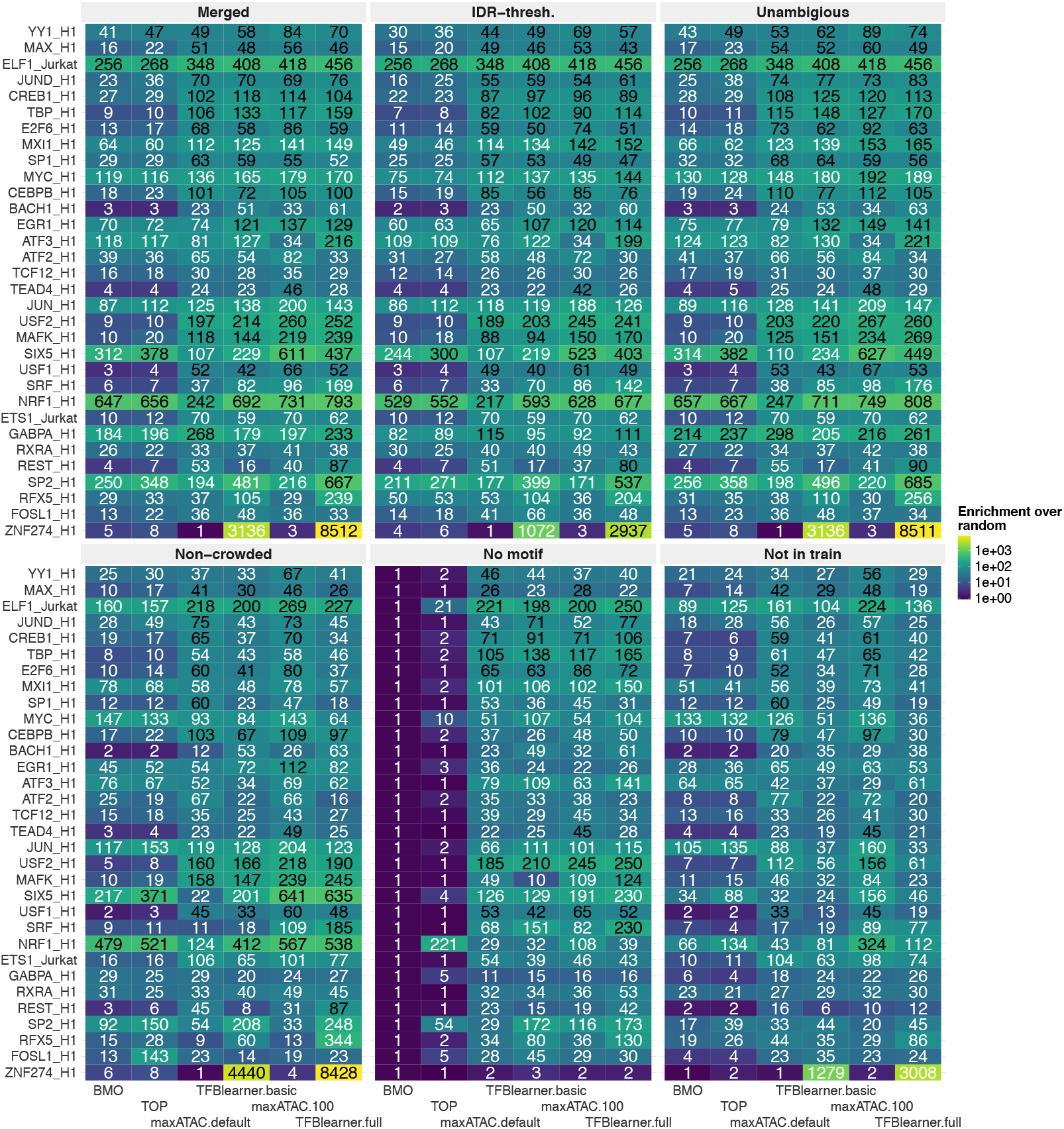
Enrichment over random on all testing TF-cellular context combinations. Enrichment over random shown for all methods and all TF-cellular context combinations evaluated on the set of PREs.

**Supplementary Fig. 9.**
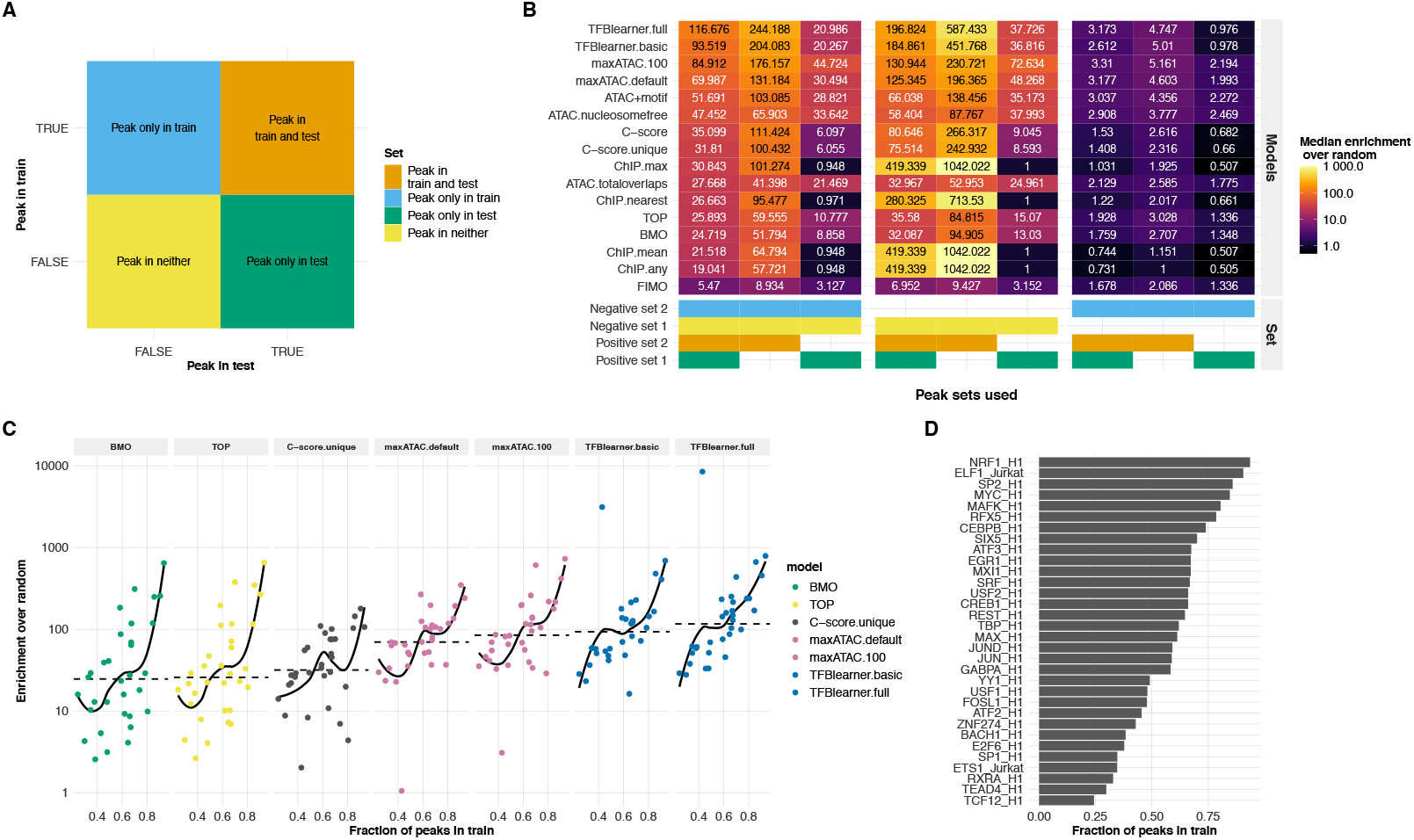
Benchmark performance in dependence of fraction of peaks observed in training contexts. **A**: Overview scheme for different sets of sites defined by whether at a site a peak is observed for a given TF in a training and/or the testing cellular context or never. **B**: Enrichment over random for the different combinations of sets constituting the negative and positive instances. **C**: Dependence of methods performance (enrichment over random) on the fraction of peaks observed in training. **D**: For each testing TF-cellular context combination the fraction of peaks observed in any of the training contexts.

**Supplementary Fig. 10.**
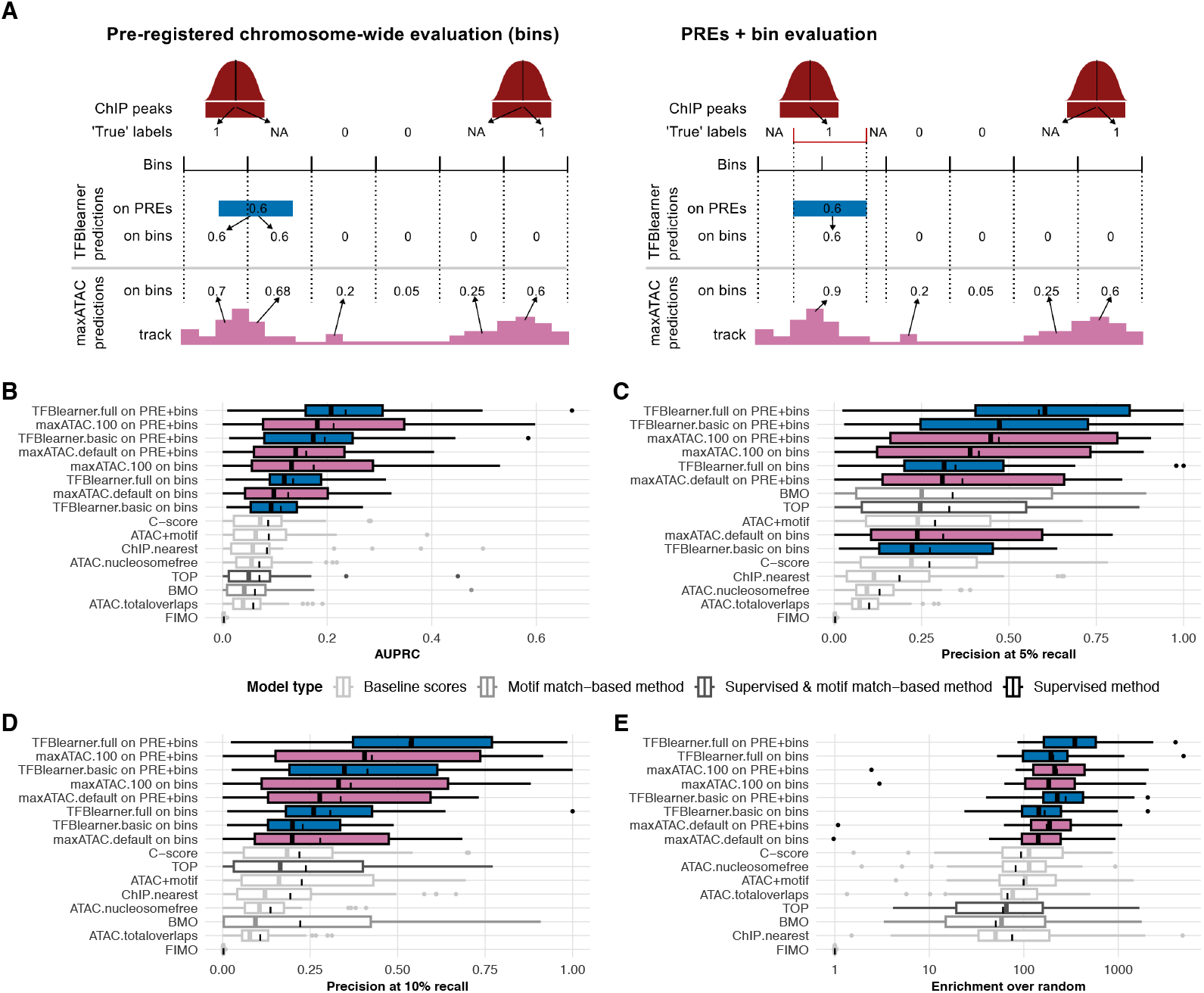
Benchmark performance on full chromosomes. Equivalent to Fig. 2 with performance metrics computed on full chromosomes 2,4 and 9, divided into non-overlapping 200bp bins. **A:** Scheme of the two ways in which TFblearner and maxATAC were evaluated. As pre-registered, true labels and predictions are ported to the bin resolution for evaluation, in the way illustrated on the left (and referred to in the following panels as ‘on bins’). Alternatively, as illustrated on the right (and referred to in the following panels as ‘PREs + bins’), whenever bins overlap a PRE the true labels and predictions are ported on the resolution of the PRE for evaluation, whereas the bin resolution is used for all bins not overlapping a PRE. In both cases, a ‘true’ binding is considered if the peak center is contained in the window (e.g. bin), while overlapping bins not containing the center are treated as neither positive nor negative and removed from evaluation. Methods are then ranked by their median performance metric, AUPRC (**A**), enrichment over random (**B**), precision at 5% recall (**C**) and precision at 10% recall (**D**). Notably, the discrepancy between the two evaluations schemes (bins vs ‘PREs+bins’) indicates that TFBlearner’s drop in performance when going from PREs to the full genome is not due to bindings not overlapping PREs, but to limits in its PRE-level resolution. An implication is that if more fine-grained resolution is needed, maxATAC might be preferable.

**Supplementary Fig. 11.**
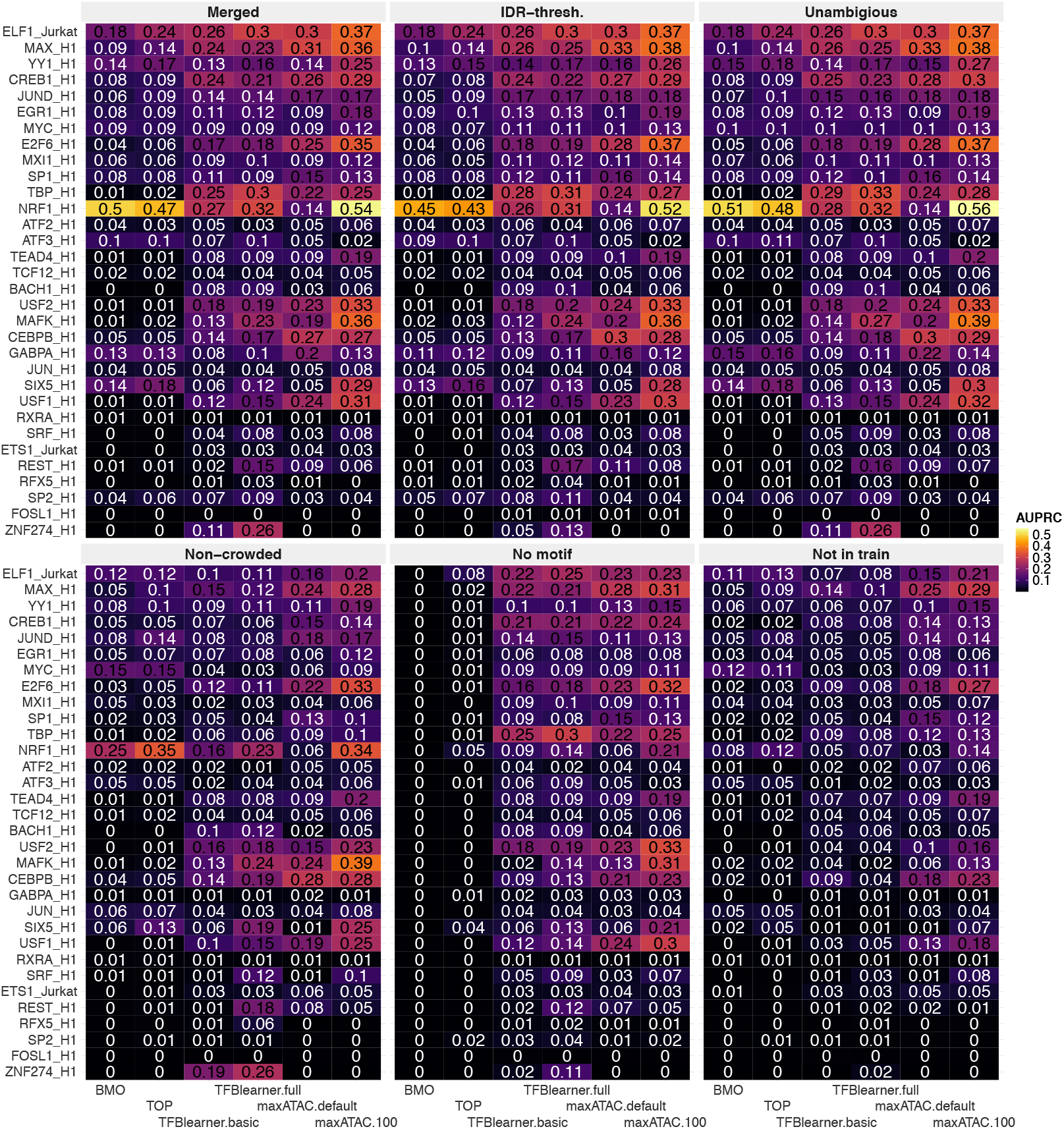
AUPRC on all testing TF-cellular context combinations. Enrichment over random shown for all methods and all TF-cellular context combinations evaluated on full chromosomes.

**Supplementary Fig. 12.**
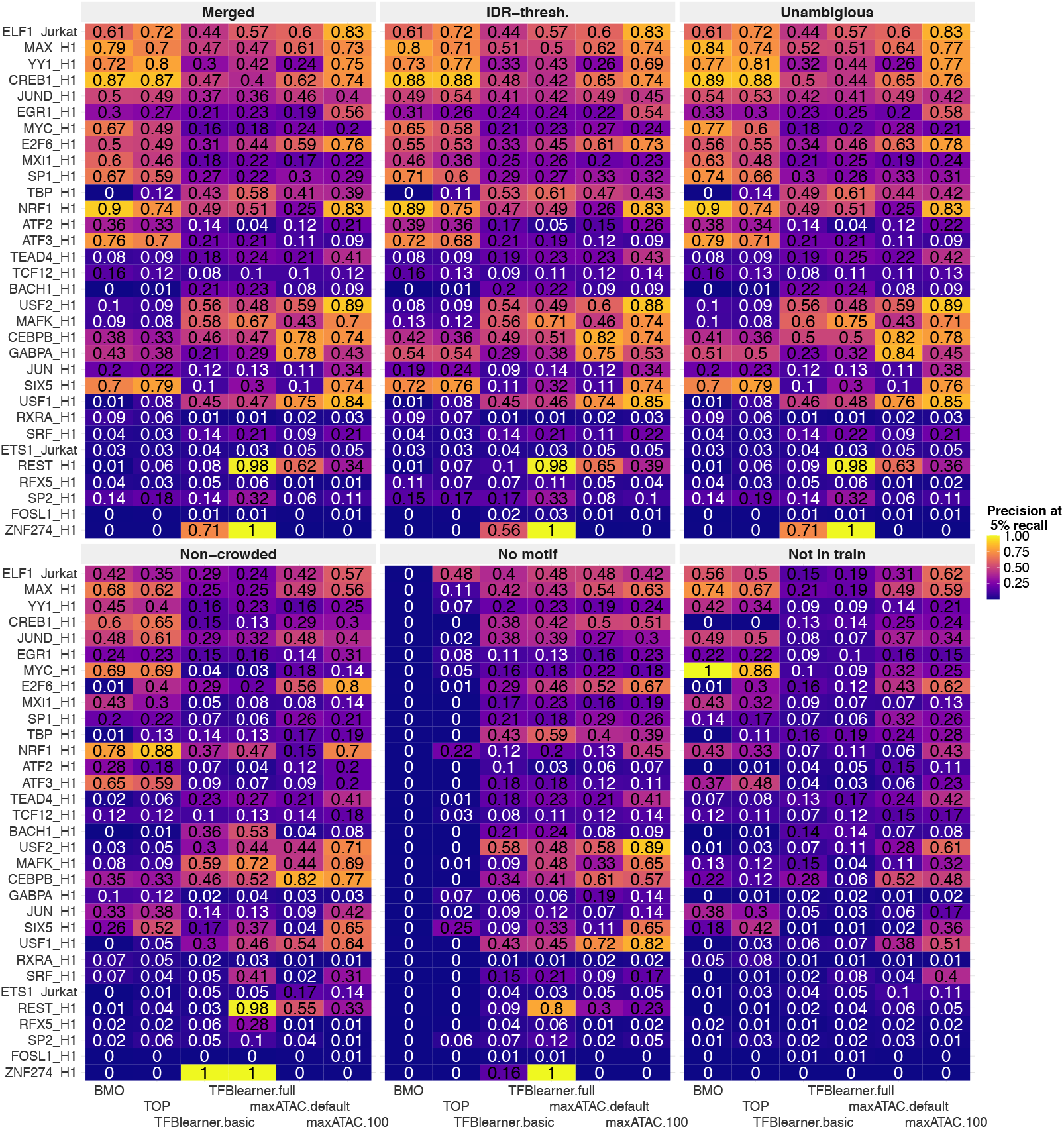
Precision at 5% recall on all testing TF-cellular context combinations. Enrichment over random shown for all methods and all TF-cellular context combinations evaluated on full chromosomes.

**Supplementary Fig. 13.**
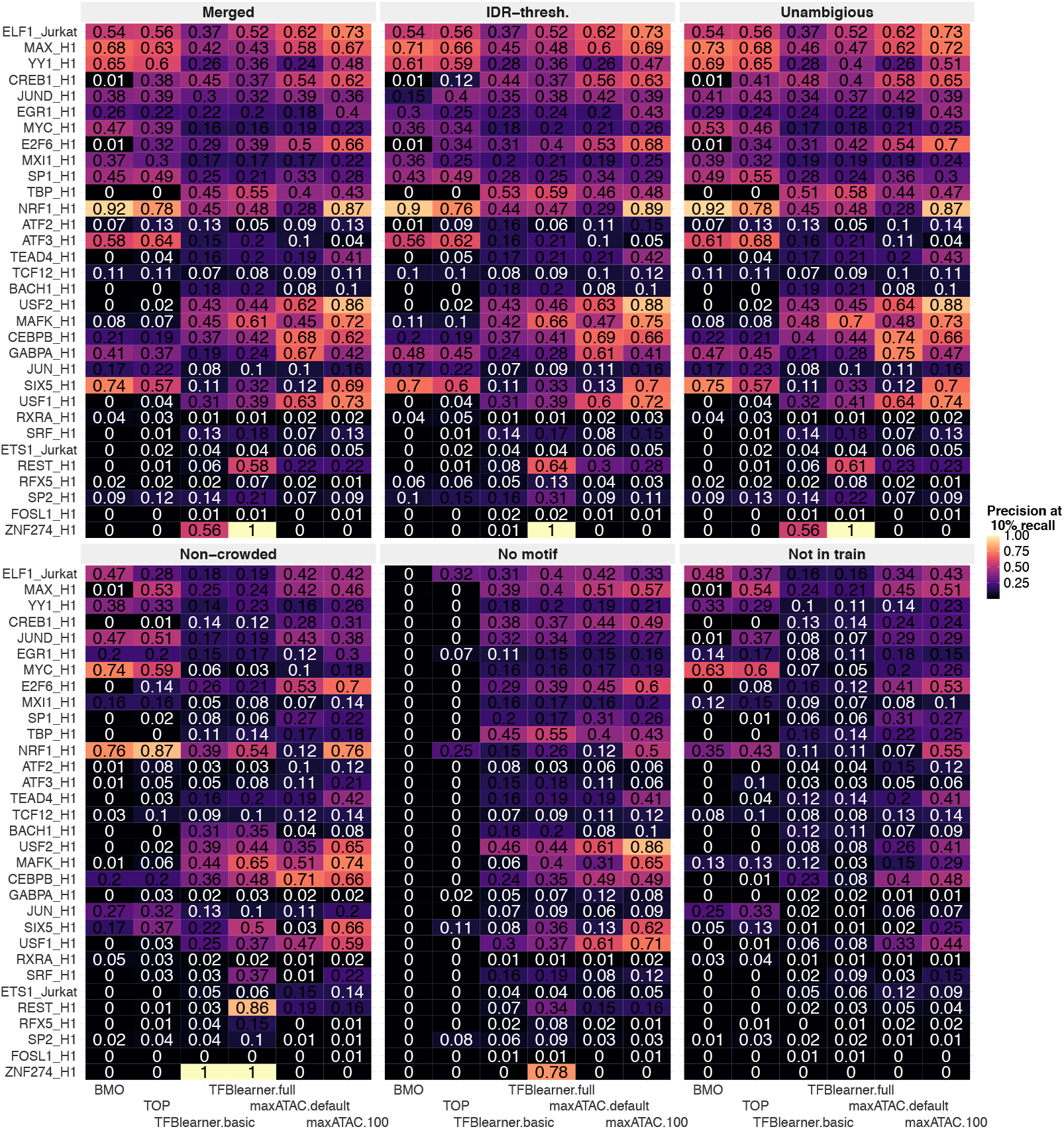
Precision at 10% recall on all testing TF-cellular context combinations. Enrichment over random shown for all methods and all TF-cellular context combinations evaluated on full chromosomes.

**Supplementary Fig. 14.**
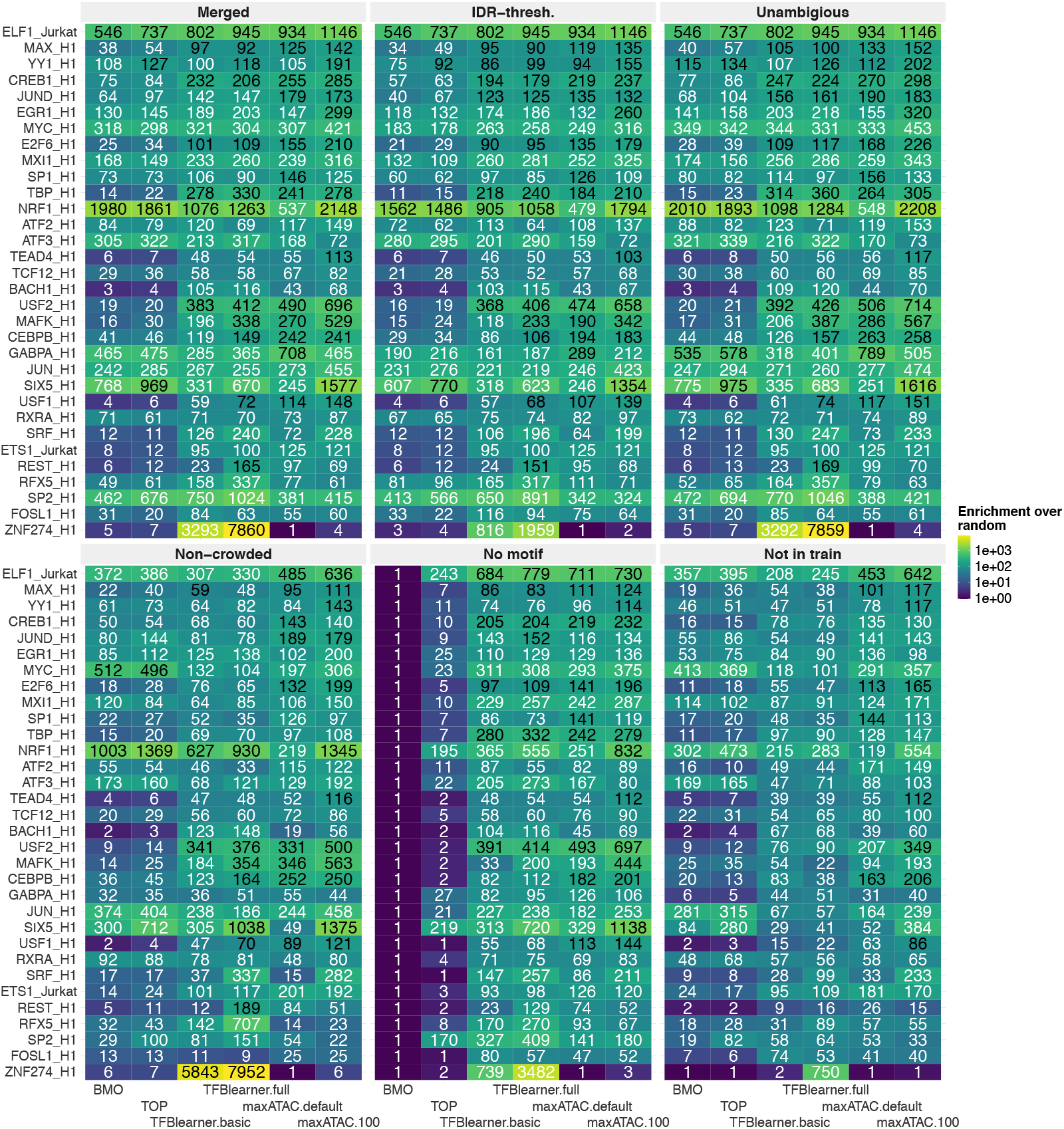
Enrichment over random on all testing TF-cellular context combinations. Enrichment over random shown for all methods and all TF-cellular context combinations evaluated on full chromosomes.

**Supplementary Fig. 15.**
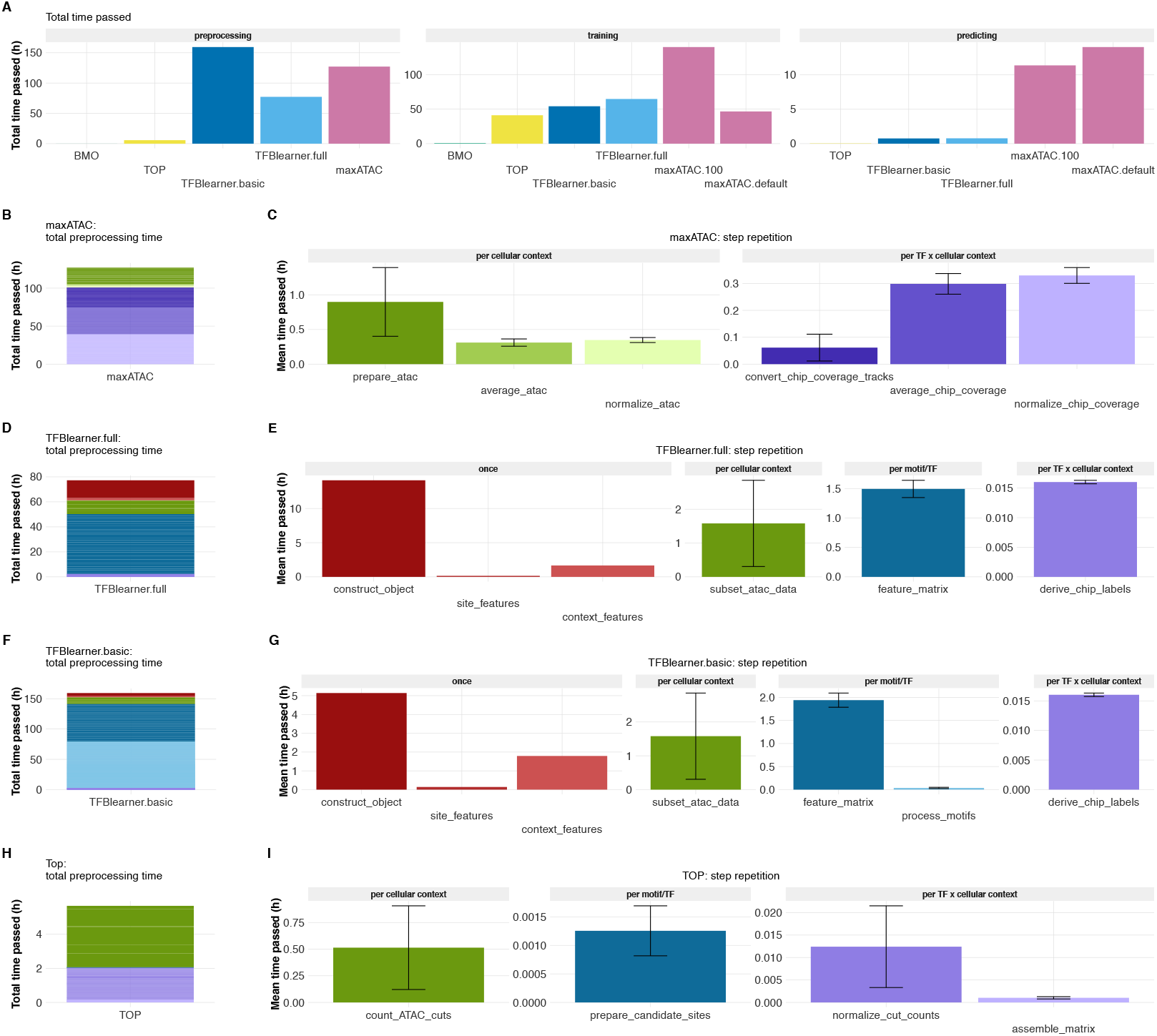
Run times of benchmarked methods. **A**: Total time passed for methods-specific preprocessing steps, training and predicting. BMO is an unsupervised method and fits several negative binomial models, the time required for this step is shown in the training panel. TFBlearner.full uses generic precompiled data for some of the features and thus omits several preprocessing steps which TFBlearner.basic conducts. **B**: total preprocessing time for maxATAC (same for maxATAC.default and maxATAC.100). **C** Preprocessing time stratified by the level of repetition, e.g. for which level of specification it has to be conducted. Levels of specification are indicated by color groups; red – once, green – per cellular context, blue – per TF/motif and purple – per TF-cellular context combination. Equivalently for TFBlearner.full (**D, E**), TFBlearner.basic (**F, G**) and TOP (**H, I**). Omitted for BMO as it requires no additional method-specific preprocessing steps beyond the data preprocessing common to all methods (not shown).

**Supplementary Fig. 16.**
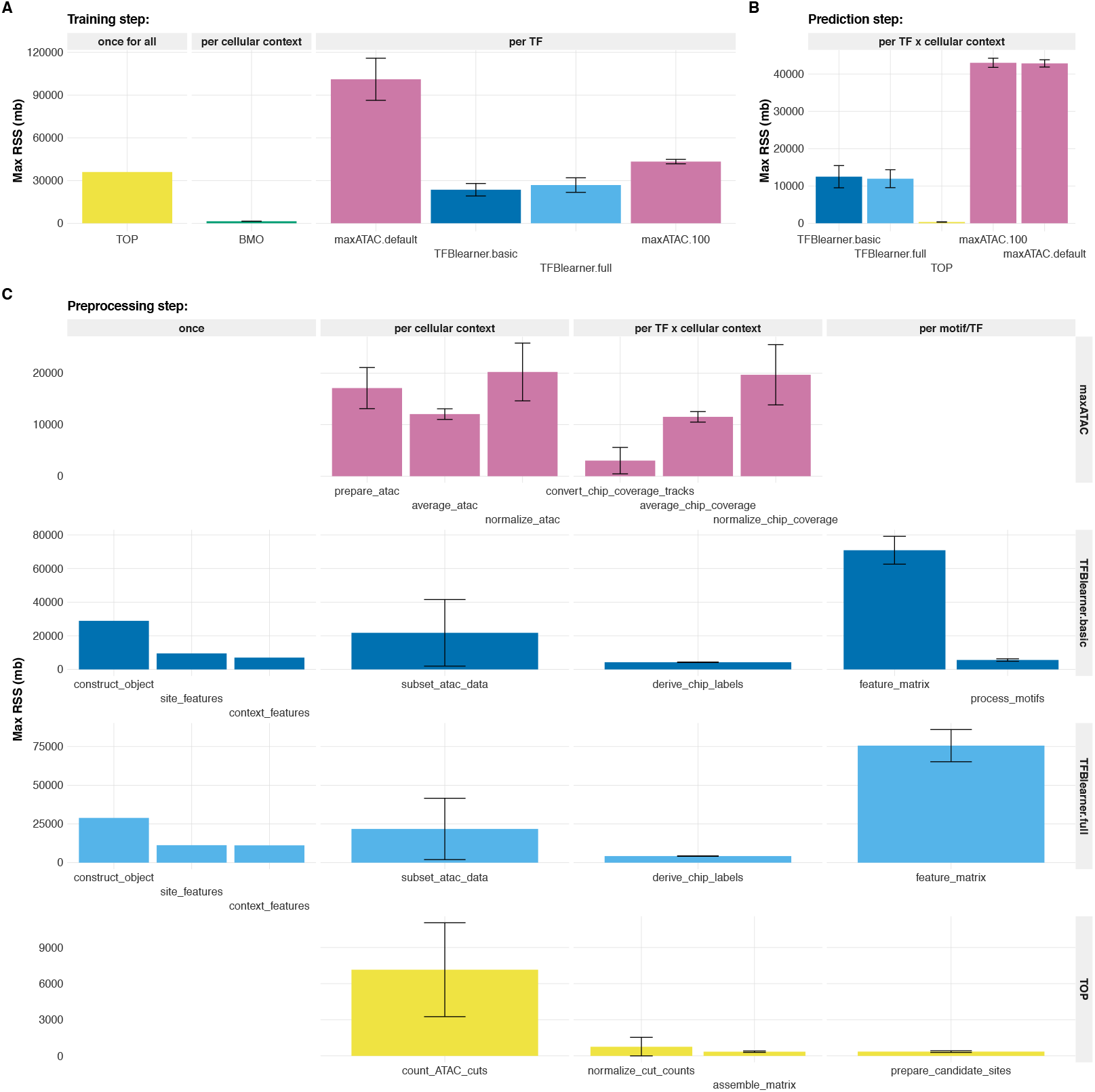
Maximum memory required by benchmarked methods. **A**: Maximum resident set size (RSS) required by methods for training. The panel is stratified by the level of specification at which training is conducted, e.g. TOP trains one model across all TFs and cellular contexts, BMO fits distributions per cellular context, whereas maxATAC and TFBlearner variations are single-task models trained per TF. MaxATAC.100 is trained on a GPU while all other methods are trained on CPUs. **B**: Accordingly for the prediction step and (**C**) stratified for the preprocessing steps.

**Supplementary Fig. 17.**
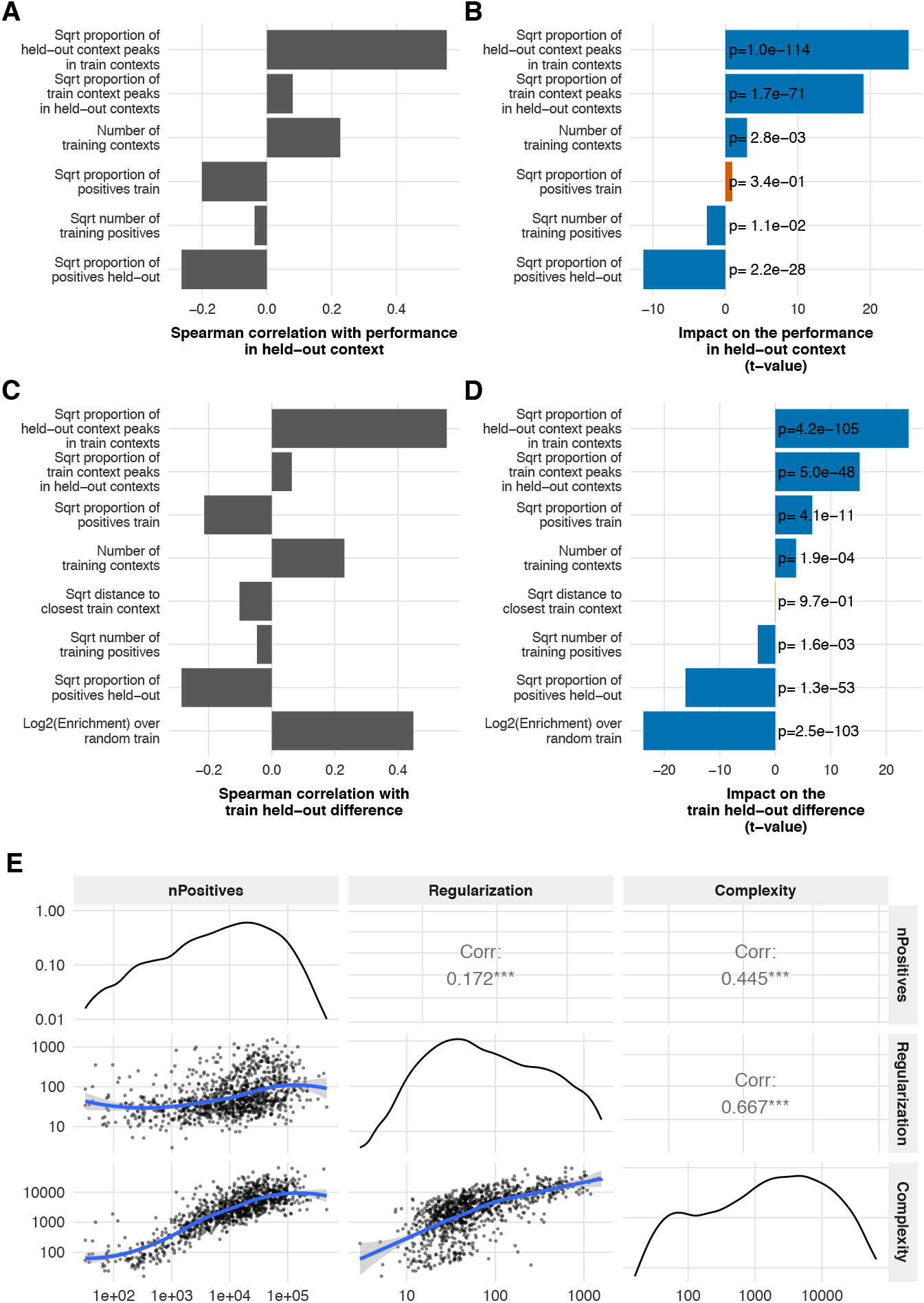
Association between performance and several potential determinants. **A** Spearman correlation between several potential performance determinants and the log2 enrichment over random. Obtained on all held-out TF-cellular context combinations assessed in LOCO rounds during the large scale prediction effort. **B**: Log2 enrichment over random regressed on the latter terms. T-values of the respective coefficients are shown. **C**: Similarly, spearman correlation between the log2 ratio of the enrichment over random in the held-out contexts and the training contexts and (**D**) t-values of the respective regression coefficients. **E**: Model complexity and regularization increase with the number of positives in training.

**Supplementary Fig. 18.**
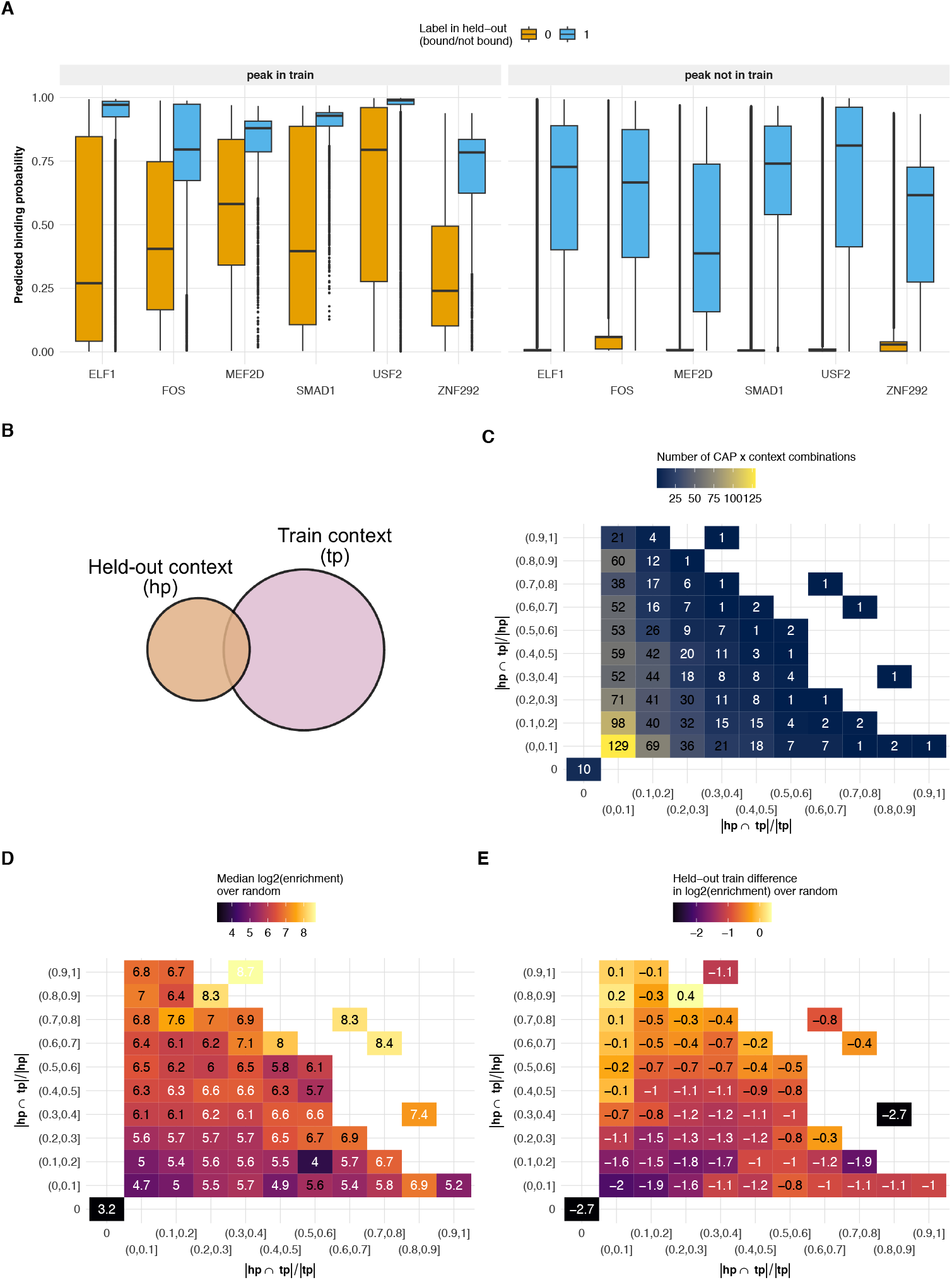
TFBlearner performance in dependence of fraction of peaks observed in training contexts. **A**: Predicted probabilities stratified by whether a site was observed with a peak in any of the training contexts. The color fill corresponds to the label of the respective site in the held-out context. **B**: Overview scheme of the peak sets. **C**: All held-out TF-cellular context combinations of the large scale prediction effort binned by fraction of all training peaks observed in the held-out context (x-axis) and the fraction of all held-out peaks observed during training. Colored by the number of held-out TF-cellular context combinations in the respective bin. **D**: The same bins colored by the median log2 enrichment over random obtained for the respective held-out TF-cellular context combinations and (**E**) colored by the median held-out train difference.

**Supplementary Fig. 19.**
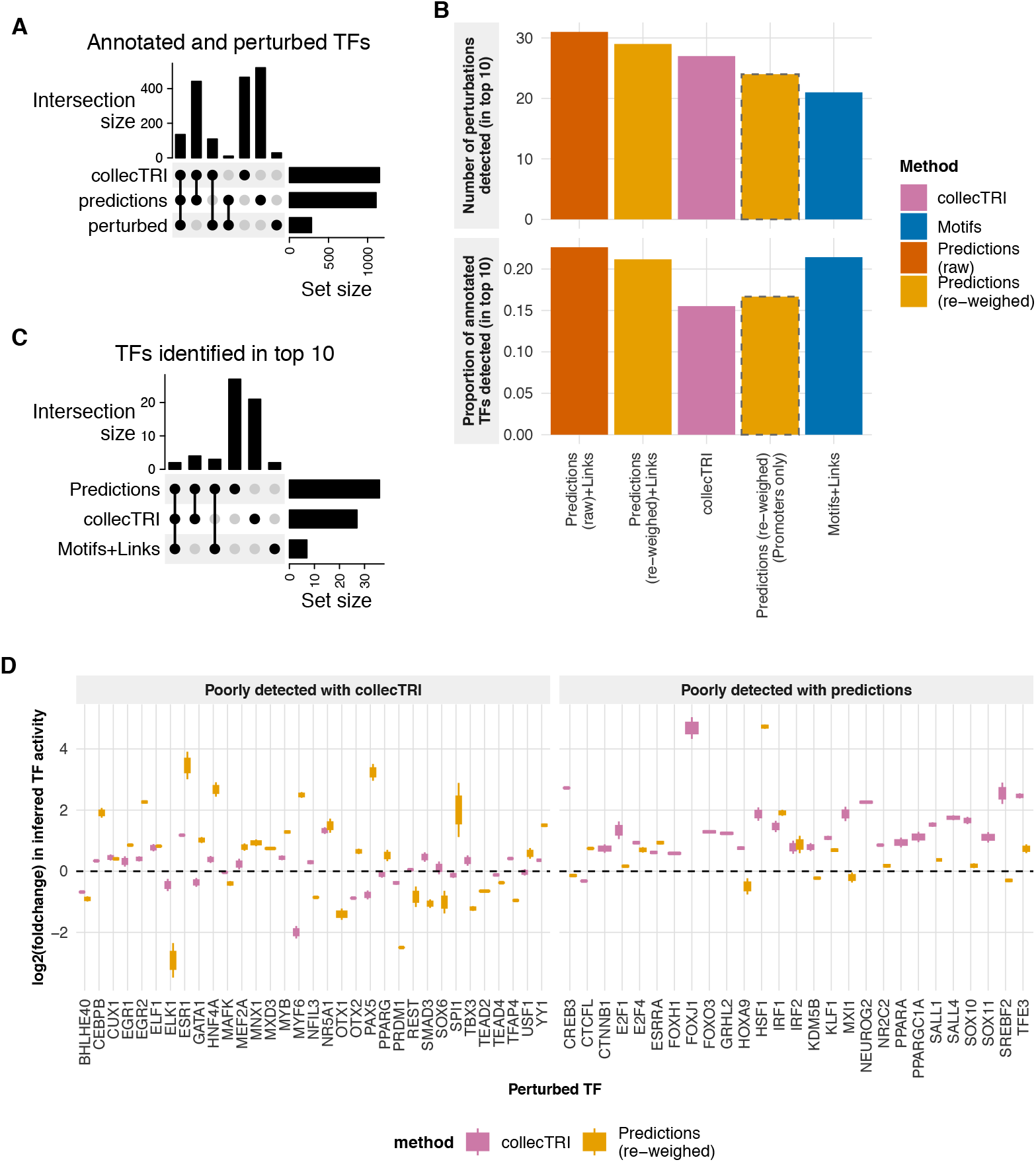
Differential TF activity inference from the transcriptome. **A**: Overlaps of the perturbed TFs with TFs annotated in both resources. **B**: Benchmark of differential TF activity inference from the transcriptome, using log2-foldchanges instead of logCPMs as input. **C**: Overlaps between the perturbed TFs accurately detected by each method, highlighting their complementarity. **D**: Perturbations accurately detected (i.e. in top 10) by only one of the two method, despite being also annotated in the other. Plotted are the inferred changes in activity (of the perturbed TF) in perturbed samples, relative to the average of the controls.

**Supplementary Fig. 20.**
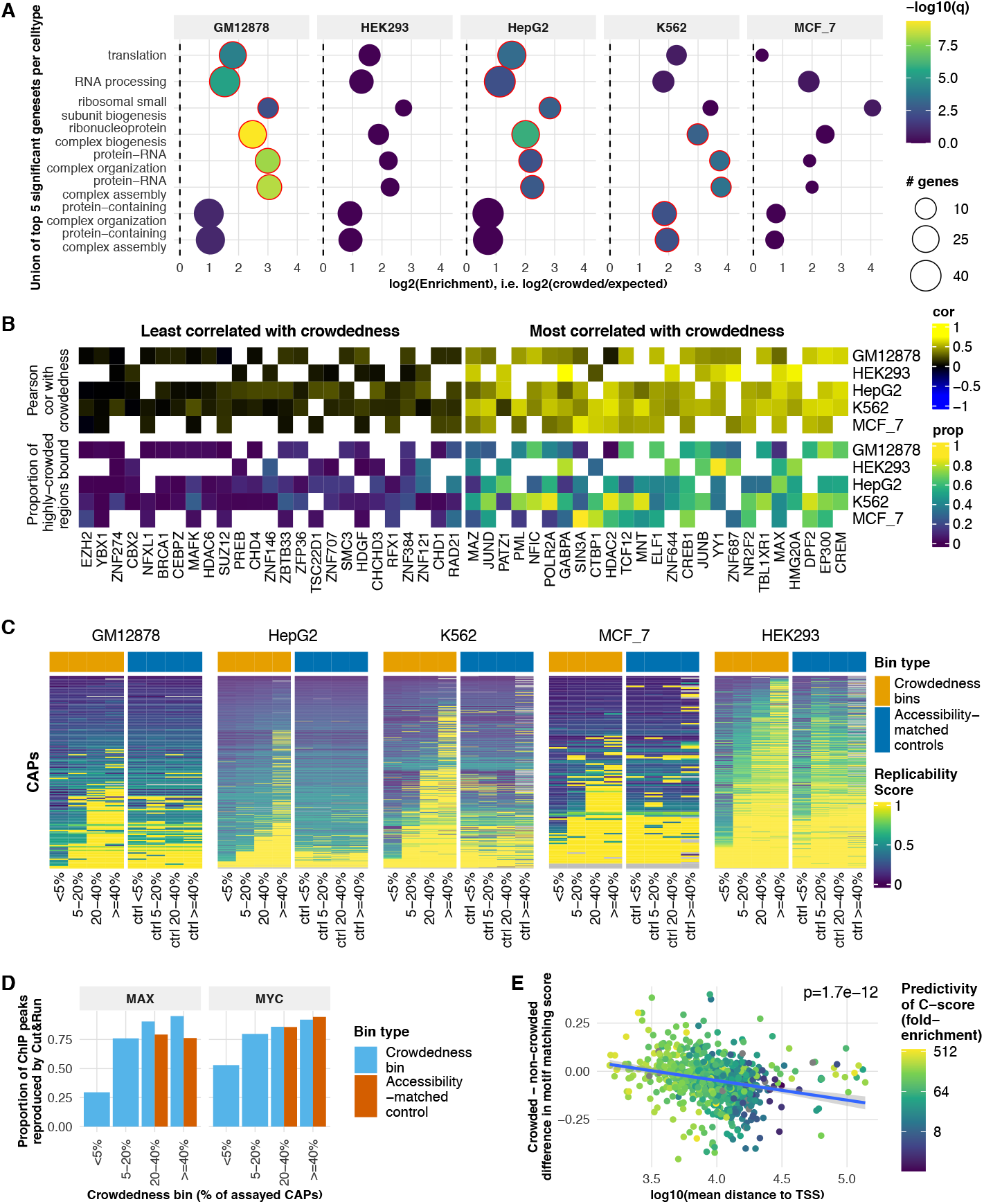
Characterization of highly-crowded regions. **A**: Gene ontology enrichments of highly-crowded (*>*=40%) promoters across cell types, compared to all genes. **B**: TFs most (right) and least (left) associated with crowdedness across cell types. **C**: Median replicability score of peaks within each crowdedness bin or accessibility-matched control bin for each TF for which we have ChIP-seq data. In general, peaks in crowded regions are equally or more reproducible than those in control regions. **D**: Proportion of the ChIP-seq peaks that are reproduced by Cutrun data (i.e. overlap a cut&run peak) for each crowdedness bin (or accessibility-matched control bin). Cut&run peaks were downloaded from GEO (accession IDs GSM3677846 and GSM3677847) and lifted-over to hg38. **E**: TFs that tend to bind closer to TSS tend to be better predicted by the C-score (cell type agnostic crowdedness) and show a stronger motif in crowded versus non-crowded bound places, whereas those binding more distally show weaker motifs in bound crowded regions than in accessibility-matched bound non-crowded regions. Each dot represents one TF, the x-axis represents the mean distance of ChIP peaks to the nearest TSS (averaged across cellular contexts), and the y-axis represents the difference in relative motif matching score (as a proportion of the TF’s maximum matching score) between highly-crowded bound sites and accessibility-matched bound sites. The reported p-value is from the correlation between the x- and y-axes.

